# Tabula Microcebus: A transcriptomic cell atlas of mouse lemur, an emerging primate model organism

**DOI:** 10.1101/2021.12.12.469460

**Authors:** The Tabula Microcebus Consortium, Camille Ezran, Shixuan Liu, Stephen Chang, Jingsi Ming, Olga Botvinnik, Lolita Penland, Alexander Tarashansky, Antoine de Morree, Kyle J. Travaglini, Jia Zhao, Gefei Wang, Kazuteru Hasegawa, Hosu Sin, Rene Sit, Jennifer Okamoto, Rahul Sinha, Yue Zhang, Caitlin J. Karanewsky, Jozeph L. Pendleton, Maurizio Morri, Martine Perret, Fabienne Aujard, Lubert Stryer, Steven Artandi, Margaret Fuller, Irving L. Weissman, Thomas A. Rando, James E. Ferrell, Bo Wang, Iwijn De Vlaminck, Can Yang, Kerriann M. Casey, Megan A. Albertelli, Angela Oliveira Pisco, Jim Karkanias, Norma Neff, Angela Ruohao Wu, Stephen R. Quake, Mark A. Krasnow

## Abstract

Mouse lemurs are the smallest, fastest reproducing, and among the most abundant primates, and an emerging model organism for primate biology, behavior, health and conservation. Although much has been learned about their physiology and their Madagascar ecology and phylogeny, little is known about their cellular and molecular biology. Here we used droplet- and plate-based single cell RNA-sequencing to profile 226,000 cells from 27 mouse lemur organs and tissues opportunistically procured from four donors clinically and histologically characterized. Using computational cell clustering, integration, and expert cell annotation, we defined and biologically organized over 750 mouse lemur molecular cell types and their full gene expression profiles. These include cognates of most classical human cell types, including stem and progenitor cells, and the developmental programs for spermatogenesis, hematopoiesis, and other adult tissues. We also described dozens of previously unidentified or sparsely characterized cell types and subtypes. We globally compared cell type expression profiles to define the molecular relationships of cell types across the body, and explored primate cell and gene expression evolution by comparing mouse lemur cell transcriptomes to those of human, mouse, and macaque. This revealed cell type specific patterns of primate specialization, as well as many cell types and genes for which lemur provides a better human model than mouse. The atlas provides a cellular and molecular foundation for studying this primate model organism, and establishes a general approach for other emerging model organisms.

## INTRODUCTION

Systematic genetic and genomic studies of a handful of diverse organisms over the past half century have transformed our understanding of biology. But many of the most interesting and important aspects of primate biology, behavior, disease, and conservation are absent or poorly modeled in mice or any of the other established genetic model organisms^1–3^. Mouse lemurs (*Microcebus spp*.) are the smallest (∼50 g), fastest reproducing (2 month gestation, 8 month (minimum) generation, 1-4 offspring per pregnancy) and among the most abundant primates (millions to tens of millions)^4^, and an emerging primate model organism^5^. Although much has been learned from laboratory studies of their physiology and aging^6–11^, and from field studies in Madagascar of their ecology, behavior, and phylogeny^12–15^, little is known about their genetics or cellular and molecular biology.

To establish a new genetic model organism, the first step has traditionally been to characterize the wild type and then systematically screen for interesting phenotypes and map the affected genes and underlying mutations or, since the advent of gene targeting, to create targeted mutations and assess their phenotype, as is standard in mouse. Systematic screens are underway for mouse lemur, leveraging their standing genetic diversity and the large pool of naturally-occurring mutations, as in human genetics research. The next step is to create a genetic map or reference genome sequence, which is already available for mouse lemur^16^ and is becoming increasingly affordable, accurate, and complete with powerful new genomic sequencing techniques^17^. With the accompanying development of single cell RNA-sequencing (scRNA-seq) technologies^18^, we reasoned that a reference molecular cell atlas would provide a cellular and molecular foundation that would aid definition and understanding of wild type organism, organ, cell and gene function, enable new types of cellular and molecular analyses, and speed genetic mapping, while providing unprecedented resolution and insights into primate specializations and evolution.

Here we set out to create an extensive transcriptomic cell atlas of the adult mouse lemur *Microcebus murinus*, using a similar strategy to the one we used to construct the Tabula Muris mouse cell atlas^19,20^ and the human lung cell atlas^21^ (Fig. 1). We adapted the strategy to address several challenges for a new model organism. First, because there was no classical histological atlas, little molecular information, and few cell markers, we relied on the extensive knowledge of human and mouse cell markers (Table 1) from the long history of these fields plus the new atlases. Second, unlike classical model organisms but similar to human studies, donors are of different genetic backgrounds, ages, and diseases. Hence we collected extensive clinical data and histopathology on every donor and organ^22^, and, as in the prior Tabula Muris atlases, we procured multiple organs from each donor and processed them in parallel (Fig. 1 a-c, Table 2), thereby controlling for the many technical and biological variables, at least among cell profiles from the same donor. Finally, because this is a primate study, our strategy was opportunistic and designed to maximize information from each donor. To achieve our goal, we brought together experts from diverse fields, including mouse lemur biologists, veterinarians, pathologists, tissue experts, single cell genomics specialists, and computational leaders, to create the Tabula Microcebus Consortium - a team of over 150 collaborating scientists from over 50 laboratories at fifteen institutions worldwide.

**Figure 1.**
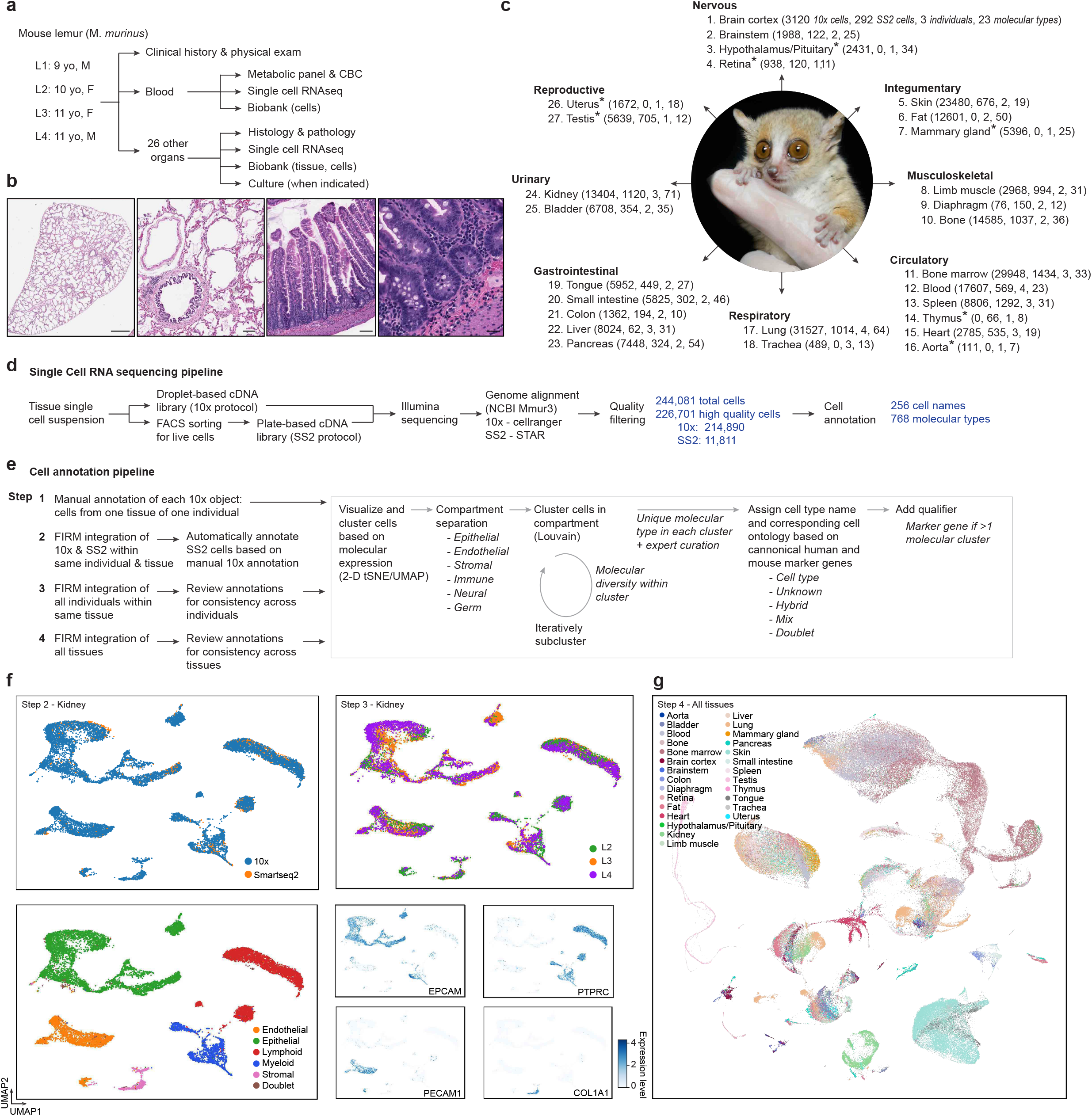
Experimental scheme for constructing the mouse lemur cell atlas. **a**. Overview of four mouse lemurs profiled (name, age, sex), associated metadata collected, and uses of the procured tissues. L1, mouse lemur 1; yo, years old; M, male; F, female; CBC, complete blood count. **b**. Representative tissue histology. Micrographs of hematoxylin-and-eosin-stained (H&E) lung section (left) and close-up (second from left) from L1 and small intestine (second from right) and close-up (right) from L3. Full histological atlas of all tissues analyzed from all individuals is available online at the Tabula Microcebus portal, and identified histopathology is described in Casey et al.^22^. Scale bars (left to right) 1 mm, 100 μm, 100 μm, 25 μm. **c**. The 27 mouse lemur tissues harvested, organized by system, showing for each tissue the number of high-quality cells obtained by 10× scRNA-seq protocol, by SS2 scRNA-seq protocol, number of biological replicates (individuals), and number of molecular cell types identified. *, Tissues which were technically challenging to obtain and for which we only have one biological replicate. **d**. Flow diagram for obtaining and processing single cell RNA-sequencing data. **e**. Flow diagram for clustering cells with related transcriptomic profiles and annotating the molecular cell types. **f**. Representative tissue UMAP showing scRNA-seq profiles of kidney cells (dots) integrated in the same embedded space via FIRM across 10x and SS2 datasets (top left, Step 2) and three individuals (top right, Step 3). Compartment identities of the cell clusters are shown (bottom left) along with heat maps of expression levels (ln(UP10K+1) for 10x data and ln(CP10K+1) for SS2 data, see Methods) of the indicated compartment marker genes (bottom right; *EPCAM*, epithelial; *PTPRC*, immune lymphoid/myeloid; *PECAM*, endothelial; *COL1A1*, stromal). **g**. UMAP of all 244,081 cells in the atlas integrated by FIRM algorithm across all 27 tissues analyzed from four individuals.

## RESULTS

### 1. Expression profiles of over 226,000 cells from 27 mouse lemur organs

The approach used to create a mouse lemur molecular cell atlas is diagrammed in Fig. 1. Two male and two female aged laboratory *Microcebus murinus* mouse lemurs (L1, 9 y/o male; L2, 10 y/o female; L3 11 y/o female; L4, 11 y/o male) were euthanized for humane reasons over a 5 year period due to clinical conditions that failed to respond to therapy, as detailed in a companion manuscript^22^. At the time of euthanasia, blood was drawn, then fresh tissues were quickly isolated (<2 hours post-euthanasia) by a veterinary pathologist and divided into samples that were immediately fixed for pathology or rapidly transported (minutes) at 4°C to the lab where organ-specific experts dissociated each tissue into single cell suspensions for expression profiling using protocols optimized for each organ (Fig. 1a and Supplementary Methods). Full veterinary evaluation, clinical pathology, and histopathological analysis are provided in a companion manuscript^22^ as well as metadata for each individual, organ, and cell profiled in Supplementary Methods. This created a classical histological atlas of the mouse lemur (Fig. 1b, and online at the Tabula Microcebus portal).

From each individual, 3 to 24 organs and tissues were profiled by scRNA-seq, cumulatively totaling 27 different organs and tissues, most of which (20) were profiled in at least two subjects, although seven sex-specific or technically challenging tissues were profiled in only one (Fig. 1c and Table 2). Beyond the 19 tissues profiled in mouse for Tabula Muris^19,20^ (aorta, bladder, bone marrow, brain, diaphragm, fat depot (mesenteric, subcutaneous, brown interscapular, and peri-gonadal), heart, kidney, colon, limb muscle, liver, lung, mammary gland, pancreas, skin, spleen, thymus, tongue, and trachea), we also profiled blood, bone, brainstem, pituitary gland, retina, small intestine, testis, and uterus (Fig. 1c).

RNA sequencing libraries were prepared directly from single cell suspensions of each organ from each individual using the droplet-based 10x Chromium (10x) protocol. For most organs, aliquots of the cell suspensions were also flow sorted for live cells and then RNA sequencing libraries were prepared robotically from individual cells using the plate-based Smart-seq2 (SS2) protocol. The 10x and SS2 libraries were sequenced to achieve saturation. The higher throughput and lower cost of 10x allowed profiling of more cells, whereas SS2 provided greater transcriptomic coverage that aided cell classification, detection of genes expressed at low levels, and gene discovery and structure characterization (accompanying manuscript^23^). Sequencing reads were aligned to the *M. murinus* genome^16^, and aligned reads were counted (as unique molecular identifiers (UMIs) for 10x, and as reads for SS2) to determine the level of expression in each cell for each gene and scaled using Seurat v2^24^. After initial filtering of cell expression profiles for gene count (>500 expressed genes per cell) and sequencing read/UMI parameters (>1000 UMIs for 10x and <5000 reads for SS2, except for heart as described in Methods), and removing cells with profiles compromised by index switching^25^, we obtained 244,081 cell profiles. We then removed low quality cells and putative cell doublets, leaving 226,701 high quality single cell transcriptomic profiles: 214,890 from 10x (16,682 - 88,910 per individual) and 11,811 from SS2 (394 - 6,723 per individual) (Fig. 1 c-d, Table 2).

For cell annotation, the profiles obtained from 10x scRNA-seq analysis of each organ and individual were separately analyzed through dimensionality reduction by principal component analysis (PCA), visualization in 2-dimensional projections with t-Stochastic Neighbor Embedding (t-SNE) and Uniform Manifold Approximation and Projection (UMAP), and clustering by the Louvain method in Seurat v2. For each obtained cluster of cells with similar profiles, the tissue compartment (epithelial, endothelial, stromal, immune, neural, germ) was assigned based on expression of the mouse lemur orthologs of compartment-specific marker genes from mouse and human (Table 1)^21^. Cells from each compartment, organ, and individual were then separately and iteratively clustered until the differentially-expressed genes that distinguished the resultant cell clusters were deemed not biologically relevant (e.g., stress or ribosomal genes) (Fig. 1e). Each resultant cluster was assigned a cell type designation, as detailed below.

We then integrated the SS2 data with the 10x data from the same organ and individual (Fig. 1e-f). For this we used the cross-platform integration algorithm, FIRM^26^, which provided superior integration of shared cell types across platforms while preserving the original structure for each dataset. The cell type designation of each SS2 cell profile was automatically assigned based on the designation of the neighboring 10x cells, and manually curated. We next used FIRM to integrate the combined 10x/SS2 datasets for each organ across the profiled individuals, and then to integrate the profiles across all 27 organs into a single embedded space (Fig. 1g). At each integration step, cell type designations were manually verified to ensure consistency of nomenclature throughout the atlas. Cell profiles that co-clustered across organs were given the same cell type designation, and clusters distinguished only in the merged atlas (e.g., related cells too rare in an individual organ to separate from other clusters in that organ) were re-assigned their own cell type designation. This approach identified 768 molecularly-distinct cell populations (“molecular cell types”) across the 27 profiled organs, with 28±17 (mean±SD) cell populations per organ and 294±1007 cells in each population (Fig. 1c-d, Table 2), which were given 256 different cell designations as detailed below (Fig. 2a).

**Figure 2.**
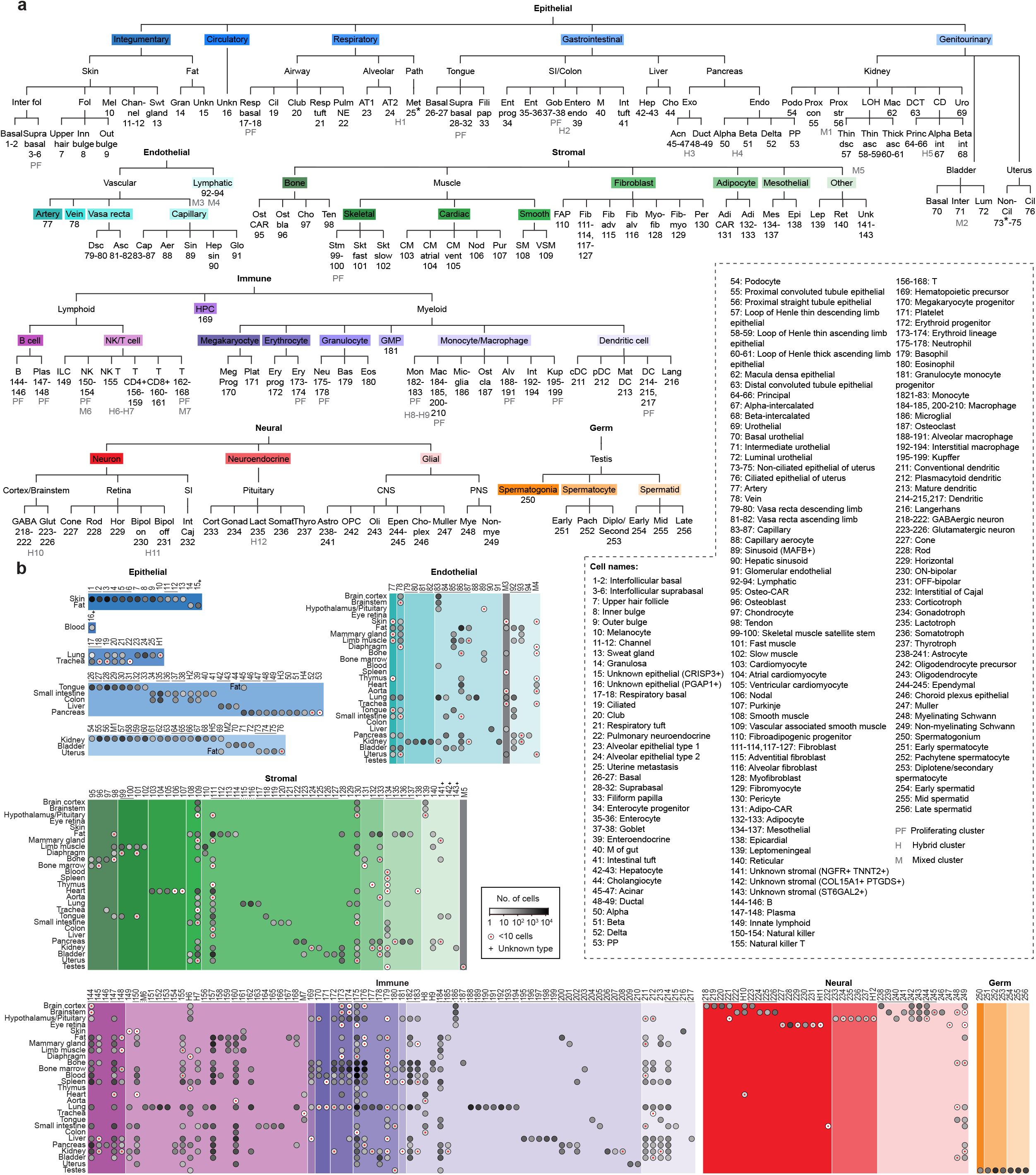
Taxonomy of identified mouse lemur molecular cell types. **a**. Dendrogram of the 256 assigned designations of molecular cell types across cell atlas. Designations are arranged by compartment (epithelial, endothelial, stromal, immune, neural, germ) and then ordered by organ system (epithelial compartment) or biological relatedness (other compartments). Designation number is given below the abbreviation and full names listed in box at right. Some closely related molecular cell types and states are grouped together but shown separately in panel b and Fig. S1, and described further in Section 4. PF, group includes a proliferative cell state; H1-12, hybrid cell types with symbol placed between the two types whose expression signatures the hybrid type shares; M1-7, mixed clusters of distinct cell types too few to assign separately; *, pathological cell states, lung tumor metastasized from uterus (L2) and uterine tumor (L3), as described in accompanying manuscript^23^. **b**. Dot plot showing number of profiled cells (dot intensity shown by heat map scale, small red dot indicates <10 cells) for each of the 768 identified molecular cell types (including 38 hybrid types) plus 24 mixed clusters isolated from the tissues indicated at left. Molecular cell types in each tissue (rows) are arranged (columns) by cell type number/designation and separated by compartment as in panel a. Black bars, closely related molecular types/subtypes/states (see Fig. 4). +, unknown molecular cell type (Section 4).

### 2. Identification of hundreds of mouse lemur cell types and their expression profiles

To assign provisional cell identities and names to the 768 molecular cell types, we first compiled a list from the literature of canonical marker genes for all of the mouse and human cell types in each compartment of the 27 profiled organs, and found the orthologous mouse lemur genes (Table 1 and Methods). We then searched among the cell clusters of each organ compartment for clusters enriched in expression of each set of cell type marker genes and assigned cells in those clusters the name of the corresponding human and/or mouse cell type and their corresponding cell ontology^27^; for cell types with small numbers, we used expert, biologically-guided manual curation. This allowed us to name almost all cell populations, although for many (34) classical cell types there were multiple corresponding cell populations, i.e., different molecular types (Fig. 2 a-b, Fig. S1). This was resolved by adding a suffix to the cell designations recognizing a differentially-expressed gene or gene signature that distinguished them (see Section 4). For six organs, we verified that these manual, expert-curated cell type assignments correlated well with those well curated and validated cell profiles we previously obtained for mouse and human (see Section 6). We identified the differentially-expressed genes that are enriched in each cell type relative to the entire atlas, to other cell types of the same tissue, and to the same compartment of that tissue (Table 3).

Examples of the identified and named cell types for six of the profiled tissues are shown in Fig. 3a and Fig. S2. For example, in limb muscle (Fig. S2 e-f), we identified 31 molecular cell types distributed across the endothelial (7 types), stromal (10 types), and immune (14 types) compartments. Stromal cells of limb muscle included putative tendon cells (*TNMD+, SCX+*) and adipocytes (*ADIPOQ+*). Fatty infiltrates are rarely seen in aged murine skeletal muscle^28^ but are common in aged human muscle^29^, suggesting mouse lemur could model fatty infiltration of muscle during human aging. We also identified vascular smooth muscle cells (*ACTA2+, MYH11+)* and pericytes (*ACTA2+, HIGD1B+*) as well as cells expressing mature myocyte markers (*ACTA1+, MYL1+*), presumably the self-sealed fragments of mature myofibers. Based on expression of troponin isoforms, myofibers were further subdivided into fast (*TNNT3+, TNNI2+, TNNC2+*) and slow (*TNNT1+, TNNI1+, TNNC1+*) myocytes.

**Figure 3.**
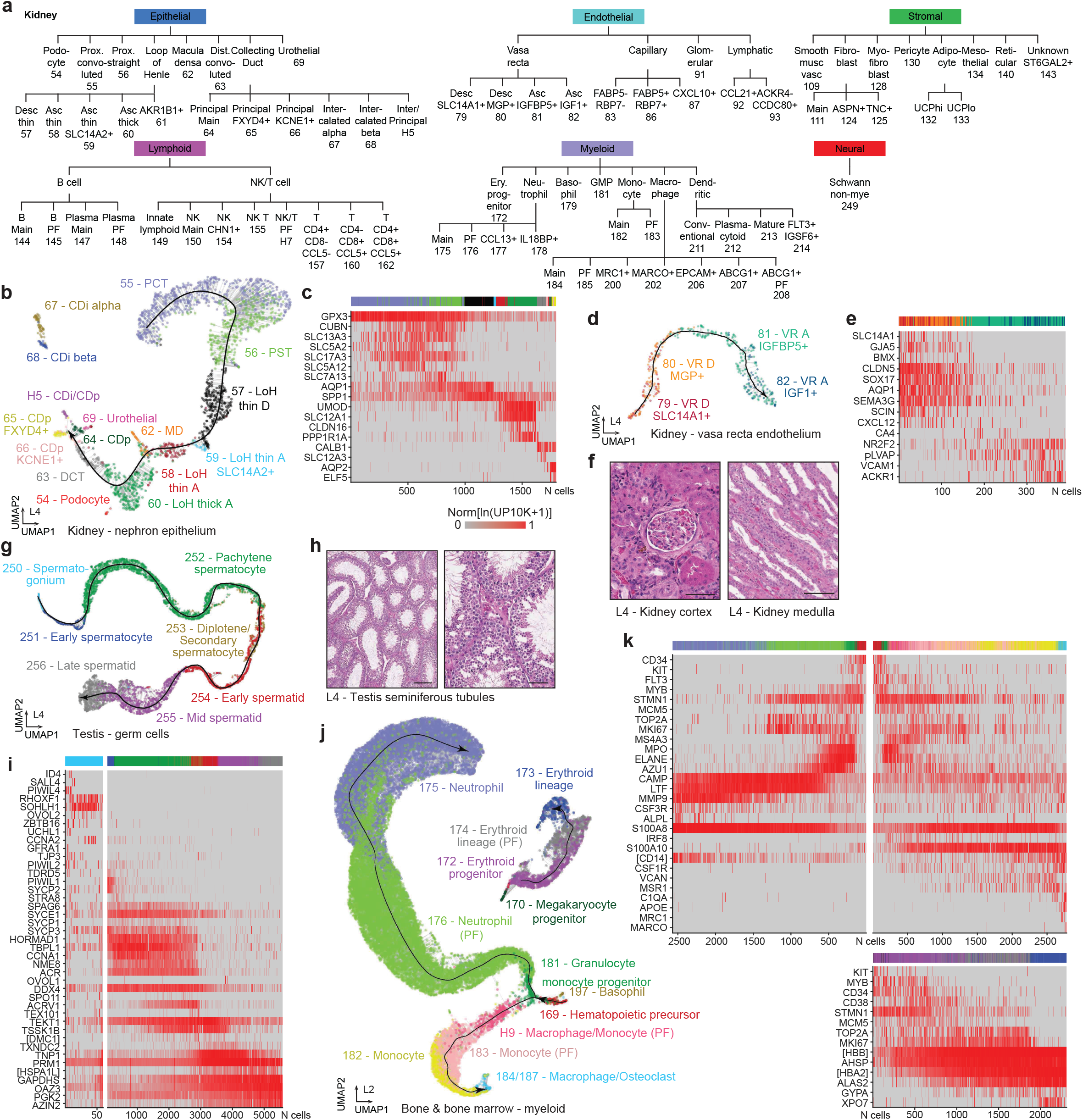
Cell types with gene expression gradients. **a**. Dendrogram of the 71 molecular cell types identified by scRNA-seq of kidney, arranged as in Fig. 2a. **b**. UMAP of kidney epithelial cells (L4, 10x dataset) showing a detected molecular trajectory. Dots, individual cells colored by molecular cell type as indicated; thick black line, cell density ridge (trajectory line); thin gray lines, shortest connecting point of cell to trajectory line. Trajectory shows spatial continuum of molecular cell identities along nephron, beginning with proximal convoluted tubule (PCT, cell type designation #55; see panel a) and ending with principal cells of collecting duct (CDp, #64-66). Macula densa cells (MD, #62) cluster between thin and thick ascending loop of Henle (LoH) cell types (#57-60), and urothelial cells (#69) cluster near CDp cells. Intercalated cells of collecting duct (CDi, #67, 68) and podocytes (#54) cluster separately from trajectory. PCT, proximal convoluted tubule; PST, proximal straight tubule; LoH thin D, loop of Henle thin descending limb; LoH thin A, loop of Henle thin ascending limb; LoH thick A, loop of Henle thick ascending limb; DCT, distal convoluted tubule; CDp, principal cell; CDi alpha, alpha intercalated cell; CDi beta, beta intercalated cell; MD, macula densa. **c**. Heat map of relative expression of indicated canonical nephron cell marker genes in cells along nephron trajectory in panel b. Colored bar at top shows cell type designations (colors as in panel b). Gene expression values (ln(UP10K+1) are normalized to stable maximal value (99.5 percentile) for each gene across all cells in trajectory. **d**. UMAP of kidney vasa recta endothelial cells (L4, 10x dataset) showing another detected molecular trajectory as in panel b. Trajectory shows spatial continuum of molecular cell identities along vasa recta. VR D, vasa recta descending limb; VR A, vasa recta ascending limb. **e**. Heat map of relative expression of indicated canonical marker genes in cells along vasa recta trajectory in panel d. Note transitions along trajectory (left to right) from artery/arteriole (*GJA5+*) to capillary (*CA4+*) to vein (*ACKR1+*) markers. **f**. Micrographs of H&E-stained sections of kidney medulla and cortex analyzed in panels b-e. Scale bars: 100 μm (left), 50 μm (right). **g**. UMAP of germ cells from testis (L4, 10x dataset) showing detected molecular trajectory as in panel b. Trajectory shows developmental continuum (developmental pseudotime) of molecular cell identities during spermatogenesis, beginning with stem cells (spermatogonium, #250) and progressing to late spermatids (#256). See Fig. 6c for dot plot of expression of canonical marker genes along trajectory. **h**. Micrographs of H&E-stained sections of testis seminiferous tubules analyzed in panel g. Scale bars: 200 μm (left), 50 μm (right). **i**. Heat map of relative expression of indicated canonical marker genes in cells along the spermatogenesis trajectory in panel g. The spermatogonium panel is enlarged for better visualization. [], description of gene identified by NCBI as a gene locus (LOC105858542 [DMC1], LOC105858168 [HSPA1L]). j. UMAP of myeloid cells from bone and bone marrow (L2, 10x dataset) showing two detected molecular trajectories. One (at left) shows developmental continuum (developmental pseudotime) of molecular cell identities beginning with hematopoietic precursor cells (HPC, #169) and bifurcates at granulocyte-monocyte progenitor cells (GMP, #181) into neutrophil lineage (#175-176) and monocyte/macrophage lineage (#182-184,187). Another trajectory (at right) connects the erythroid progenitor and lineage cells (#172-174), with megakaryocyte progenitor cells (#170) nearby. **k**. Heat map of relative expression of indicated canonical marker genes in cells (uniformly subsampled for neutrophils) along developmental pseudotime trajectories in panel i (top left, neutrophil trajectory; top right, monocyte/macrophage trajectory; bottom: erythroid trajectory). LOC105862649 [CD14], LOC105883507 [HBB], LOC105856254 [HBA2]. See also Fig. S3 for similar trajectories in additional individuals and heat maps of genes differentially-expressed along each of the six trajectories.

We identified two putative adult stem/progenitor cell populations in limb muscle (Fig. S2 e-f). One was a cell cluster in the myocyte/myogenic compartment that expressed the human and mouse muscle stem cell (MuSC, or “satellite cell”) transcription factors *MYF5* and *PAX7*. These cells also expressed *CD56* (*NCAM1*) and *CD82*, cell surface markers used to purify human MuSCs^30,31^, as well as *VCAM1* or *ITGA7* used to purify mouse MuSCs^32,33^. A similar population of putative MuSCs (*CD56+, CD82+, VCAM1+, ITGA7+*) was found in the diaphragm. We also identified putative fibroadipogenic progenitors (FAPs), which in humans and mice give rise to fibroblasts and adipocytes (and perhaps chondrocytes and osteoblasts) and promote MuSC-mediated regeneration and sustain the MuSC pool^34–36^. The putative lemur FAPs were found in both limb muscle and diaphragm as stromal populations that selectively expressed *PDGFRA* and *THY1*, surface markers used to purify human FAPs^37^ (note that *LY6A* (*SCA-1*), used to purify mouse FAPs^34^, is not annotated in the lemur genome). In a companion paper (de Morree et al.), we use the identified surface markers to purify and functionally characterize these putative stem/progenitor populations from lemur, and show they exhibit many characteristics more similar to their human counterparts than those of mouse.

From blood, we identified mouse lemur cognates of all the major human and mouse immune cell types including, in the lymphoid lineage, B cells, plasma cells, *CD4+* T cells, *CD8+* T cells, natural killer cells, natural killer T cells, and innate lymphoid cells; and in the myeloid lineage, erythrocytes, platelets, monocytes, macrophages, conventional and plasmacytoid dendritic cells, neutrophils, basophils, and even the rare and fragile eosinophils (Fig. 2). In the bone, bone marrow and other hematopoietic and lymphoid tissues, we identified presumptive progenitors including hematopoietic precursors, and progenitors of erythrocytes, megakaryocytes and granulocyte-monocytes and putative adipogenic and osteogenic progenitors. However, certain immune cell subtypes in human and/or mouse were not identified in lemur. For example, despite large numbers (>9000) captured in multiple tissues, lemur monocytes formed a single cluster in most tissues that could not be resolved into classical and non-classical monocytes based on the markers used to distinguish the two cell types in human (*CD14, CD16*) or mouse (*CCR2, CX3CR1*, and *LY6C1/2* which has no primate ortholog)^38^. These marker genes were either not annotated in the lemur genome or not differentially expressed (see Table 1 and accompanying manuscript^23^), suggesting either limitations due to current genome annotation or unique lemur biology. Similarly, conventional dendritic cells (cDCs) detected in the lemur analysis could not be divided into type 1 and type 2 subtypes characteristic of humans and mice. Conversely, other dendritic molecular types were identified in lemur (e.g., *FLT3+ IGSF6+* DCs) that had no apparent human or mouse cognates. The full spectrum of developing and mature immune cells across the body allowed us to reconstruct the early stages of hematopoietic development (Section 3) and its evolution (Section 6), as well as the subsequent dispersal of mouse lemur immune cells throughout the body and its alteration by disease (accompanying manuscript^23^).

### 3. Molecular gradients of cell identity

Although the vast majority of profiled lemur cells could be computationally separated into discrete clusters of cells with similar expression profiles, we found many examples where cells instead formed a continuous gradient of gene expression profiles indicating a gradual transition from one molecular identity to another within the tissue. Some of these reflect a spatial gradient of cell identity in the tissue, whereas others correspond to an ongoing developmental process or the induction of a physiological or pathological cell state.

The kidney provided a dramatic example of spatial gradients of cell types. Coordinated gradients of renal epithelial and endothelial cell types correspond to the spatial organization of the nephron that controls fluid and electrolyte balance. Among the ∼14,800 cells profiled from lemur kidneys, we identified lemur cognates of almost every important cell type known from human and mouse in each of the major tissue compartments (Fig 3a-f). Most notable among the profiled kidney cells were the many epithelial cells that formed a long continuous gradient of molecular identity (Fig. 3b-c, Fig. S3a, g, m-n). With canonical renal markers, we determined that this gradient corresponds to the spatial gradient of epithelial cell types along the mouse lemur nephron, starting from proximal convoluted tubule cells, through the loop of Henle, and ending with principal cells of the collecting duct. Interestingly, macula densa cells, the sodium-sensing cells that regulate glomerular blood flow and filtration rate and that are typically physically located at the distal end of the loop of Henle before the start of the distal convoluted tubule^39^, surprisingly localized in the lemur molecular cell gradient between cells of the thin and thick loop of Henle. We also found a prominent cell gradient in the endothelial compartment, with arterial markers (*GJA5+, BMX+*) expressed at one end and venous markers (*ACKR1+, VCAM1+*) at the other, and capillary markers (*CA4+*) in between (Fig. 3d-e, Fig. Sxc-d, Fig. S3b, h, o-p). This appears to comprise the vasa recta, a network of arterioles and venules intermingled with the loop of Henle in the renal medulla, because it expressed specific vasa recta markers (e.g., *AQP1, SLC14A1* for vasa recta descending arterioles) and was molecularly distinct from the clusters of glomerular endothelial cells (*EDH3+*), other capillary endothelial cells (possibly peritubular), and lymphatic endothelial cells (*CCL21+*). This deep molecular map of the lemur nephron also revealed region-specific hormonal regulation of nephron function (see accompanying hormone atlas manuscript^40^).

Several other gradients of gene expression profiles represent cells differentiating in stem cell lineages. Two such gradients in bone marrow represent the ongoing development and maturation of hematopoietic progenitors (Fig. 3j-k, Fig. S3d-f, j-l, q-r). One gradient bifurcates into the monocyte/macrophage and the granulocyte/neutrophil lineages (accompanying manuscript^23^), whereas the other represents the erythroid lineage. Some common but more subtle gradients marked differentiation of basal epithelial cells in the skin and at least four other organs (small intestine, colon, tongue, bladder) into their corresponding mature epithelial cell types along the suprabasal/luminal axis of each organ.

The most striking developmental gradient was in the male gonad. Among the ∼6500 testes cells profiled, we found on computational clustering that all germ cells formed a single continuous gradient, which the expression profiles indicated was a continuum of germline cells progressing through spermatogenesis (Fig. 3g-i, Fig. S3c, i). The gradually changing expression levels of genes across the continuum allowed us to reconstruct the full gene expression program of mouse lemur spermatogenesis, assigning seven canonical stages from stem cells (spermatogonia) to mature spermatids using orthologs of stage-specific markers from human and mouse (Fig. 3i). In addition to the essential role of male germ cell differentiation in reproduction, the lemur spermatogenesis program is of special interest because such programs are rapidly evolving during primate speciation (Section 6), with several notable evolutionary specializations already recognized for mouse lemur, including seasonal regulation and sperm competition^41^.

### 4. Previously unknown or uncharacterized cell types and subtypes

Although we were able to assign a provisional identity to the vast majority of cell populations based on expression of orthologs of canonical markers of human and mouse cell types, there were dozens of cases in which more than one cluster in a tissue expressed markers of the same cell type and their separation could not be attributed to technical differences (e.g., cell quality, batch effect, cell doublets) (Fig. 4a). In some cases these appear to be multiple states of the same cell type because the differentially-expressed genes that distinguished the clusters included proliferation markers (e.g., *MKI67, TOP2A)* indicating a proliferative state (hence PF was added to cell type name) or differentiation markers indicating a differentiating state. In most cases, however, the additional clusters appear to represent previously unknown or little characterized cell types or subtypes, and a distinguishing gene or genes was added to the name of the minor cluster (e.g. B cell (*SOX5+*) in pancreas), or to both clusters if they were similar in abundance. Such clusters were uncovered in all compartments except the germline and in most profiled organs and tissues (Fig. 4a, Fig. 2b).

**Figure 4.**
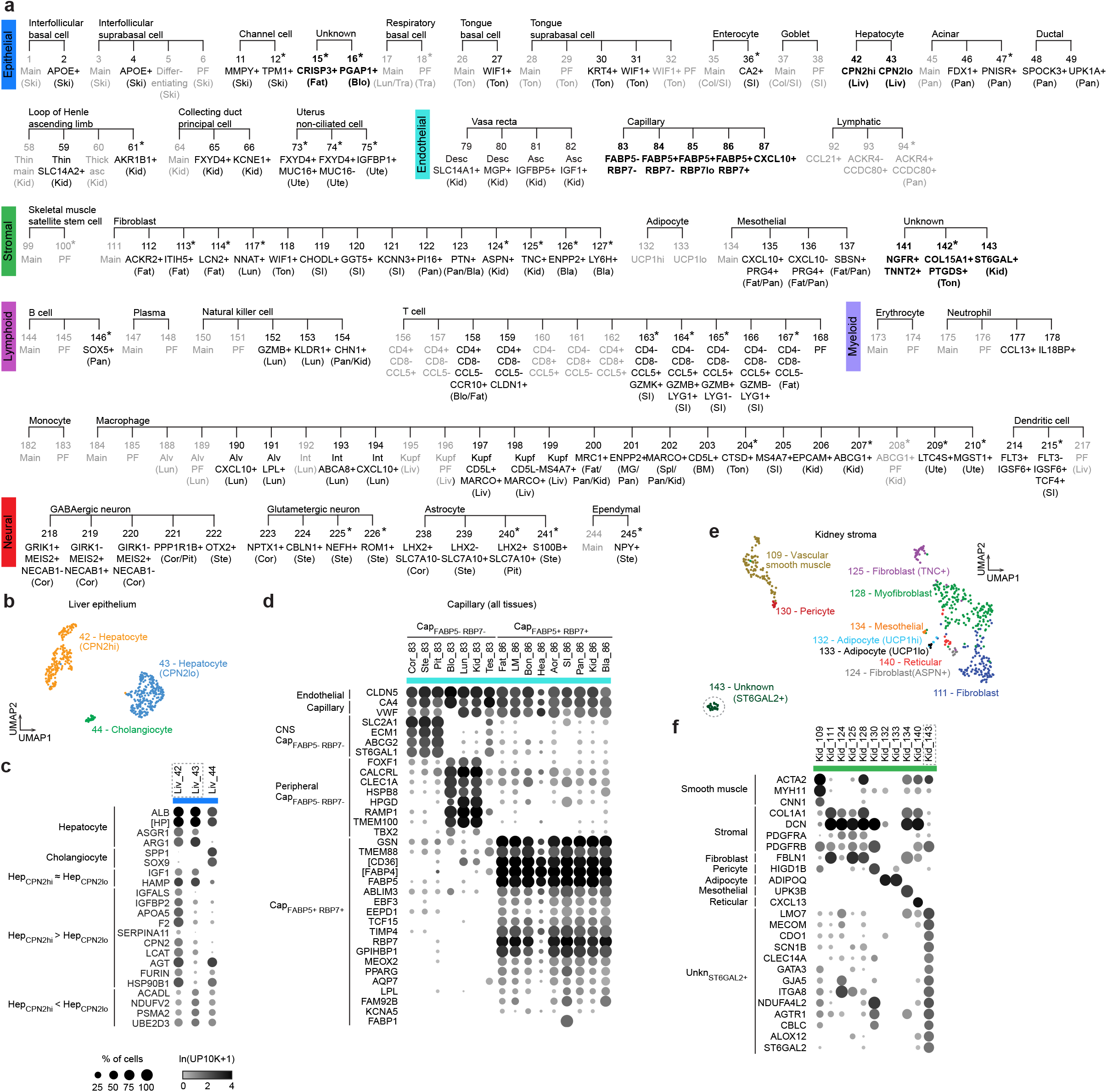
Previously unknown and understudied molecular cell types. **a**. Expansion of molecular cell type dendrogram in Fig. 2a showing each of the molecular cell types that are grouped in Fig. 2a. These include molecular types related to a known cell type but distinguished from the predominant population (“main”) or each other indicated by a distinguishing marker gene (e.g., B cell (*SOX5+*)) or by a proliferative gene signature (PF), and unidentified (“unknown”) molecular cell types. Known molecular types and proliferating states in gray, novel molecular types in black with those highlighted in the text in bold. *, molecular types found in only one individual. (), tissue abbreviation indicated for molecular types found in ≤3 tissues. Bla, Bladder; Blo, Blood; Bon, Bone; BM, Bone marrow; Col, Colon; Cor, Brain cortex; Fat, Fat; Pit, Pituitary; Kid, Kidney; Liv, Liver; Lun, Lung; MG, Mammary gland; Pan, Pancreas; Ski, Skin; SI, Small intestine; Spl, Spleen; Ste, Brainstem; Ton, Tongue; Tra, Trachea; Ute, Uterus. **b**. UMAP of liver hepatocytes and cholangiocytes from three individuals (L2-4, 10x dataset) integrated by FIRM. See also Fig. S4. **c**. Dot plot of average expression (ln(UMI_*g*_/UMI_total_ *1e4 +1), abbreviated ln(UP10K+1)) and percent of cells (circle size) expressing indicated cell type markers and other differentially-expressed genes in the two hepatocyte subtypes and cholangiocytes of panel b. LOC105859005 [HP]. **d**. Dot plot of average expression of indicated endothelial and capillary markers and differentially-expressed genes for all *FABP5+ RBP7+* and *FABP5-RBP7-*capillary molecular types in atlas (L1-4, 10x dataset). Note the two sets of differentially-expressed genes (CNS, Peripheral) that distinguish *FABP5-RBP7-*capillary cells from CNS (Cortex (Cor), Brainstem (Ste), Pituitary (Pit)) and peripheral tissues (Blood (Blo), Lung (Lun), Kidney (Kid)). LOC105857591 [FABP4-like], LOC105879342 [CD36]. **e**. UMAP of kidney stromal cells (L2-4; 10x dataset) integrated by FIRM. Dashed circle, unknown molecular cell type (#143, *ST6GAL2+*; potentially mesangial cells). **f**. Dot plot of average expression of indicated cell type markers and differentially-expressed genes for the kidney stromal cells indicated in panel e. Note unknown molecular cell type (#143) and the set of differentially-expressed genes (labeled Unkn_ST6GAL2+_) that distinguish it from the other molecular types. See also Fig. S5 for additional examples and information of unknown/understudied molecular cell types.

Fibroblast subtypes were particularly diverse (Fig. 4a), with multiple molecular types identified in many organs, most appear organ-specific with little co-clustering across organs and without known parallels in human or mouse. Similarly, macrophages formed multiple distinct molecular clusters in many tissues (Fig. 4a, and accompanying manuscript^23^), and there was diversity among T and natural killer cells that could not be readily harmonized with classical T cell subtypes defined in humans and mice^42,43^.

We also identified interesting molecular subtypes of epithelial cells, including pancreatic acinar and ductal types, kidney collecting duct principal cells, intestinal enterocytes, and hepatocytes (Fig. 4a). These subtypes may serve specialized functions. For example, the *CPN2*hi hepatocytes expressed higher levels of many classical liver secreted proteins (e.g., *APOA5, LCAT, CPN2, AGT*) and their processing enzymes (e.g., *FURIN*) as well as additional hormones and receptors^40^; in contrast, *CPN2*lo hepatocytes expressed higher levels of several mitochondrial metabolic genes (e.g. *ACADL, NDUFV2*) and genes involved in proteasome-mediated protein degradation (e.g., *UBE2D3, PSMA2*) (Fig. 4 b-c). The two types do not correspond to the known zonal heterogeneity of human and mouse hepatocytes^44^, but notably the *CPN2*hi hepatocytes expressed more transcripts and genes than *CPN2*lo hepatocytes (Fig. S4c) so the *CPN2*hi cells could correspond to the larger, polyploid hepatocytes and *CPN2*lo cells to the smaller, diploid hepatocytes^45^. We uncovered a similar molecular distinction among hepatocytes in mouse^20^ and human^46^ liver scRNA-seq datasets (Fig. S4), suggesting these are evolutionarily conserved hepatocyte subtypes.

In the endothelial compartment, there was substantial diversity among blood capillary cells including subtypes found across multiple tissues (Fig. 2b). In some tissues endothelial cells were comprised exclusively of one of these types (e.g. heart, *FABP5+ RBP7+*; brain cortex, *FABP5-RBP7-*), but other tissues contained a mix of two or more (Fig. 2b, Fig. 4a). *FABP5+ RBP7+* capillary cells appear to be specialized for energy storage because they were found in high-energy demand tissues (e.g., heart, limb muscle, kidney) and enriched in genes for fatty acid uptake and binding (e.g., *RBP7, FABP1, FABP5*), as well as for transcription factors *MEOX2* and *TCF15* (Fig. 4d), all genes that are also enriched in capillary cells of several high energy-demand tissues in human and mouse^47–50^. *CXCL10*+ capillary cells express genes associated with interferon activation (e.g., *CXCL9, CXCL10, CX3CLl, GBP1, IFIT3*)^51^ suggesting an inflammatory state in several organs of L2 and the lung of L4 (accompanying manuscript^23^). We also identified molecular subtypes of lymphatic endothelial cells (*CCL21+ CCDC80+* vs. *CCDC80+ ACKR4+*) (Fig. 4a), perhaps representing different lymphatic cell types in peripheral vessels and lining lymph node sinuses^52–54^ (also see accompanying manuscript^23^).

For eight of the 768 molecular cell types (∼1%) we were unable to assign a specific identity, so they were designated “unknown” along with the organ they were isolated from and the compartment inferred from expression of canonical compartment markers. These include stromal types in tongue (cell type designation #142) and kidney (#143), and epithelial types in fat (#15) and blood (#16) (Fig. 4a, e-f, Fig. S5c-m). The remaining four stromal types, from bone, mammary gland, pancreas, and tongue, were given the same designation (#141, “unknown stromal *NGFR+ TNNT2+*”) because they shared similar transcriptomic profiles (including high expression of *NGFR, IGFBP6, OGN, ALDH3A1, CLDN1, ITGB4*) and curiously also expressed high levels of cardiac troponin T (*TNNT2*), an actomyosin contractile component and sensitive and specific cardiac myocyte marker and clinical marker of myocardial infarction^55–57^ (Fig. S5a-e). Some of the unknown cell types (#15, #141, #142) resemble mesothelial cells because many of their differentially-expressed genes were enriched in mesothelial cells (Fig. S5e, j). However, some of the differentially-expressed genes were also characteristic of other cell types (e.g. leptomeningeal and Schwann cells for #141 and 142, urothelial cells for #15), and in the global comparison of cell types (Section 5) they did not closely localize with any of these cell types. The unknown kidney stromal type (#143) might be mesangial cells because they expressed at high levels two of the recently reported genes (*LMO7* and *ITGA8*) enriched in mouse mesangial cells^58^ (Fig. 4e-f, Fig. S5f-g). The most perplexing unknown was the epithelial (*EPCAM+*) population from blood (#16) of one individual (L2). It had a unique gene signature including *PGAP1, PTPRG, FGF13, SOX2, CPNE6, CDH3*, many of which are also expressed by brain ependymal cells, astrocytes, and oligodendrocytes; however, this population did not express canonical markers of those cell types (Fig. S5k-m).

Some of the molecular subtypes and unknown cell types described above may represent previously unrecognized or sparsely characterized cell states including pathological states (accompanying manuscript^23^). Others may be new cell types or subtypes, like the novel lung aerocyte and ionocyte populations uncovered by scRNA-seq^59–61^. It will be important to characterize each of these potentially novel molecular cell types and determine which are conserved across species but previously unrecognized, and which are mouse lemur or primate innovations.

### 5. Global molecular comparison of cell types across the body

We used two approaches to examine the molecular relationships of the over 750 lemur molecular cell types we identified across the organism. One approach was simplified UMAPs in which each molecular type was condensed into a single data point representing the average expression value of all cells of that type (“pseudo-bulk” expression profile) (Fig. 5). The other was to calculate pairwise correlation coefficients of all cell type “pseudo-bulk” expression profiles and display them in a large (∼750x750) matrix (Fig. S6a-b, Fig. S7). Both approaches revealed global patterns of similarity as well as unexpected molecular convergence of seemingly distantly-related cell types.

**Figure 5.**
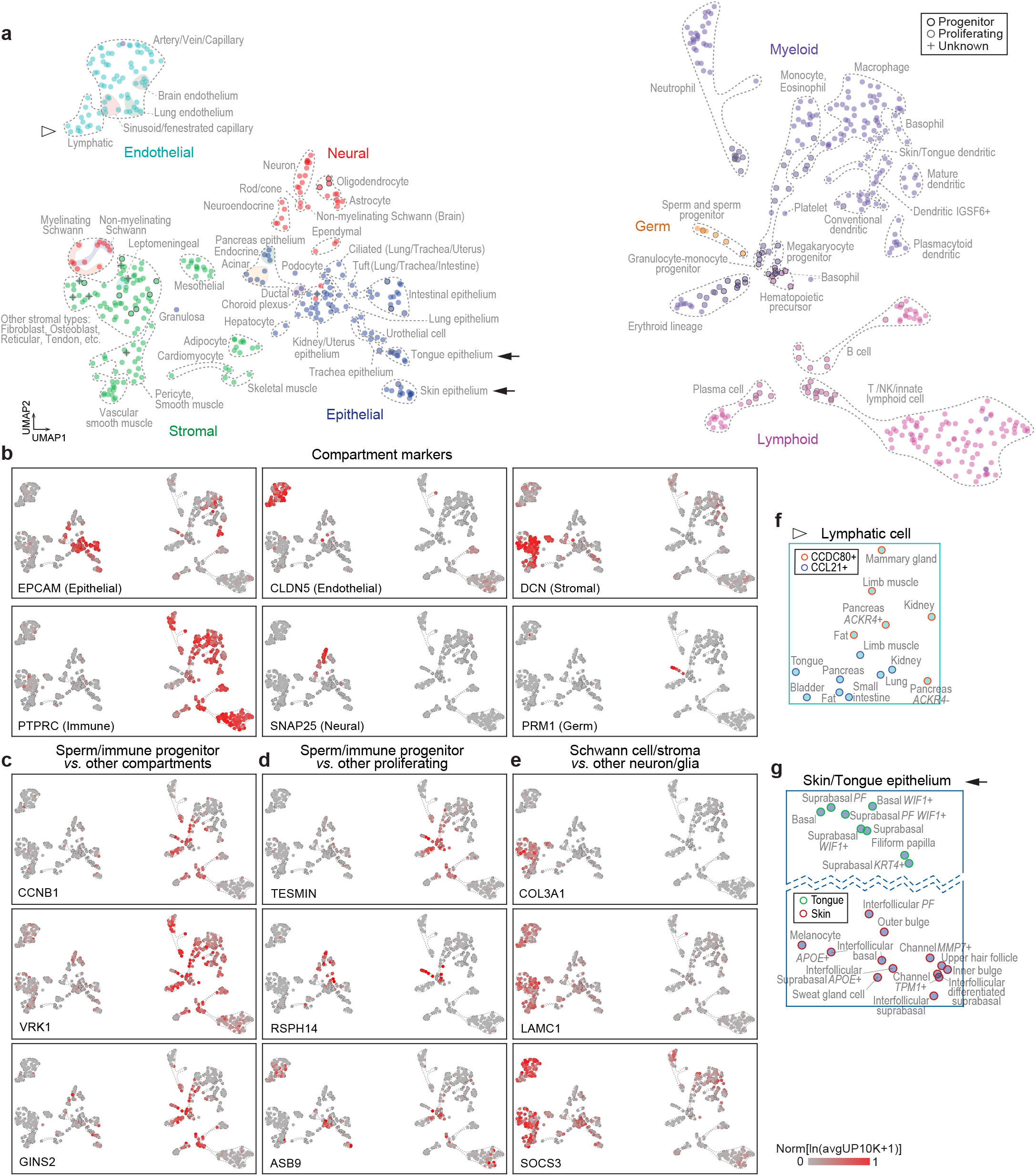
Relationships of molecular cell types across the mouse lemur cell atlas. **a**. UMAP of molecular cell types (dots) in the atlas (L1-L4, 10x dataset) based on their mean transcriptomic profiles. Dot fill color indicates tissue compartment; black outline, progenitor cell types; gray outline, proliferating cell types; +, unknown cell types. Dashed gray contours, groups of the related cell types indicated. Open triangle indicates lymphatic endothelial cell types magnified in panel f. Arrows indicate tongue and skin magnified in panel g. See also Fig. S6-7 for cell type pairwise similarity viewed in a heat map, and Fig. S8 for cell type UMAPs compared across human, lemur, and mouse. **b-e**. The cell type UMAP as in panel a overlayed with relative expression level of compartment (e.g., epithelial, endothelial, stromal) markers (b) or the indicated differentially-expressed genes (c-e) as heat maps. Cell type mean expression values are normalized by the maximally expressed cell type. The differentially expressed genes are those with expression pattern specific to immune progenitor/proliferating cells and germ cells vs. cell types in other compartments (c), or vs. proliferating cells in other compartment (d), and genes with expression pattern specific to Schwann and stromal cells vs. non-Schwann cell types in the neural compartment (e). **f-g**. Close up of portions of UMAP in panel a showing segregation of the two types of lymphatic cells indicated (f), and segregation of skin and tongue epithelial cell types including those with similar locations/functions in the two tissues (e.g., basal cells) (g).

Molecular cell types within a tissue compartment generally showed more similar expression profiles, even those from different organs (Fig. 5a-b, Fig. S6a). Such within-compartment similarity appears general as it was also observed for human and mouse (Fig. S8a-b). Endothelial cell types across the body formed the most coherent compartment. Next was the neural compartment including CNS glial cells, which are surprisingly similar to neurons. Immune compartment cell types were by far the most divergent, and particularly so between lymphoid and myeloid populations. But the global analyses also identified a few exceptions: cell types that were more closely related to cell types in another compartment than to those in its own compartment. Some of these cross-compartment similarities were predictable. For example, neuroepithelial cells of the airway (neuroendocrine cells) and gut (enteroendocrine cells) were found to be more closely related to neurons and pituitary neuroendocrine cells than to most other epithelial cell types.

Other identified cross-compartment similarities were surprising. The most striking was that male germ cells (spermatogonia) are more closely related to immune progenitor cells than to any other cells in the atlas, including progenitors and proliferating cells of other compartments (Fig. 5a, Fig. 6a-b). Maturing male germ cells formed a developmental trajectory emanating out from the immune progenitor cluster, just like the various immune cell lineages (Fig. 5a). Similar convergence of spermatogonia and immune progenitors was observed for human and mouse (Fig. S8a-c). Differential gene expression analysis showed that this similarity included shared enrichment in both spermatogonia and hematopoietic progenitors of specific cell cycle genes, particularly genes involved in M phase (e.g., *CCNB1, VRK1, GINS2*) (Fig. 5c, Fig. S8e), indicating similarity in the mitotic machinery of proliferating spermatogonia and hematopoietic progenitors. Compared to progenitors and proliferating cells of other compartments, spermatogonia and immune progenitors also shared selective expression of non-cell-cycle genes (e.g., *TESMIN, RSPH14, ASB9, ARHGAP15, HEMGN*) (Fig. 5d), many of which were also expressed in human and mouse germ and immune cells/progenitors (Fig. S8e). This suggests a common and potentially conserved role in regulating stemness of immune and sperm progenitors.

**Figure 6.**
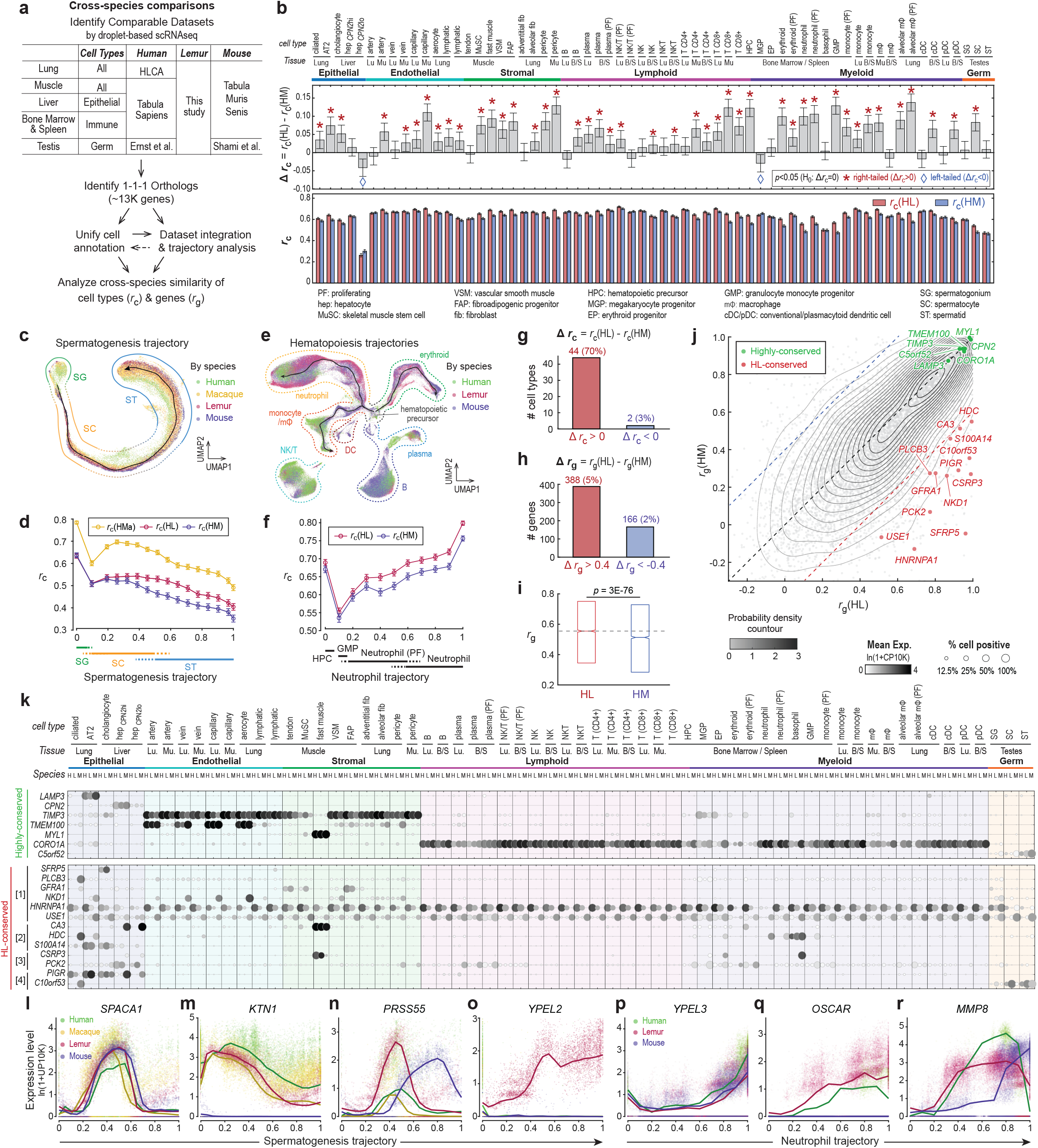
Evolutionary comparison of cell type and gene expression patterns across species. **a**. Overview of the evolutionary comparison, datasets used, and major steps. **b**. Bar plots showing human-to-lemur and human-to-mouse correlation coefficients (*r*_c_, top) and their difference (Δ*r*_c_, bottom) for each of the 63 orthologous cell types across human, lemur, and mouse. Cell types are ordered first by compartments then by tissue. *p*-value calculated using one-tailed *t* test (see Methods). **c**. Species-integrated UMAPs of the male germ cells with cells color coded by spermatogenesis species. Black arrowed curve shows the detected spermatogenesis trajectory. Colored curves indicate spermatogenesis stages with annotations abbreviated as in panel b. See also Fig. S10a-b. **d**. Correlation coefficients of human germline cell transcriptomic profiles to those of lemur (*r*_c_(HL)), macaque (*r*_c_(HMa)), and mouse (*r*_c_(HM)) across different stages of the spermatogenesis trajectory, based on the one-to-one orthologs across the four species. Note that human-macaque similarity is potentially overestimated here because the human and macaque data were from the same study using drop-seq methods^62^, whereas the lemur and mouse data were from two independent studies (this study and Ernst et al.^63^) using 10x scRNAseq. **e**. Species-integrated UMAPs of the bone marrow and spleen immune cells with cells color coded by species. Black arrowed curves show the three detected hematopoiesis branches. Colored curves indicate hematopoietic lineages with annotations abbreviated as in panel b. See also Fig. S10f-i. **f**. Correlation coefficients of the transcriptomic profiles of the human immune cell and progenitors to those of lemur (*r*_c_(HL)) and mouse (*r*_c_(HM)) across different stages of the neutrophil trajectory, based on the one-to-one orthologs across the three species. See also Fig. S10j-k for comparison along the monocyte/macrophage and erythroid trajectories. **g-h**. Bar plots showing the number of cell types (g) or genes (h) with a highly human/lemur-conserved and mouse divergent transcriptional profile (red) vs. the opposite (blue, highly human/mouse-conserved and lemur divergent). **i-j**. Comparison of gene expression conservation patterns between human and lemur (*r*_g_(HL)) vs. that between human and mouse (*r*_g_(HM)) for individual one-to-one orthologs across human, lemur, and mouse., in a box plot (i) or a scatter plot (j). In panel i, bars show median (central line) as well as 25th (bottom) and 75th (top) percentiles. *p*-value calculated using paired two-tailed *t*-test. Dashed horizontal line indicates the median of *r*_g_(HL). In panel j, gray dots are different genes with the contours showing their probability density. Black dashed line indicates 1-1 relationship between the two coefficients. Blue and red dashed lines indicate the thresholds used to identify HL and HM genes, respectively. Green and red dots highlight examples of highly-conserved and HL genes, respectively, with their cell type expression patterns shown in panel k. See also Fig. S12a-c. **k**. Dot plots showing expression patterns of highly species-conserved (top) and HL-conserved/mouse-divergent (bottom) example genes in the 63 orthologous cell types across human, lemur, and mouse. Shown here are example HL genes exhibiting different patterns of evolutionary expression rewiring: [1] simple gain or loss of expression in the two primates vs. mouse; [2] conserved expression across the three species in some cell types and expression expansion and contraction in other cell types in primates vs. mouse; [3] expression switched from one (or more) mouse cell types to different cell type(s) in primates; and [4] complex expression rewiring with combinations of the above patterns. Rows are orthologous genes, indicated with the respective human gene symbols. Columns are cell types, displayed as triplets of the respective expression in humans, lemurs, and mice. Cell types are ordered first by compartments, then by tissue, and finally by species. **l-o**. Scatter plots showing expression along the spermatogenesis pseudotime for example genes with expression patterns that are highly-conserved (l), primate-conserved and mouse deviated (m, lack of expression in mouse; n, heterochromatic expression in mouse), or lemur specific (o). Points are single cells color coded by species. Solid curves show a moving average of the expression level along the spermatogenesis trajectory. See also Fig. S10d for more examples. **p-r**. Scatter plots showing expression along the pseudotime of neutrophil development for example genes with expression patterns that are highly-conserved (p) or primate-conserved and mouse deviated (q, lack of expression in mouse; r, heterochromatic expression in mouse). Points are single cells color coded by species. Solid curves show a moving average of the expression level along the neutrophil trajectory.

Another example of cross-compartment similarity was myelinating and non-myelinating Schwann cells (peripheral glia), which surprisingly segregated with stromal cells and away from central glia (e.g., oligodendrocytes) (Fig. 5a). Differential expression analysis identified genes enriched in stromal and Schwann cells but not other neural compartment cells (e.g., *COL3A1, LAMC1, SOCS3*) (Fig. 5e) and a complementary set enriched in neurons and central glia but not Schwann cells (e.g., *OMG, GPR137C, TCEAL3*). Many of the genes expressed in common by Schwann and stromal cells are components or regulators of extracellular matrix, suggesting a distinctive function of peripheral glia in collaborative remodeling of the matrix with surrounding stromal cells.

Within a compartment, cells of the same designated type or subtype generally clustered closely despite their different tissue origins (Fig. 5a, f, Fig. S6c-d), except for epithelial cell types which are highly tissue-specific and generally clustered with other epithelial cells from the same organ, even those for which there were cell types of the same designation in another organ (e.g., skin and tongue basal and suprabasal cell types) (Fig. 5g). One example of initially perplexing cross-organ similarity was a distinctive population of epithelial cells from the lung of one individual (L2), which curiously clustered closely with uterine epithelial cells; these were later shown to be lung metastases of a uterine endometrial cancer (accompanying manuscript^23^).

### 6. Evolution of gene expression in primate cell types

To elucidate how gene expression has changed during primate evolution, we compared the transcriptomic profiles of mouse lemur cell types from six organs (lung, skeletal muscle, liver, testis, bone marrow, spleen) to the corresponding cell types of human, mouse, and where available, macaque, using mouse as the non-primate outgroup (Fig. 6a, Table 5). To ensure comparisons were made across truly homologous cell types and minimize technical artifacts, we focused on our own human (Tabula Sapiens, Human Lung Cell Atlas)^21,46^ and mouse datasets (Tabula Muris Senis)^20^ obtained with the same scRNA-seq method and profiled by the same tissue expert labs, but also included additional testis data from mouse, human, and macaque (*Macaca mulatta*)^62,63^, and lung data from macaque (*Macaca fascicularis*)^64^. We re-clustered and re-annotated cells from human, mouse, and macaque datasets using the same pipeline and marker genes used for lemur. Assignments of homologous cell types were then refined and verified by demonstration of co-clustering of the expression profiles of the corresponding cell types using the scRNAseq integration algorithm Portal^65^, and the cross-species data alignment algorithm SAMap^66^ (Fig. S9, Fig. 6c, e, Fig. S10a, f). We restricted analysis at the gene level to the ∼13,000 one-to-one-to-one gene orthologs across the three main species (human, lemur, mouse), or the ∼12,000 when additionally including macaque, which we curated by combining homology assignments from ENSEMBL and NCBI (Table 4). The above analysis identified and validated 63 orthologous cell types across human, lemur, and mouse (18 with macaque), as well as continuous gradients of developing hematopoietic and male germ cells.

Comparison of the transcription profiles of each of the 63 orthologous cell types showed their transcriptional similarity across species ranged from a correlation coefficient *r*_c_ of 0.26 - 0.72 (0.63±0.06, mean±SD) (Fig. 6b). The transcriptomic similarity between the orthologous human and lemur cell type was almost always greater than that between the human and mouse cell type, as expected from the closer evolutionary relationship between humans and lemurs (Fig. 6b, g). A similar result obtained when we compared developing progenitors in the spermatogenesis program (Fig. 6c-d, Fig. S10a-b), and in the neutrophil, erythroid, and monocyte/macrophage hematopoietic lineages (Fig. 6e-f, Fig. S10f-k). Interestingly, the magnitude of cell type transcriptional differences across species, and the expected advantage of lemur over mouse in modeling the corresponding human cell type, differed by cell type and varied along the developmental trajectories, implying cell-type-specific rates of evolutionary diversification in their transcriptional programs. For example, developing male germ cells showed decreasing cross-species similarity (*r*_c_) and increasing human-lemur to human-mouse difference (Δ*r*_c_) along the developmental trajectory (Fig. 6d), implying more rapid molecular evolution of the late stages of spermatogenesis. In contrast, neutrophils showed increasing cross-species similarity (*r*_c_) and generally increasing human-lemur to human-mouse difference (Δ*r*_c_) as progenitors matured (Fig. 6f, Fig. S9j-k). Comparison to available transcriptomes of orthologous macaque cell types showed similar trends, with macaque cell types generally displaying greater transcriptional similarity to those of human (Fig. 6d, Fig. S11a), as expected from their closer phylogenetic relationship. Here too though there were exceptions. For several cell types, notably in the lung endothelial and stromal compartments, the lemur cell type better mimicked the human orthologue transcriptome than the macaque did (Fig. S11a), suggesting a high degree of evolutionary adaptation in gene expression for those cell types in the macaque lineage. Thus, although the transcriptional similarity of orthologous cell types and progenitors is generally consistent with expectations from the phylogenetic relationship of the species, individual cell types and differentiating progenitors have transcriptionally diversified at different rates during primate evolution, some so much so that they violate the phylogenetic expectations.

To provide molecular insight into these cell type specializations in primate evolution, we identified for each cell type the cell type-selective genes whose expression were conserved across all species analyzed, defining mammalian cell type core gene expression programs, along with genes whose expression was conserved in primates but not mouse and thus may contribute to primate-selective properties of the cells (Table 6). Each cell type had dozens to hundreds (range 18-595, 174±95 mean±SD) of genes with a primate-selective expression pattern. These included genes highly enriched (>5-fold greater expression, *p* < 1e-5)) in the primate cell types relative to the corresponding mouse cell types, and ones that were highly depleted (>5-fold lower expression, *p* < 1e-5), suggesting cell type specific gain or loss of expression, respectively, in the primate lineage (or the opposite in the mouse lineage).

We also analyzed the conservation of the global pattern of expression for each gene across the analyzed cell types in human, lemur, and mouse (Table 7, Fig. 6h-j, Fig. S12a-d, Fig. S13). Surprisingly, only a small fraction of the genes exhibited highly-conserved expression patterns (11% at *r*_g_ threshold of 0.8). These highly conserved genes were enriched for compartment or cell type-specific genes (e.g., *LAMP3* in AT2 cells, *TIMP3* in endothelial and stromal compartments) and genes encoding structural and regulatory proteins of the cytoskeleton, cilia, and extracellular matrix (e.g., *ACTA1* and *MYL1* in the skeletal muscle cells and *ACTA2* and *MYL9* in the cardiac muscle cells and pericytes) (Table 8, Fig. 6k, Fig. S13a). Curiously, multiple uncharacterized genes (e.g., *C5orf52, C11orf65, C11orf71, C22orf23*) exhibited conserved expression in ciliated cells and/or male germ cells, perhaps encoding specialized features of the cytoskeleton of these cells. The orthologous macaque genes were similarly expressed, but with exceptions (Fig. S11b). For example, potassium channel *KCNK3*, mutations in which cause pulmonary arterial hypertension in humans^67^, was selectively expressed in lung pericytes (but not muscle pericytes) of human, lemur, and mouse (Fig. 13a), but not of macaque (Fig. S11b).

The vast majority of genes (89%) displayed evolutionarily divergent expression patterns (*r*_g_<0.8) (Fig. 6j, Fig. S12a-c). Although expression pattern conservation was overall greater between human and lemur (*r*_g_(HL)) than between human and mouse (*r*_g_(HM)) (Fig. 6h-j), as expected from their phylogenetic relationship, there was a wide range in expression plasticity of individual genes, including 7% that showed extreme plasticity (*r*_g_<0.3) (Fig. S12a-c). Expression conservation did not correlate with coding sequence conservation (Fig. S12e-f), implying separate evolutionary diversification mechanisms and/or selective pressures for the expression control sequences and protein coding sequences at each gene.

Fig. 6j highlights genes (in red) whose expression pattern was selectively conserved in the primates (Human-Lemur or HL genes), and hence may contribute to primate-specific traits. Some showed simple gain (or loss) of expression in the primate lineage, involving a single cell type (e.g., *SFRP5, PLCB3*), several types (e.g., *GFRA1, NKD1*), or nearly all of the analyzed cell types (e.g., *HNRNPA1, USE1*) (Fig. 6k, Fig. S13b). Others showed expression expansion (or contraction) into more (or fewer) primate cell types than in mouse (e.g., *CA3, S100A14*), while some switched expression from one cell type to another between primates and mice (e.g., *CSRP3, PCK2*) (Fig. 6k, Fig. S13b). However, many HL genes displayed combinations of these types of differences from mouse, suggesting complex evolutionary rewiring of their expression patterns (e.g., *PIGR, C10orf53*) (Fig. 6k, Fig. S13b). In many cases, such expression rewirings were to cell type(s) in a different tissue compartment or distant organ (e.g. *HDC, FGFR3, FIBIN, EFHD1*) (Fig. 6k, Fig. S13b). HL genes did not show enrichment in specific GO terms, their diverse functions spanning immune regulation (e.g., *CXCL16, CX3CR1, LBP, SH2B3, VSIG4, ISG20, CD82*), cell growth and proliferation (e.g., *CDK6, ABL1, POLE3, CCNY, S100A14, HAUS1, ARPP19*), protein synthesis and degradation (e.g., *ASXL1, ZNF467, ATOH8, MSI2, CSDE1, SIAH2, ASB9, CTSF*), and cell and hormonal signaling (e.g., *HHIP, DLK1, SFRP5, NKD1, NKD2, FGFR3, GFRA1, PRKCZ, VIPR1*; see also accompanying hormone atlas^40^), whereas some remain poorly characterized (e.g., *C19orf47, C11orf24, C10orf53, WDR38, RNF150*) (Table 7). Many HL genes (26%) are linked to human diseases (Table 7; e.g., *PCK2* (PEPCK deficiency), *F2* (hemophilia a), *ARSA* (metachromatic leukodystrophy), *SH2B3* (familial erythrocytosis and thrombocythemia), *PNPLA6* (Laurence-Moon Syndrome), and *PSEN2* (Alzheimer’s disease)). In a similar way, we also identified lemur/mouse (LM) genes and human/mouse (HM) genes with divergent expression patterns in humans and in lemurs, respectively (Fig.S12a-d, Fig. S13c-d), which may contribute to human-selective and lemur-selective biology.

We carried out a similar analysis of the conservation of primate gene expression along the spermatogenesis developmental trajectory. Many of the genes that mark canonical stages of spermatogenesis are similarly expressed in all four species (e.g., *SALL4, PIWIL4*, spermatogonia; *SYCP2*, early spermatocytes; *ACR*, pachytene/diplotene spermatocytes; *SPACA1*, late spermatocyte to early spermatids; *FSCN3* spermatids; Fig. 6l, Fig. S10c-d), defining a conserved core program of mammalian spermatogenesis. However, we also identified primate-specific features of the program including genes expressed only in primate spermatogenesis (e.g., testis seminoma associated *L1TD1*^*68*^, phosphatase *PHOSPHO1*, kinectin *KTN1*) (Fig. 6m, Fig. S10d), along with dozens of primate-selective genes (orthologs identified in human and lemur but missing in mouse) that are selectively expressed during spermatogenesis (accompanying manuscript ^23^). We also found genes expressed in all species during spermatogenesis but the expression peaked at different times in primates vs. mouse, indicating heterochronic re-wiring of the spermatogenesis program during primate evolution (e.g., *PHOSPHO2*; serine protease *PRSS55*, required for male mouse fertility^69^) (Fig. 6n, Fig. S10d). Additionally, we found genes whose expression patterns during spermatogenesis were lemur-specific (e.g., *YPEL2, STX7, CA2*) (Fig. 6o, Fig. S10d). These genes are of particular interest because of their potential role in the remarkable radiation of the Lemuroidea clade (>100 species, nearly one-quarter of all primate species), and because several notable evolutionary specializations have already been recognized for mouse lemur, including the dramatic seasonal regulation of the testes and the role of sperm competition in reproduction^41^.

We similarly compared gene expression across the primate and mouse for three hematopoietic (myeloid) trajectories. Known marker genes generally exhibited conserved expression patterns (Fig. 6p, Fig. S10l-n), consistent with the higher cell type conservation observed in these immune trajectories compared to mid and late spermatogenesis (Fig. 6d, f, Fig. S11j-k). Nevertheless, we still identified genes (e.g., *MMP8*^*70*^, *OSCAR, RETN*) that were similarly expressed in the primates (human and lemur) but showed no expression or heterochronic changes in mouse hematopoiesis, particularly in the neutrophil lineage (Fig. 6q-r, Fig. S10o, see also accompanying manuscript^23^).

Extending such comparisons to all homologous cell types and more broadly across phylogeny will provide insight into the selection pressures and molecular changes that underlie the evolutionary specializations of primate cell types. Even at this stage the three- (lemur, human, mouse) and four-way (plus macaque) comparisons described here suggest interesting biological hypotheses and identify many cell types and genes for which lemur could provide a human modeling advantage over mice.

## DISCUSSION

Our single cell RNA-sequencing and analytical pipeline defined over 750 mouse lemur molecular cell types and their expression profiles, comprising nearly all major cell types for most (27) tissues. The cell types include cognates of most canonical human cell types plus stem and progenitor cells and their developmental programs for spermatogenesis, hematopoiesis, and other adult tissues, many of which did not form discrete molecular types but merged into continuous gradients of cell intermediates along the spatial or developmental pathways. Biologically-guided expert curation also uncovered dozens of previously unidentified or little characterized cell types, some of which appear to be conserved (e.g., two types of hepatocytes, several capillary types specialized for energy storage or activated by inflammation) but others may be primate or lemur-specific innovations.

By organizing the mouse lemur cell types by organ, compartment, and function, and then globally comparing their expression profiles in a simplified UMAP or matrix of all pairwise comparisons, we defined the molecular relationships of cell types across the body. This revealed global features like the high molecular similarity of cell types within some compartments (e.g., endothelial) but marked divergence of cell types in others (e.g., immune), as well as the surprising similarity of a few cell types across compartments, such as spermatogonia to hematopoietic progenitors and peripheral glia to stromal populations.

The atlas provides a broad cellular and molecular foundation for studies of this emerging primate model organism. Although the first steps in establishing a new model organism have traditionally been mutant screens and generation of a genetic map or reference genome, with the technological advances and declining cost of scRNA-seq, the creation of a reference transcriptomic cell atlas like ours can now be prioritized. It systematically defines cell types and aids elucidation of their functions and allows molecular comparisons of mouse lemur cell types to each other and to their orthologs in humans, macaques, and mice, allowing exploration of primate biology and evolution at cellular resolution. This revealed the many cell types (nearly all analyzed) and expressed genes (many implicated in human diseases) for which mouse lemur provides a human modeling advantage over mice and some even over macaque, as well as cases like sperm with a multitude of primate- and lemur-specific expression innovations. But the atlas also provides a powerful new way of detecting genes, defining their structures and splicing, and assigning their function, as well as elucidating organism-wide processes such as hormonal signaling, immune cell activation and inflammation, and primate-specific physiology, diseases, and genes, as we show in the accompanying papers (accompanying Tabula Microcebus manuscript^23^; hormone atlas^40^).

Our cell atlas strategy can be adapted to other emerging model organisms. This strategy should include the opportunistic identification of donors and systematic procurement of tissues from each; application of the scRNA-seq technologies and analytical pipeline including computational cell clustering, integration, and expert cell annotation; and the biological organization of cell types and comparisons across the organism and between organisms.

Importantly, extensive clinical and histopathological metadata should be collected from each donor and tissue to mitigate against complexities arising from genetic and environmental differences, as this can provide insight into individual-specific features of the atlas, as we exploit in the accompanying paper^23^. Application of this strategy to a wide variety of organisms^71–74^ will rapidly expand our cellular, genetic, and molecular understanding of biology that has been dominated for a half century by a small number of model organisms but now includes non-human primates.

## ACKNOWLEDGMENTS

This work was supported by funding from The Chan-Zuckerberg Biohub, Howard Hughes Medical Institute and Vera Moulton Wall Center for Pulmonary Vascular Disease; unrestricted grant from Research to Prevent Blindness and NEI P30-EY026877 to the Stanford Department of Ophthalmology to Albert Y. Wu; The Hong Kong University of Science and Technology start-up grant R9364, The Hong Kong University of Science and Technology Big Data for Bio Intelligence Laboratory (BDBI), and The Chau Hoi Shuen Foundation (R9056) to Angela Ruohao Wu; NSF-DBI-1701984 to Anne D. Yoder; funding from Developmental and Stem Cell Biology Graduate Program to Aris Taychameekiatchai; funding from Hong Kong Research Grant Council (16307818, 16301419, 16308120, 16307221, C6021-19E), the Hong Kong University of Science and Technology (startup grant R9405), and The Hong Kong University of Science and Technology Big Data for Bio Intelligence Laboratory (BDBI) to Can Yang; NIH grant R01 GM122951 and R35 GM136433 to Margaret Fuller; funding from Shanghai Sailing Program to Jingsi Ming (21YF1410600); NIA 1K99AG066963 to Thomas Ambrosi; NIH DP2AI138242 to Iwijn De Vlaminck; National Sciences and Engineering Research Council of Canada fellowship PGS-D2 to Michael F.Z. Wang; NIH R35 GM139517, R01 GM116847, R35 GM139517, NSF MCB1552196 to Julia Salzman; NSF Graduate Research Fellowship DGE-1656518 and a Stanford Graduate Fellowship to Julia Olivieri; Cancer Systems Biology Scholars Fellowship (Grant R25 CA180993) and Clinical Data Science Fellowship (Grant T15 LM7033-36) to Roozbeh Dehghannasiri; Urology Care Foundation Research Scholar Award Program and AUA Western Section Research Scholar Fund II to Hosu Sin; Stanford Graduate Fellowship/HHMI/NIH CMB Training Grant to Yue Zhang; American Cancer Society Postdoctoral Fellowship to SoRi Jang; Walter V. and Idun Berry Postdoctoral Fellowship to Andrea R. Yung; NSF Graduate Research Fellowship and Stanford Graduate Fellowship to Youcef Ouadah; the Wu Tsai Neurosciences Institute Interdisciplinary Scholar Award to Shixuan Liu; European Community’s 7th Framework Programme (FP7/2007-2013) under grant agreement Nbr 278486 (DEVELAGE), from the Fonds Unique Interministériel and Région Languedoc-Roussillon under grant agreement Nbr 110284 (DiaTrAl), and from the Fondation Plan Alzheimer (PRADNET) to Jean-Michel Verdier and Corinne Lautier; NIH R01 AI024258 to Peter Parham and Lisbeth A. Guethlein; NIH R01DC016892 to Wan-Jin Lu; NIH P30DK116074 to Yan Hang; NSF Graduate Research Fellowship to Connor V. Duffy; Postdoctoral Fellowships from the DFG (NE 2006/1-1) and California TRDRP (25FT-0011) to Patrick Neuhöfer; a NovoNordiskFonden Start Package grant (0071116) to Antoine de Morree; funding from Independent Research Fund Denmark (DFF–5053-00195) and Lundbeck Foundation (R232-2016-2459) to Jean Farup; NIH AG068667, AR073248 and AG036695 to Thomas A. Rando; funding from Wu Tsai Neurosciences Institute to Tony Wyss-Coray; NSF BCS 0647402 to Liza Shapiro and E. Christopher Kirk.

## METHODS

### Animal husbandry

All four gray mouse lemurs (*Microcebus murinu*s) included in this atlas originated from the closed captive breeding colony at the Muséum National d’Histoire Naturelle in Brunoy, France, and were transferred together to the University of Texas (Austin) in 2009 and then Stanford University in 2015 and maintained for noninvasive phenotyping and genetic research as approved by the Stanford University Administrative Panel on Laboratory Animal Care (APLAC #27439) and in accordance with the Guide for the Care and Use of Laboratory Animals, as detailed in Casey et al.^22^. Briefly, mouse lemurs were housed indoors in modified marmoset caging with multiple PVC perches and nest boxes in an AAALAC-accredited facility in a temperature (23.3-24.4°C) and light-controlled environment (daily 14:10 h and 10:14 h light:dark alternating every 6 months to synchronize seasonal breeding behavior and metabolic changes) and were fed *ad libitum* with fresh fruits and vegetables, crushed primate chow (Teklad Global 20% Protein Primate Diet #2050, Envigo), and live insect larvae as enrichment items. Animals were socially housed, in single sex groups, or individually housed due to behavioral incompatibility or health management requirements. Health and welfare were monitored daily and clinical care provided by the Veterinary Service Center, Stanford University, including diagnosis and treatment of spontaneously occurring health conditions. Animals in declining health despite medical care were euthanized for humane reasons as determined by a veterinarian. Prior to euthanasia, it happens that all four lemurs were living in summer-like long days (14:10 h) for at least 3 months (range 3-6 months), and all showed standard activity patterns without signs of torpor. Given these individuals were housed at constant temperature conditions and fed a non-calorie restrictive diet, spontaneous torpor was not observed in any of the analyzed lemurs throughout their time in the Stanford colony.

### Tissue procurement and processing

Animals in declining health that did not respond to standard therapy were euthanized by pentobarbital overdose under isoflurane anesthesia as described in Casey et al.^22^. Prior to euthanasia, a veterinary examination was performed, and animal body weight and electrocardiogram (ECG) were obtained (KardiaMobile 6L, AliveCor). Blood was immediately collected via cardiocentesis for serum chemistry, complete blood count, biobanking, and single cell RNA-sequencing. In three animals (L2, L3, and L4), transcardial perfusion of the lungs with phosphate buffered saline (PBS) was done to reduce circulating cells. Organs and tissues were sequentially removed and divided by a veterinary pathologist. One sample of each tissue was immediately placed in formalin fixative for histopathology^22^, and a second was embedded in Optimal Cutting Temperature compound (OCT) and then flash frozen on dry ice and stored at - 80°C for biobanking. A third sample was placed directly in cold (4°C) PBS pH 7.4 and immediately distributed to the tissue expert for cell dissociation and preparation for scRNA-seq as detailed below. Additional diagnostics such as microbiological cultures were performed where clinically indicated. The entire necropsy was completed within 1-2 hrs, with ischemia-sensitive tissues prioritized as described in Supplementary Methods.

### Histological and pathological analysis

Tissues were immersion-fixed in 10% neutral buffered formalin for 72 hours. Formalin-fixed tissues were processed routinely, embedded in paraffin, sectioned at 5 μm, and stained with hematoxylin and eosin (H&E). Tissues included the following: heart, aorta, lungs, trachea, thyroid gland, parathyroid gland, kidneys, urinary bladder, male reproductive tract (testicle, epididymis, seminal vesicle, prostate, and penile urethra), female reproductive tract (uterus, cervix, vagina, and ovaries), salivary glands, tongue, epiglottis, esophagus, stomach, small and large intestine, liver (with gallbladder), adrenal gland, spleen, lymph nodes, white adipose, brown adipose, bone, spinal cord, eyes, and bone marrow. Selected tissues stained with Von Kossa (for mineralization), Masson’s trichrome (for collagen), Congo Red (for amyloid), and Gram stain (for bacteria) as part of the pathological analysis. H&E stained slides were scanned with a Leica Aperio AT2 High Volume Digital Whole Slide Scanner (40x objective), uploaded into Napari image viewer^75^, software adapted by CZB, and posted on the Tabula Microcebus portal.

### Preparation of single cell suspensions and FACS-sorting for scRNA-seq

Fresh tissue samples procured as described above were placed on ice, received by organ/tissue experts and immediately dissociated and processed into single cell suspensions, except for samples from L3, which were kept cool overnight after necropsy and processed the next morning (see Supplementary Methods). For each solid tissue, this involved a standard combination of enzymatic digestion and mechanical disruption methods that were optimized for the specific tissue, many of which were adapted from procedures used for the corresponding mouse tissue^19,20^. For blood, immune cells were isolated using a high density ficoll gradient (Histoplaque-1119, Sigma-Aldrich) to include peripheral blood mononuclear cells (PBMCs) and polymorphonuclear leukocytes (PMNs)^21^.

The specific protocols for each of the 27 tissues are detailed in Supplementary Methods. Cell number and concentration for each single cell suspension were determined by manual counts using a hemocytometer, and then adjusted with 2% fetal bovine serum (FBS) in PBS to a target concentration of about 10^6 cells/mL. Samples were then used for droplet-based 10x library preparation and/or flow sorted for single live cells (Sytox blue negative; ThermoFisher S34857) for plate-based Smart-seq2 library preparation. To enrich for cardiomyocytes, the standard procedure for cardiac cell isolation was supplemented by hand-picking cardiomyocytes (Supplementary Methods). Residual cell suspensions were diluted 1:1 with serum-free Bambanker cell freezing media (GC Lymphotec #BB01) and cryopreserved at -80°C.

### scRNA-seq library preparation, quality control, and sequencing

For 10x, single cells were profiled using the 10x Genomics single cell RNA-sequencing pipeline (Chromium Single Cell 3’ Library and Gel Bead v2 Chemistry kit) and sequenced on a NovaSeq 6000 System as previously described^19–21^ and detailed in Supplementary Methods. For SS2, single cells were sorted into 384- or 96-well lysis plates, reverse transcribed to complementary DNA (cDNA) and amplified, as previously described^19,20^. cDNA libraries were prepared using the Nextera XT Library Sample Preparation kit (Illumina, FC-131-1096) or (for L4) an in-house protocol detailed in Supplementary Methods; no significant differences between protocols were observed in library read depth or quality. Pooling of individual libraries and subsequent quality control and DNA sequencing were done as previously described^19–21^ with minor modifications (Supplementary Methods). Both 10x and SS2 libraries were sequenced to achieve saturation on an Illumina NovaSeq 6000 System (10x, 26 bp and 90 bp paired-end reads; SS2, 2 x 100 bp paired-end reads).

### Genome alignment of scRNA-seq sequencing reads and gene counts

Microcebus murinus genome assembly (Mmur 3.0, accession: GCF_000165445.2; annotation: NCBI Refseq Annotation Release 101) with NCBI annotation release 101 (date acquired, September 21, 2018) was used for downstream alignment and data analysis. A total of 31,509 genes were detected, including annotated genes and unannotated loci but excluding mitochondrial and Y-chromosome genes (unannotated at our acquisition date).

For 10x samples, downstream data was processed by standard methods using Cell Ranger (version 2.2, 10x Genomics). Raw base call (BCL) files directly generated by the NovaSeq were demultiplexed and converted to FASTQ files, and then aligned to the 10x genome index, with barcode and UMI counting performed to generate a gene counts table. Alignment files were outputted in standard BAM format.

For SS2 samples, demultiplexed fastq files were mapped to the genome using STAR aligner (version 2.6.1a). Briefly, the genome FASTA file was augmented with ERCC sequences to create a STAR genome index with 99 bp overhangs (optimized for Illumina 2 x 100 bp paired-end reads). Two-pass mapping was executed, in which the first pass identified splice junctions that were added to the gene reference to improve second pass mapping, with specific STAR options and parameters detailed in Supplementary Methods.

### Contamination filtering of 10x data

We performed stringent contamination filtering to resolve cross-sample contamination in an Illumina sequencing run caused by cell barcode hopping among multiplexed 10x samples^76,77^. Such cross-sample contamination can occur when low levels of ambient mRNA containing the 10x cell barcode in one sample gets added onto the transcript of other samples during Illumina sequencing amplification, resulting in the incorrect assignment of a cell barcode to other samples; hence in subsequent analyses, a cell from one tissue could falsely appear as multiple cells from different tissues (samples). To exclude such artifacts, for each sequencing run we identified all cell barcodes that were assigned to multiple samples, and for each such barcode identified we compared the number of UMIs in each sample. If there was one dominant sample index (i.e., number of UMIs of the dominant sample was 10 times or more greater than that of the second most abundant sample), then the cell with the dominant sample index was kept (but labeled in its metadata as ‘potentially contaminated’) whereas all other instances of that “cell” were removed. If there was no dominant sample index, then all instances of the “cell” with that barcode were removed from the dataset. (This was not an issue for SS2 samples because they were sequenced using dual unique indices for each cell.)

### Cell clustering, annotation, and cluster markers from scRNA-seq profiles

#### Step 1 – Cell clustering and annotation of each tissue processed by 10x

Transcriptomic profiles of cells from each tissue from each individual lemur were clustered separately using Seurat software package (version 2.3.0) for R studio (version 3.6.1). We included in this step all cells with >100 genes or >1000 UMIs detected, a minimal threshold that was used to ensure inclusion of all cell types including ones in which the cells (or RNA) were unstable (see below for more stringent criteria used for final cell quality control). For each cell, expression of a gene *g* is normalized in 10x data as: ln(UMI_*g*_/UMI_total_ *1e4 +1), abbreviated as ln(UP10K+1); in SS2: ln(reads_*g*_/reads_total_ *1e4 +1), abbreviated as ln(CP10K+1). Next, data scaling, dimensionality reduction (PCA), clustering, and visualization (t-SNE, UMAP) were performed following the standard Seurat pipeline as previously described^21^ with parameters including the numbers of principal components, perplexity, and resolution adjusted manually for each iteration of cell clustering. Resultant cell clusters were manually assigned to a compartment (endothelial, epithelial, stromal, lymphoid, myeloid, megakaryocyte-erythroid, neural, germ) based on expression of the mouse lemur orthologs of canonical marker genes for each compartment in human and mouse (see Table S2). Clusters expressing markers from more than one compartment were annotated as “doublets.” Cells within each assigned compartment were then subclustered, repeating the data processing steps above, and then the clusters in each compartment were annotated separately. To annotate (determine) the cell type of each cluster, a list of canonical human and mouse gene markers for each cell type in each tissue was curated from the literature (Table S2), including genes previously validated by in situ hybridization and/or immunohistochemistry as well as differentially-expressed genes culled from recent scRNA-seq studies, and the orthologous mouse lemur genes were identified and their expression visualized on the t-SNE plots. Based on the enriched expression of marker genes, each cluster of cells within a compartment was manually assigned a cell type identity. Clusters that contained more than one cell type were further subclustered to better resolve the cell types. Cell types represented by only a small number of cells that did not form a separate cluster were manually curated, aided by the cellxgene gene expression visualization tool^78^ as detailed below.

Each cluster was assigned both a ‘cell ontology’ cell type designation using the standardized and structured nomenclature^27^, and a ‘free annotation’ that resolves biologically significant clusters not contained in the current cell ontology. Free annotations were assigned as follows. In cases where a smaller cluster stemmed off a larger (“main”) cluster in t-SNE embedded space, the smaller cluster was distinguished with one or more differentially-expressed genes added to the cell type name (e.g. B cell (*SOX5+*) clustered near the main population of B cells in the pancreas); differentially-expressed genes driving the subtype clustering were ascertained by Wilcoxon rank-sum tests. In cases where two approximately equal-sized clusters separated on the t-SNE plot, a marker gene was added to the cell type name for both clusters. Clusters with a small number of cells that contained more than one cell type but could not be partitioned into separate clusters by subclustering with the Louvain algorithm or manually with cellxgene (see step 3) were labeled as a ‘mix’ cell type (e.g. the cluster labeled ‘endothelial cell’ in the uterus contains a mixture of artery, vein, and capillary cells). Clusters with cells that expressed markers for more than one cell type and it was biologically plausible they were not a technical artifact (e.g., doublet of two distinct cell types) were labeled as a ‘hybrid’ cell type (e.g. the cluster labeled as ‘monocyte/macrophage’ in the trachea contains cells that expressed markers of both cell types and could not be further distinguished based on current molecular definitions of these cell types). After examining the human/mouse markers for all known cell types in a tissue, clusters that could not be assigned a cell type were labeled ‘unknown,’ with the tissue, compartment, and one or more differentially-expressed genes added to the cell type name (e.g. ‘unknown_Bone_stromal_G1 (*NGFR+ TNNT2+*)’ are bone stromal cells that do not correspond to any extant stromal cell type reported for human and mouse). To detect the differentially-expressed genes of an unknown cell type, we compared the unknown cell type to all other cells of the same compartment and tissue (see Fig. S5). Clusters containing a majority of cells that expressed cell proliferation markers (e.g. *TOP2A, MKI67, STMN1*) were appended the abbreviation “PF.” Clusters that separated from a main cluster but did not express any distinguishing markers (other than tRNAs, rRNAs, and/or immediate-early genes) and differed only in parameters of technical quality (i.e. fewer genes and counts detected per cell) were considered low quality and “LQ” was appended to the cell type name.

After annotations were assigned, the cutoff for the minimum number of genes per cell was increased from 100 to 500 and only the qualifying cells were further analyzed. For most tissues, this more stringent cutoff only resulted in removal of some erythrocytes and neutrophils; the only exception were cardiac cardiomyocytes, most of which expressed fewer than 500 genes per cell so separate filtering criteria were applied (see Supplementary Methods).

#### Step 2 – Annotation of each tissue processed by SS2

Cells processed by the SS2 protocol with <500 genes or <5000 reads were excluded from further analysis, and gene expression levels in the remaining cells were scaled and log transformed as described above for the 10x datasets. Cells from a particular tissue and individual were integrated with the 10x dataset of the same tissue and individual into the same UMAP embedded space using the FIRM algorithm (detailed below). Cells from SS2 were automatically annotated with the same label as the nearest neighboring 10x cell. Annotations were manually verified in Step 3 aided by cellxgene gene expression visualization. SS2 datasets for which there were no corresponding 10x dataset from the same individual/tissue were manually annotated using the method described in Step 1 for 10x datasets.

#### Step 3 – Integration of datasets across individuals

For each tissue, the combined 10x/SS2 datasets from each individual were further integrated into the same UMAP embedded space using the FIRM algorithm^26^. This step resulted in 27 separate tissue UMAPs, each containing data from up to 4 individuals. To ensure consistency of cell type labeling across all individuals, annotations were verified and adjusted manually using cellxgene, an interactive tool to visualize and annotate single cell RNA-sequencing data (https://chanzuckerberg.github.io/cellxgene/)^78^.

#### Step 4 – Integration of datasets across tissues

All 27 tissue-level objects were integrated into a single UMAP embedded space using the FIRM algorithm. As above, annotations were verified/adjusted manually in cellxgene to ensure consistency of cell designations across all tissues. In most instances, each cell type clustered separately, irrespective of the tissue of origin, and the same designation was used across all tissues. Occasionally, similar cells types (e.g. fibroblasts, macrophages) clustered separately by tissue of origin, making it challenging to distinguish whether the separation was due to tissue-level batch effect or because of true biological differences. In these cases, the original tissue-level annotation label was kept for each cluster.

#### Step 5 – Detect differentially-expressed genes for each cell type

We calculated the top 300 differentially-expressed genes (adjusted *p*-value<0.05) for each cell type in the 10x dataset (represented by at least 5 individual cells, after removing doublets, low quality, and mixed cell types) using the Wilcoxon rank-sum test with Benjamini-Honchberg FDR correction (Table 3). We compared each cell type to: (i) all other cell types from the same tissue (e.g., capillary cell type of the lung compared to all other lung cell types, ‘tissue-wide’ comparison), (ii) all other cell types from the same compartment of that tissue (e.g., capillary cell type of the lung compared to all other lung endothelial cell types, ‘tissue-compartment-wide’ comparison), (iii) all other cell types from the atlas (capillary cell type of the lung to all other cells in the atlas, ‘atlas-wide’ comparison), (iv) all other cell types from the same compartment across the atlas (capillary cell type of the lung to all other endothelial cell types in the atlas, ‘atlas-compartment-wide’ comparison).

### FIRM integration

FIRM is a newly developed algorithm that integrates multiple scRNA-seq datasets^26^ (e.g., from different sequencing platforms, tissue types, and experimental batches). In brief, FIRM optimizes dataset integration by harmonizing differences in cell type composition and computing the dataset-specific scale factors for gene-level normalization. Different datasets generally have varied cell-type compositions, resulting in dataset disparity when scaling the gene expression levels to unit variance for each dataset. Different from classical scaling procedures, FIRM computes the scale factors based on subsets of cells which have matched cell-type compositions between datasets. To construct these subsets, FIRM detects paired clusters between datasets based on similar overall gene expression and then samples the cells so that paired cell types have the same proportional representation in each dataset. Parameters used for integration are given in Supplementary Methods. The integrated datasets generated using FIRM show accurate mixing of shared cell-type identities and preserve the structure of the original datasets, as confirmed by expert manual inspection during cell annotation.

### Trajectory analysis

We used two independent methods to detect and characterize spatial and developmental pseudotime cell trajectories: 1) a custom in-house program in Matlab and 2) Slingshot^79^. For the mouse lemur kidney nephron spatial trajectory, all kidney epithelial cells were included in the analysis; podocytes, macula densa cells, intercalated cells, and urothelial cells were not part of nephron gradient and hence excluded from trajectory detection. For vasa recta endothelium spatial trajectory, all four vasa recta cell types were used. For spermatogenesis pseudotime trajectory, all seven sperm and sperm progenitor cell types were used. For the myeloid cell developmental pseudotime trajectory, hematopoietic precursor cells and all myeloid cell types except dendritic cells (which did not form part of the continuum) were used. Analysis was performed independently for each trajectory using values from the 10x scRNA-seq profiles of the indicated cells (low quality cells and technical doublets were excluded) that had been pre-processed (scaled: (ln(UP10K +1), normalized) as described above.

Principal component analysis (PCA) with highly-variable genes (dispersion > 0.5) was done with the PCA function of Matlab, and the high quality principal components (not driven by extreme outlier data points or immediate-early genes) were selected from the top 20 principal components and used to generate a 2D UMAP using cell-cell Euclidean distances as input (https://www.mathworks.com/matlabcentral/fileexchange/71902). The trajectory of the cell continuum was detected as the probability density ridge of the data points in the UMAP, using automated image processing (Matlab Image Processing Toolbox™); any interruptions in the detected density ridge line were connected manually along the direction of the ridge line and guided by prior knowledge of the biological process, and the direction of the trajectory was assigned based on expression of marker genes. Individual cells were then aligned to the trajectory by the shortest connecting point to the trajectory; if the trajectory branched (e.g., in myeloid cell development), cells were assigned to the closest branch. Individual cells that were too distant from the trajectory (adaptive thresholding along the trajectory) were deemed outliers and removed from further analysis.

To detect genes whose expression followed the trajectory, we calculated the Spearman correlation coefficient and corresponding *p*-values (Bonferroni corrected) between the expression level of each gene and 20 preassigned unimodal patterns that smoothly change along the trajectory (with their single peaks uniformly distributed from the beginning of the trajectory to its end point). Expression patterns of the top ranking (top 1000 with *p*-value<0.01) and highly variable (dispersion > 0.5) genes were smoothed with a moving average filter and clustered by k-means clustering to detect the major trajectory-dependent expression patterns. The trajectory differentially-expressed genes were then ranked by the associated cluster (ranked by trajectory location of peak expression), and within the cluster by *p*-value from smallest to largest, and with the same *p*-value by mean expression level from highest to lowest.

For the myeloid cell analysis, four trajectories were independently detected: 1) from hematopoietic precursors to granulocyte monocyte progenitors, 2) from granulocyte monocyte progenitors, proliferating neutrophils, to neutrophils, 3) from granulocyte monocyte progenitors, proliferating macrophages/monocytes, proliferating monocytes, to monocytes and macrophages, and 4) from megakaryocyte progenitors, erythroid progenitor cells, proliferating erythroid lineage cells, to erythroid lineage cells. On the UMAP, trajectory 1 branched into trajectories 2 and 3, so two longer trajectories were generated (1+2, 1+3). Differential gene expression analysis was then independently performed for each of the constituent trajectories (1+2, neutrophil lineage; 1+3, monocyte/macrophage lineage; 4, erythrocyte lineage).

As an alternative method, we also applied the Slingshot method^79^, which first computes the global lineage structure by constructing a cluster-based minimum spanning tree (MST) followed by pseudotime inference using simultaneous principal curves to fit smooth branching curves to these lineages. We used the annotated clusters and UMAP coordinates to first obtain a global lineage structure with ‘getLineages’, then constructed smooth curves, ordered cells along the trajectory, and generated pseudotime values using ‘getCurves’. For each tissue, the longest trajectory that incorporated the most clusters were used. For the immune trajectories, the neutrophil cluster was subdivided into higher resolution clusters that were combined with the main annotations to facilitate MST building. For each trajectory, coordinates were normalized by the maximal value for comparison with the other method.

### Comparison of expression profiles among mouse lemur cell types

#### UMAP of cell types

To visualize similarities among the mouse lemur cell type expression profiles, we applied uniform manifold approximation and projection (UMAP) and embedded the high dimensional scRNA-seq expression data (∼30,000 genes) to 2D. For this analysis, the 10x scRNA-seq dataset was used and molecular cell types that were low quality (labeled with LQ in free annotation) or represented by less than 4 individual cells were excluded, resulting in a comparison of 681 molecular cell types. Molecular cell types were treated as pseudo bulk, with gene expression levels calculated by averaging the expression level of each gene for all cells of that type and then natural log transformed (ln(Avg Count Per 10K UMIs +1)). Expression levels were further normalized by the maximal value of each gene across all cell types, so that all ranged from 0 to 1. The cell type gene expression matrix was then projected onto a 2D space with cosine distances between pairs of cell types used in the UMAP function (https://www.mathworks.com/matlabcentral/fileexchange/71902). The Wilcoxon rank-sum test was used to identify differentially-expressed genes that distinguished related molecular cell types identified in the cell type UMAP from other cell types (e.g., sperm/sperm progenitor cells and immune progenitor/proliferating cells vs. proliferating cells of other compartments) as described in Supplementary Methods.

#### Heat map of cell type pairwise correlation scores

To compare the overall gene expression profiles of molecular cell types, Pearson’s correlation scores were calculated for every pair of molecular cell types. To compare data from different sequencing platforms (10x and SS2), we used the FIRM integrated dataset as described above which contains FIRM-generated principal component (PC) coefficients for each cell. Cell types were treated as pseudo-bulk and the cell type average PC coefficients were calculated and used to determine the correlation coefficients. The cell type pairwise correlation scores were plotted as heat map matrices with cell types arranged in three different orders in the matrices to facilitate comparison (Fig. 5 g-h and Fig. S7). Interactive forms of the heat map matrices are available online at Tabula Microcebus portal.

### Evolutionary comparison of mouse lemur, human, macaque, and mouse transcriptional profiles

#### Step 1 - Compiling comparable datasets

For the comparisons, we used published human, mouse, and macaque scRNA-seq (and not single nucleus) datasets obtained using approaches and standards similar to those described above for mouse lemur, and re-annotated where necessary for consistency with the lemur annotations. We compared cells of the lung, skeletal muscle, liver, testis, bone marrow, and spleen. For lung and muscle, all cell types were included in the analysis, whereas we only analyzed the epithelial cells for the liver, immune cells for the bone marrow and spleen, and germ cells for the testis. All lemur data was from the 10x data of this study plus additional muscle data from L5 (de Morree et al., in prep). Human data were from the 10x data of the Tabula Sapiens^46^, except for the lung which we used the 10x data from the Human Lung Cell Atlas^21^ and the testis which we used drop-seq data from Shami et al.^62^. Mouse data were from 10x data of the Tabula Muris Senis^20^ except for the testis which we used 10x data from Ernst et al.^63^. Given the limited data availability (either lack of the tissue or relevant cell types), we only performed cross-species for the lung and testis for the macaque data. Crab-eating macaque lung data were from the 10x data by Qu et al.^64^ and rhesus macaque testis data from Shami et al.^62^. All datasets profiled adult animals except for the two mouse datasets which also included data from animals during postnatal development. Thus, we used only data from adult mice to increase consistency of the datasets.

#### Step 2 - Orthology mapping across species

For orthology mapping, we merged the orthology databases from both the National Center for Biotechnology Information (NCBI) and Ensembl (Table 4, 5). We began by compiling all mouse lemur genes annotated in NCBI (mouse lemur taxonomy ID: 30608), then merged the corresponding human and mouse orthologs from NCBI (gene_info.gz and gene_orthologs.gz from https://ftp.ncbi.nlm.nih.gov/gene/DATA/, February 2020). We next added Ensembl gene ID numbers, gene names, and human/mouse ortholog assignments from Ensembl Biomart (Ensembl Genes version 99, February 2020) using the Ensembl gene ID (variable ‘Gene_stable_ID’) for each NCBI gene ID (variable ‘NCBI_gene_ID ‘) in Ensembl Biomart. Mouse lemur genes that did not have an assigned human and mouse ortholog in either Ensembl or NCBI were removed, as were mouse lemur genes that had more than one human or mouse ortholog assigned, or that shared the same human or mouse ortholog with another mouse lemur gene. Note that unlike NCBI, Ensembl specifies the type of ortholog assignment (e.g., ‘ortholog_one2one’, ‘ortholog_one2many’); however, we did not use the Ensembl specification to filter one-to-one-to-one orthologs because occasionally a mouse lemur gene name was assigned by homology to multiple currently unnamed loci in Ensembl and because of this imperfect genome annotation was labeled as sharing an ‘ortholog_one2many’ with human/mouse instead of ‘ortholog_one2one’. Finally, we appended the one-to-one orthologs between human and rhesus macaque and between human and crab-eating macaque, as assigned by Ensembl. A total of ∼15,000 one-to-one gene orthologs were thus uncovered across human, lemur, and mouse, ∼14,000 across human, lemur, mouse and rhesus macaque, and ∼13,000 across human, lemur, mouse and crab-eating macaque (Table 4). Sequence identity was based on those reported in the Ensembl homology database.

#### Step 3 - Integrating cross-species datasets and unify cell type annotations/designations

For the cross-species comparisons, we used the one-to-one orthologs that exist in all relevant datasets. Orthology mapping for the datasets was based on the NCBI or Ensembl gene ID, if the original datasets provided the respective gene ID, and on the gene symbol if gene ID was not provided. The choice of NCBI versus Ensembl depends on which version of the genome annotations the original dataset was aligned to. Some of the one-to-one orthologs were missing from one or more of the datasets, therefore were removed from the cross-species comparison. Together, we identified ∼13,000 genes for the comparisons across human, lemur, and mouse, and ∼12,000 genes for the comparisons that also include either of the macaque species.

To unify cell type annotations, human, mouse, and macaque datasets were first re-annotated separately for each tissue and species using the same pipeline and marker genes as for the lemur data. For the male germ cells which formed a molecular gradient. We simplified the annotations into three discrete stages, the spermatogonia, spermatocytes, and spermatids, based on their original annotations, and also applied trajectory analysis (see below). Next, to ensure consistency of cell annotations/designations across species, we applied Portal^65^ to integrate data from different species. Through adversarial learning of neural networks, Portal projects data into a space that minimizes species differences, from which an integrated UMAP is generated to visualize cell clustering from different species. Portal integration was performed separately for each tissue, except for bone marrow and spleen which were integrated jointly. We manually inspected each integration UMAP and ensured that cells of the same designation showed reasonable cross-species co-clustering and separation from other cell types. We also made small modifications to the cell annotations during this process to unify designations across species. For example, proliferating cells might co-cluster with the main non-proliferating population in the original dataset if the number of proliferating cells are too few, but they often form a separate cluster together with the proliferating populations of the other species in the integrated UMAP. In such a scenario, we would separately annotate these cells as a proliferating subtype. Additionally, we merged cell types and subtypes that had unclear cross-species correspondence but were almost indistinguishable in the species-integrated UMAP (e.g., proliferating T, NK and NKT cells were grouped together and designated NK/T cells (PF)).

As additional validations of annotation consistency across species, we also applied SAMap^66,80^, a self-assembling manifold algorithm and graph-based data integration method to the lung and muscle datasets to identify homologous (reciprocally-connected) cell types based on shared expression profiles across species. Cross-species cell types similarity (visualized by edge width in Fig. S8 a, c), is defined as the average number of cross-species neighbors of each cell relative to the maximum possible number of neighbors in the combined manifold. The default SAMap parameters were used in the analysis, and similarity scores less than 0.1 were removed.

#### Step 4 - Identify species-unified trajectories

Trajectories were calculated for spermatogenesis across human, macaque, lemur, and mouse, as well as three myeloid lineages (neutrophil lineage, monocyte/macrophage lineage, and erythroid lineage) of hematopoiesis across human, lemur, and mouse (macaque data not available). Trajectory detection and cell alignment was performed using the same custom in-house program as described above (in Trajectory analysis), with the species-integrated UMAPs as input.

#### Step 5 - Calculate cross-species similarity scores for each cell type

Cell types with more than 15 cells in each of the relevant species were used for the cross-species comparisons. This resulted in a total of 63 orthologous cell types for the comparisons across human, lemur, and mouse (63*3=189 total cell type entries across all species), and 18 cell types for the comparisons of the lung cell types across human, macaque, lemur, and mouse (18*4=72 cell type entries). Cell type mean gene expression was calculated for each gene. Single cell expression levels in the species-integrated dataset were normalized and log-transformed the same way as described above for the lemur-only dataset, i.e., ln(UMI_*g*_/UMI_total_ *1e4 +1). Note, however, that because there were fewer genes in the cross-species dataset (only one-to-one orthologs), the absolute expression levels were higher than that in the lemur-only dataset.

We used correlation coefficients as a proxy for cross-species similarity. To score similarity for individual cell types, we calculated Spearman rank-based correlation coefficients of cell type mean expressions between human and lemur, *r*_c_(HL), and between human and mouse, *r*_c_(HM). The cell type mean expression levels were thresholded at 0.4 to mitigate the effect of background noise. Cross-species similarity was similarly calculated for cells at different stages/lineages of spermatogenesis and hematopoiesis by applying a moving window along the respective trajectories.

#### Step 6 - Calculate cross-species similarity scores for each gene

Cross-species similarity was calculated separately for individual genes using the tissue and three-species (human, lemur, mouse) integrated dataset across the 63 orthologous cell types. We first quantified the mean expression (*E*_max_) in the maximally expressed cell type in each species. Next, we filtered genes that were not expressed or lowly expressed across the analyzed cell types, requiring *E*_max_>0.5 in each species or *E*_max_>0.1 in each species and *E*_max_>1.5 in at least one species. This resulted in a total of 7787 genes for follow-up analysis. Mean cell type expression levels across the 63 cell types were then normalized by *E*_max_ in each species and Pearson’s correlation coefficients between human and lemur (*r*_g_(HL)), between human and mouse (*r*_g_(HM)), and between lemur and mouse (*r*_g_(LM)) were calculated. We then calculated Δ*r*_g_ = *r*_g_(HL) - *r*_g_(HM) for each gene and tested the *p*-value for Δ*r*_g_ being significantly higher (right-tailed) or lower (left-tailed) than 0 (see below). To identify genes with human/lemur-conserved but mouse-divergent expression patterns (i.e., HL genes), we applied a threshold of Δ*r*_g_>0.4 and a right-tailed *p*-value< 0.05. We also identified human/mouse-conserved and lemur-divergent (HM) genes and lemur/mouse-conserved and human-divergent (LM) genes using the same threshold levels. Similar analysis was also performed to detect genes that show a species-conserved or -diverged expression patterns along the spermatogenesis trajectory and the neutrophil lineage of the hematopoiesis trajectory. Additionally, we identified genes that are highly-conserved (*r*_g_>0.8), lowly conserved (*r*_g_<0.3), or moderately conserved (*r*_g_>0.3 and *r*_g_<0.8). Full list of analyzed genes and their statistics is provided in Table 7. Expression patterns of representative genes visualized in Fig. S13. Gene set enrichment analysis was performed using g:Profiler^81^ for the highly-conserved genes and HL genes, with all the analyzed genes provided as a custom background gene set and otherwise default parameters.

To test if one correlation coefficient (*r*) is significantly higher or lower than the other, we estimated the statistical significance of their difference (Δ*r*) being larger or smaller than 0, through Fisher’s Z-transformation^82^. In essence, correlation coefficients, which were bounded and not normally distributed, were Fisher’s Z-transformed to the unbounded, and approximately normally distributed space by inverse hyperbolic tangent function and their difference and respective *p*-value calculated by standard one-tailed *t*-test in this transformed space. For display purposes, the mean and 95% confidence intervals of were then inverse transformed and displayed in Fig. 6b, Fig. S11a. Note that this inverse transformed Δ*r*, which is bounded between -1 and 1, does not necessarily equals the initial Δ*r* which is between -2 and 2.

#### Step 6 - Identify genes with primate-selective expression for each cell type

Using the cross–species dataset across human, lemur, and mouse, we performed two-tailed Wilcoxon rank-sum tests comparing expression in lemur vs. human, lemur vs. mouse, and human vs. mouse, independently for each gene and each homologous cell type. We also performed, for each cell type, one-tailed Wilcoxon rank-sum tests comparing expression in the cells of the cell type vs. all rest cells in the dataset. We also calculated fold-change in average expression for the above comparisons. Next, for each cell type, we searched for three categories of genes. First, genes with significantly primate-enriched expression, which requires (1) cell type average expression of the gene is above 0.5 in both primates, and (2) 5-fold greater expression and *p* < 1e-5 when comparing both the human and lemur cell types vs. the homologous mouse cell type. Second, genes with significantly primate-depleted expression, which requires (1) cell type average expression of the gene is above 0.5 in mouse, and (2) 5-fold lower expression and *p* < 1e-5 when comparing both the human and lemur cell types vs. the homologous mouse cell type. Third, genes that are significantly enriched in the cell type compared to the other cell types, regardless of the species, which requires (1) cell type average expression of the gene is above 0.5 in all three species, and (2) 5-fold greater expression and *p* < 1e-5 when comparing this cell type vs. other cell types. The full list of the identified genes is provided in Table 6.

## DATA & CODE AVAILABILITY

scRNA-seq gene expression Counts/UMI tables, and cellular metadata are available on figshare (https://figshare.com/projects/Tabula_Microcebus/112227). Data can be explored interactively using cellxgene on the Tabula Microcebus portal: https://tabula-microcebus.ds.czbiohub.org/. Raw sequencing data (fastq files) are available at https://app.globus.org/file-manager?origin_id=c9fc0a15-54a0-4182-8d64-fd8afc12f1fc&origin_path=%2F.

## TABLES

**Table 1. Marker genes from human and/or mouse used to annotate the lemur cells from the atlas**

The table lists for each classically defined histological cell type (rows) the known marker genes from corresponding human and/or mouse cell types as well as related references. Marker genes are listed in lemur symbols followed by special notes in brackets (see “Marker gene key”). Cell types are ordered by source tissue with cell types that exist in multiple tissues on the top. For each cell type, the table also lists the corresponding cell ontology assignments, tissue source, compartment assignment, and whether it is identified in the atlas from this study. The “comments” column also describes details regarding the annotation of the cell type.

**Table 2. Summary of number of cells and cell types sequenced per individual and tissue**

The first sheet (“Summary”) provides for each tissue (rows) and for all tissues combined (last row) the total number of cells, number of high quality cells (excluding cells labeled as doublets or low quality) in each individual separately and combined, mean gene count per high quality cell and standard deviation, mean sequencing read (Smart-seq2) or UMI count (10x) per high quality cell and standard deviation, and number of cell types (excluding cells labeled as mix, doublets or low quality). This information is provided for the 10x and SS2 datasets separately and combined. The subsequent sheets are labeled by tissue and provide a list of the cell types annotated in that tissue (including cells labeled as doublets, mix, hybrid, and low quality) with the corresponding assigned tissue system (organized as in Fig. 1c), compartment, and cell ontology designation, dendrogram_annotation_number, order compartment_freeannotation_tissue, order tissue_compartment_freeannotation (variables explained online at the Tabula Microcebus portal, see “Data” tab). The number of cells labeled for that cell type are provided for the 10x and SS2 datasets separately and combined, as well as for each individual separately and combined.

**Table 3. Differentially-expressed genes for each mouse lemur molecular cell type**

Four tables list the top 300 significant (adjusted *p*-value<0.05) differentially-expressed genes (DEGs) detected for each cell type in the 10x dataset (≥5 cells) using four different comparisons: vs. all other cell types in the atlas (“Table 3a DEG_atlas-wide”), vs. all other cell types from the same compartment across the atlas (“Table 3b DEG_atlas-compartment-wide”), vs. all other cell types from the same tissue (sheet “Table 3c DEG_tissue-wide”), or vs. all other cell types from the same compartment of that tissue (sheet “Table 3d DEG_tissue-compartment-wide”). For each cell type, three columns are included: gene symbol (ordered with ascending *p*-values), adjusted *p*-value, and log2-transformed fold-change of the DEGs. Columns of cell types with fewer than 5 10x cells or with no significant DEGs were left blank.

**Table 4. One-to-one orthologous genes across human, lemur, mouse**

Rows are individual genes with one-to-one orthologs across the three species, with columns showing gene type, description, as well as gene ID, symbol, and homology type assigned by the NCBI and Ensembl orthology databases for each of the species. The table also included columns showing the orthologs in rhesus and crab-eating macaques as assigned in the Ensembl orthology database. Genes without a macaque ortholog or with non one-to-one mapping were left blank in the respective macaque columns. A small number of one-to-one orthologs were curated manually as indicated in the column ‘isManualCuration’.

**Table 5. Datasets and molecular cell types analyzed across species**

The first sheet ‘datasets’ lists the datasets used in the cross-species comparison, separately for each species and tissue type, as well as metadata showing the individuals and number of high quality cells analyzed.

The second sheet ‘cell types’ lists in rows all molecular cell types analyzed in the cross-species comparisons with columns indicating their tissue group, compartment, unified cell type annotations, as well as number of cells from each species, and whether the cell types were analyzed in the 3-species (human, lemur, mouse) or 4-species (additionally including macaque comparison.

**Table 6. Genes with a species conserved, primate-enriched, or primate-depleted expression for each molecular cell type across human, lemur, and mouse**

The first sheet ‘conserved’ lists the genes that are enriched in each cell type, regardless of species. Columns indicate the gene (in human symbol), information of the respective cell type (tissue, compartment, and annotation), as well as *p*-value and fold-change comparing the expression of the cell type to the rest cells in the dataset.

The second (‘primate high’) and third (‘primate low’) sheets list the genes with a primate-enriched or primate-depleted expression, respectively, for each of the cell types. Columns indicate the gene (in human symbol), information of the respective cell type (tissue, compartment, and annotation), as well as *p*-values and fold-changes comparing both the human and lemur cells of the cell type vs. mouse cells of the homologous cell type.

**Table 7. Global patterns of gene expression conversation across human, lemur, and mouse**

Rows are individual genes/orthologs with columns showing the human gene symbol, the expression similarity scores between human and lemur (*r*_g_(HL)), human and mouse (*r*_g_(HM)), and lemur and mouse (*r*_g_(LM)), as well as three pairwise differences of these scores and their significance (*p*-value) (see Methods). Additional columns include diseases that the genes are known to be associated with, as assigned by the OMIM database, and the categorization of the gene.

**Table 8. Gene set enrichment analysis of the genes showing a highly-conserved expression pattern**.

Table exported from g:Profiler with rows listing statistically enriched gene sets and columns showing information of the gene set, the enrichment statistics, and the related highly-conserved genes.

## SUPPLEMENTAL MATERIAL

### Legends for Supplementary Figures

**Figure S1. Full dendrogram of cell type designations in mouse lemur atlas**

Expansion of Fig. 2a showing full dendrogram of the 256 assigned designations of molecular cell type across atlas. Designations are arranged by compartment (epithelial, endothelial, stromal, immune-lymphoid/hematopoietic/myeloid, neural, germ) and then ordered by tissue system (epithelial compartment) or biological relatedness (other compartments). PF, proliferative cell state; H1-12, hybrid cell types (cell clusters with gene signature of two cell types); M1-7, mixed clusters of distinct cell types too few to assign separately; (), tissue abbreviation indicated for cell types present in <4 tissues with full tissue name in black box; *, pathological cell states, lung tumor metastasized from uterus (L2) and uterine tumor (L3), as described in accompanying Tabula Microcebus manuscript 2.

**Figure S2. Dendrogram of cell types in brain cortex, brainstem, retina, hypothalamus/pituitary, and limb muscle**.

**a, b**. Dendrograms of the 23 molecular cell types identified from scRNA-seq of brain cortex (3120 10x cells, 292 SS2 cells; L1, L2, L4) (a), and the 25 molecular cell types from brainstem (1988 10x cells, 122 SS2 cells; L2, L4) (b). The cortex and brainstem samples captured 1099 neurons including excitatory (*SLC17A6+, SLC17A7+*) and inhibitory (*GAD1+, GAD2+, SLC32A1+*) neurons that with further resolved by subclustering into at least 10 major molecular types including four cortex-specific excitatory, one brainstem-specific excitatory, one cortex-specific inhibitory, three brainstem-specific inhibitory, and one hybrid population expressing both excitatory and inhibitory markers. Different glial cell types were also identified including three molecular types of astrocytes, oligodendrocytes and their precursors (OPCs), choroid plexus cells, ependymal cells, as well as surprisingly Schwann cells (myelinating and non-myelinating) which are generally associated with peripheral nervous system. In the endothelial compartment we identified vein cells as well as capillary cells (*FABP5-RBP7-*) that showed distinct gene signatures compared to capillary cells of peripheral tissues (Fig. 4d). In the stromal compartment, vascular smooth muscle cells, pericytes, and leptomeningeal cells were identified. For the immune compartment, major classes of lymphoid and myeloid cells (B, NK/T, macrophages/microglia, neutrophils, and erythroid cells) were identified.

**c**. Dendrogram of the 11 molecular cell types identified from retina scRNA-seq (938 10x cell, 120 SS2 cells, L4). Identified cells include two types of photoreceptors (dim-light monochromatic rod cells and bright-light multichromatic cones), ON- and OFF-bipolar cells^83^ that rods or cones synapse on plus a hybrid bipolar cell type, a type of horizontal neurons, two types of glial cells (Muller and non-myelinating Schwann cells), and three immune populations (macrophages/microglia, neutrophils, erythroid cells). We found that the photoreceptor cells of the mouse lemur retina is rod-dominated (99.4% rods vs. 0.6% cones in the 10x scRNA-seq unsorted sample), which fits the nocturnal behavior of the mouse lemur and is consistent with reported photoreceptor cell densities^84^. Lemurs (and new world monkeys) have only two opsins (unlike old world monkeys and apes which have three^84^), and two of the six sequenced cone cells expressed the long-wave-sensitive cone opsin (*OPN1LW*) but none expressed the short/medium-wave-opsin (*OPN1SW*). Two cell types not obtained are the ganglion cells on which bipolar cells synapse and amacrine cells which serve as inhibitory interneurons that modulate and shape signaling.

**d**. Dendrogram of the 34 molecular cell types identified from hypothalamus/pituitary scRNA-seq (2431 10x cells, L4). Identified cells include all five canonical neuroendocrine cell types of the anterior pituitary (based on expression of their corresponding hormones and other classical markers, see Table 2); interestingly, a few cells (labeled “hybrid”) expressed markers for more than one neuroendocrine cell type. We also identified a GABAergic neuron and multiple glial molecular types including oligodendrocytes, OPCs, ependymal cells, and astrocytes. In the endothelial compartment, two capillary types were identified, one (capillary *FABP5-RBP7-*) shared with brain cortex and brainstem and another (sinusoid *MAFB+*) likely the fenestrated sinusoidal capillaries of the pituitary gland. Stromal compartment cell types identified include vascular smooth muscle cells, pericytes, leptomeningeal cells, and fibroblasts, and immune cell types spanned diverse lymphoid and myeloid cells including several types of immune progenitors and proliferating cells.

**e**. Dendrogram of the 31 molecular cell types identified from limb muscle scRNA-seq (2968 10x cells, 994 SS2 cells; L2, L4). Identified cells include 10 distinct stromal molecular types as described in the main text, as well as diverse endothelial, lymphoid, and myeloid cells. One significant cell type that was not identified is Schwann cells, which cover the neuromuscular junction and the terminal axon. This might be because, as in humans, the neuromuscular junctions in mouse lemurs may be small^85^ and have few associated Schwann cells.

**f**. UMAPs of limb muscle cells (L2, 10x dataset, L2) separated by compartment. Left to right: endothelial, stromal, lymphoid, and myeloid compartments.

**Figure S3. Expression profiles of selected differentially-expressed genes along cell trajectories in Fig. 3**.

**a-f**. Heat map showing relative expression (10x dataset from lemur, tissue, and compartment indicated at top of panel) of the indicated differentially-expressed genes normalized to stable maximal value (99.5 percentile) of the gene along trajectories of Fig. 3. Genes shown are top three genes from each of the detected trajectory-dependent expression patterns described in Methods. Cells are ordered left to right by their trajectory coordinates, and their cell type designations are indicated by color in top bar (color key as in UMAPs of Fig. 3). Trajectories in panels d and e include bone and bone marrow hematopoietic precursor cells (HPC) and granulocyte/monocyte progenitors (GMP), and heat map in panel f includes bone and bone marrow erythroid progenitors and lineage cells. [], description of gene identified by NCBI as a gene locus (LOC105884179 [BEX1], LOC105877478 [CCSAP], LOC105862715 [DCUN1D1], LOC105881500 [DEFA1], LOC105865511 [FCGR1A], LOC105867419 [GSTP1], LOC105856255 [HBA1], LOC105883507 [HBB], LOC105859005 [HP], LOC105872012 [HLA-DRB1-1], LOC105859819 [HRNR], LOC105874071 [IFITM3], LOC105882927 [LRRC9], LOC105884612 [NAT8B], LOC105880511 [PTMA], LOC105883741 [SERPINB3], LOC105876721 [SERPINB3], LOC105859377 [TMEM45A], LOC105865212 [TMEM14C], LOC105871594 [RGCC], LOC105865610 [RDH7], LOC105865617 [RDH16], LOC105864771 [RNASE2], LOC105876678 [uncharacterized 1], LOC105873147 [uncharacterized 2], LOC105862290 [uncharacterized 3], LOC105858108 [uncharacterized 4], LOC105880776 [uncharacterized 5], LOC105881161 [uncharacterized 6], LOC105871650 [uncharacterized 7]).

**g-l**. Cell UMAPs from Fig. 3 color-coded by molecular trajectory coordinates calculated using the algorithm Slingshot (left) and comparison of the cells’ trajectory coordinates assigned by two independent methods (method 1, in-house algorithm, see Fig. 3; method 2: Slingshot) (right).

Red dashed lines indicate 1-1 relationship.

**m-r**. UMAPs and detected molecular trajectories of cells from the indicated tissues and compartments as in Fig. 3, but from other lemur individuals as indicated at bottom left of each UMAP.

**Figure S4. Hepatocyte molecular subtypes in mouse and lemur**

**a**. UMAP of liver hepatocytes and cholangiocytes, separately for human (left), lemur (middle), and mouse (right). Top to bottom: cells colored by cell type annotation, by sex of the animals from which cells were isolated, by heatmap showing normalized expression of a hepatocyte marker (*ASGR1*), a cholangiocyte marker (*SPP1*), and a hepatocyte subtype differentially expressed gene (*CPN2*).

**b**. Species-integrated UMAP of liver hepatocytes and cholangiocytes, with cells color-coded by the species-integrated cell type annotation (top) and species (bottom).

**c**. Box and whisker plots of the number of genes (top) and UMIs (bottom) detected per cell for each cell type indicated.

**d**. Dot plot of cross-species expression of the two hepatocyte molecular types and cholangiocytes of the indicated hepatocyte, cholangiocyte markers, differentially-expressed genes between the two hepatocyte types, liver zonation markers^86,87^, and cell stress markers including immediate-early genes and heat shock proteins. Rows are one-to-one orthologs labeled with the respective human symbol. Columns are cell types, displayed as triplets of the respective expression in humans, lemurs, and mice. Note that in all three species, hepatocyte molecular types do not correspond to expression of liver zonation markers and do not differ significantly in expression of cell stress markers.

**Figure S5. UMAP and marker genes of previously unknown cell types**

**a, b**. UMAP (a) of the bone stromal and neural cells (L2 and L4, 10x dataset), integrated by FIRM and colored according to molecular cell types. Dashed circle, unknown (*NGFR+ TNNT2+*) stromal cell type (#141). Dot plot (b) showing fraction of expressing cells and average expression in the molecular cell types in panel a for the indicated compartment/cell type marker genes and genes differentially-expressed in the unknown stromal cell type (#141) relative to all other cells from bone.

**c, d**. UMAP (c) of the tongue stromal and neural cells (L2 and L4, 10x dataset) integrated by FIRM and colored according to molecular cell types. Dashed circles, unknown stromal (*NGFR+ TNNT2+*) cell type (#141) and (*COL15A1+ PTGDS+*) cell type (#142). Dot plot (d) as above showing expression in the molecular cell types in panel c of compartment/cell type marker genes and genes differentially-expressed in the unknown stromal cell types (#141, 142) relative to all other cells from tongue.

**e**. Dot plot as above showing average expression (L1-L4, 10x dataset) of indicated marker genes and differentially-expressed genes in the unknown stromal (*NGFR+ TNNT2+*) populations (#141) from mammary gland, bone, tongue, and pancreas, and the (*COL15A1+ PTGDS+*) population (#142) found in tongue. Also included are cardiomyocytes, mesothelial cells, leptomeningeal cells, and Schwann cells from all tissues, which express high levels of the two unknown populations’ tissue-specific, differentially-expressed genes.

**f-g**. UMAP (f) of kidney stromal cells (L2, L3, and L4, 10x dataset), integrated by FIRM and colored according to molecular cell types. Dashed circle, unknown stromal (*ST6GAL2+*) cell type (#143). Dot plot (g) as above showing average expression in kidney epithelial, endothelial, and stromal cell types of indicated compartment/cell type markers and genes differentially-expressed in the unknown stromal cell type (#143) relative to all other cells from kidney.

**h, i**. UMAP (h) of epithelial and stromal cells from gonadal fat adipocyte tissue (GAT) (L2, 10x dataset), integrated by FIRM and colored according to molecular cell types. Dashed circle, unknown epithelial (*CRISP3+*) cell type (#15). Dot plot (i) as above showing average expression in the molecular cell types in panel h of indicated compartment/cell type marker genes and genes differentially-expressed in the unknown epithelial molecular cell type (#15) relative to all other cells in the GAT.

**j**. Dot plot as above showing average expression of indicated compartment/cell type marker genes (and genes differentially-expressed in epithelial (*CRISP3+*) cell type) in the epithelial (*CRISP3+*) population (#15) from GAT, as well as urothelial cells and mesothelial cells from all tissues (L1-L4, 10x dataset), which share expression of some of the genes differentially-expressed in the unknown epithelial (*CRISP3+*) population (#15). Fibroblasts are also included for comparison. [], description of gene identified by NCBI as a gene locus (LOC109730246 [uncharacterized 1]).

**k, l**. UMAP (k) of the blood cells (L2 and L4, SS2 dataset), integrated by FIRM and colored according to molecular cell types. Dashed circle, unknown epithelial (*PGAP1+*) cell type (#16). Dot plot (l) as above showing average expression of the molecular cell types in panel k of indicated compartment/cell type marker genes and genes differentially-expressed in the unknown epithelial molecular cell type (#16) relative to all other cells from blood. [], description of gene identified by NCBI as a gene locus (LOC105861777 [uncharacterized 2], LOC105884518 [TTC39B]).

**m**. Dot plot as above showing average expression of indicated compartment/cell type marker genes (and genes differentially-expressed in epithelial (*PGAP1+*) cell type) in the epithelial (*PGAP1+*) cell type (#16) from blood, as well as astrocytes, oligodendrocyte precursor cells (OPC), oligodendrocytes, and ependymal cells from all tissues (L1-L4, SS2 dataset), which express high levels of some of the epithelial (*PGAP1+*) tissue-specific, differentially-expressed genes.

**Figure S6. Relationships of molecular cell types across the mouse lemur cell atlas viewed in a heat map (a-b), and clustering of adipocyte (c) and neutrophil (d) molecular types**

**a**. Heat maps of pairwise Pearson correlation coefficients between the transcriptomic profiles of each of the 749 molecular cell type in atlas (10x and SS2 datasets, excluding cardiac cells), calculated from principal component values of FIRM-integrated UMAP (Fig. 1g) averaged across all cells of each type. Cell types ordered by compartment, then cell type designation/number, and then tissue.

**b**. Close up of heat map from panel a showing pairwise correlations between skin epithelial cells and all other cell types (top), and between testicular germ cells and all other cell types (bottom). Note proliferating skin interfollicular suprabasal cells show high correlation with proliferating and progenitor cell types across all compartments in atlas. Spermatogonia also show high correlation with proliferating cell types in the atlas, but especially with hematopoietic progenitor cells.

**c-d**. Close up of portions of UMAP in panel a showing segregation of three types of neutrophils (c) and of two types of adipocytes (d).

**Figure S7. Pairwise comparisons of the relationships of the transcriptomic profiles of all molecular cell types across the mouse lemur atlas**

Heat maps as in Fig. 5g of pairwise Pearson correlation coefficients between the transcriptomic profiles of each of the 749 molecular cell type in atlas (10x and SS2 datasets, excluding all cardiac cells because of low quality), calculated from principal component values of FIRM-integrated UMAP (Fig. 1g) averaged across all cells of each type. Cell types ordered by compartment, then cell type annotation number, then tissue (a); by tissue, then compartment, then cell type annotation number (b); and by hierarchical clustering of cell type expression profiles (c). Interactive forms of the heat maps are available at the Tabula Microcebus portal.

**Figure S8. Molecular relationship of cell types within- and cross-compartments in human, lemur, and mouse**

**a-c**. UMAPs of the 63 orthologous cell types (dots) analyzed across-species (see Section 6), separately in human (a), lemur (b), and mouse (c). Note the close clustering of the germline progenitors with the immune progenitors, as in Fig. 5a.

**d**. Violin plot showing transcriptomic distances of cell types within the same compartment and cross different compartments, in human, lemur, and mouse.

**e**. Dot plot showing cross-species expression patterns for the differentially expressed genes detected by comparing the mouse lemur immune progenitor/proliferating cells and germ cells vs. cell types in other compartments, or vs. proliferating cells in other compartments, as in Fig. 5c-d. Shown here are expressions in the 63 orthologous cell types analyzed across all three species. Figure formatted as in Fig. 6k.

**Figure S9. Molecular relationship of human, lemur, and mouse lung and skeletal muscle cell types**

**a, c**. UMAP of the lung (a) and skeletal muscle (c) cells integrated across species by Portal based on the one-to-one gene orthologs. In a, panels are color coded by cell type annotations separately for non-immune (left) and immune (middle), and by species (right). In c, panels are color coded by cell type annotations (left) and by species (right).

**b, d**. Sankey plot showing the relationship between human, mouse lemur, and mouse cell types for lung (a) and for skeletal muscle (c) as determined by SAMap algorithm^66^ (see Methods).

Each cell type in lemur is connected (gray line) to the cell types it maps to in human and mouse datasets; thickness of line indicates similarity score (0-1) between connected cell types. A cell type with no connecting lines indicates it did not map with similarity score > 0.1 to any cell type in the other species. Note that cell types of the same designation show higher similarity scores across species compared to other cell types.

**Figure S10. Evolutionary comparisons of spermatogenesis (a-d) and hematopoiesis (f-o)**

**a-b**. Species-integrated UMAPs of the male germ cells as in Fig. 6c, but with cells color coded by spermatogenesis stage (a) or pseudotime along the trajectory (b).

**c-d**. Dot plot showing average expression across different stages of the spermatogenesis trajectory for spermatogenesis markers and regulators (c) or the indicated evolutionary conserved/divergent genes (top, all-conserved; middle, primate-conserved and mouse deviated; bottom, lemur specific and conserved in other species). Rows are orthologous genes, indicated with the respective human gene symbols. Columns are cell types, displayed as quadruplets of the respective expression in human, macaque, lemur, and mouse.

**f-i**. Species-integrated UMAPs of the bone marrow and spleen immune cells as in Fig. 6e, but with cells color coded by cell type annotation (f) or pseudotime along each of the trajectories (g-i).

**j-k**. Correlation coefficients of the transcriptomic profiles of the human immune cell and progenitors to those of lemur (*r*_c_(HL)) and mouse (*r*_c_(HM)) across different stages of the monocyte/macrophage (j) and erythroid (k) trajectories, based on the one-to-one orthologs across the three species. Note that *r*_c_(HL) is almost always greater than *r*_c_(HM) through the trajectories, despite the end of the monocyte/macrophage trajectory which likely results from the different fractions of bone marrow and spleen macrophages in the human and lemur datasets.

**l-o**. Dot plot showing average expression of marker genes across different stages of the neutrophil (l), monocyte/macrophage (m), and erythroid (n) trajectories, and the indicated evolutionary conserved/divergent genes (top, all-conserved; bottom, primate-conserved and mouse deviated) for the neutrophil trajectory (o). Rows are orthologous genes, indicated with the respective human gene symbols. Columns are cell types, displayed as triplets of the respective expression in human, lemur, and mouse.

**Figure S11. Evolutionary comparison of transcriptomic profiles of lung cell types across human, lemur, mouse, and macaque**.

**a**. Bar plots showing human-to-macaque, human-to-lemur, and human-to-mouse correlation coefficients (*r*_c_) and their differences (Δ*r*_c_) for each of the 18 orthologous lung cell types across the four species. *p*-value calculated using one-tailed *t* test (see Methods).

**b**. Dot plot showing average expression patterns of the highly-conserved genes detected across human, lemur, and mouse (as in Fig. 6k and Fig. 13a) in the 18 orthologous lung cell types across the four species. Note that macaque generally showed a similar expression pattern as in the other three species, but there were exceptions like those indicated with arrowheads. Rows are one-to-one orthologous genes across the species, indicated with the respective human gene symbols. Columns are cell types, displayed as quadruplets of the respective expression in the four species, separated by vertical gray lines. Genes shown in Fig. 13a but lacked a one-to-one ortholog across all four species were not included.

**Figure S12. Evolutionary comparison of gene expression and sequence conservation**.

**a-c**. Scatter plots comparing gene expression conservation between human and lemur (*r*_g_(HL)), human and mouse (*r*_g_(HM)), and lemur and mouse (*r*_g_(LM)), for individual one-to-one orthologs across the three species. Gray dots are different genes with the contours showing their probability density. Black dashed line indicates 1-1 relationship. Expression of the highlight genes are shown in Fig. 6k and Fig. S13.

**d**. Ratio of HL, HM, and LM genes detected at different Δ*r*_g_ thresholds. Supporting the lemur as a genetic intermediates between human and mouse, the numbers of HL and LM genes were consistently higher than (more than doubling) that of HM genes, whereas the numbers of HL and LM genes were more comparable.

**e**. Scatter plots comparing in gene expression conservation (*r*_g_) vs. gene sequence identity (*I*) between human and lemur (HL) and between human and mouse (HM), for individual one-to-one orthologs across the three species. Note the lack of positive correlation between two measurements (Pearson’s *r*=-0.14).

**e**. Scatter plots comparing HL-HM differences in expression conservation (Δ*r*_g_) vs. gene sequence identity (Δ*I*), for individual one-to-one orthologs across the three species. Note the lack of positive correlation between two measurements (Pearson’s *r*=0.002).

**Figure S13. Expression patterns of evolutionarily conserved (a) or different types of evolutionarily divergent genes (b-f) across human, lemur, and mouse**.

**a-f**. Dot plots in the same format as in Fig. 6k showing highly species-conserved genes (a), HL genes (b) whose expression pattern was similar in the two primates but diverged in mouse thus may underlie primate-specific biology, HM (c) and LM (d) genes which may underlie lemur- and human-specific biology, respectively, as well as genes that show an overall moderately-conserved (e) and lowly-conserved (f) expression pattern across the three species. See also Fig. S12a-c.

## REFERENCES

1. Elsea, S. H. & Lucas, R. E. The mousetrap: what we can learn when the mouse model does not mimic the human disease. ILAR J. 43, 66–79 (2002).

2. Mestas, J. & Hughes, C. C. Of mice and not men: differences between mouse and human immunology. J. Immunol. 172, 2731–2738 (2004).

3. Watase, K. & Zoghbi, H. Y. Modelling brain diseases in mice: the challenges of design and analysis. Nat. Rev. Genet. 4, 296–307 (2003).

4. IUCN. IUCN Red List of Threatened Species https://www.iucnredlist.org/en (2022).

5. Ezran, C. et al. The Mouse Lemur, a Genetic Model Organism for Primate Biology, Behavior, and Health. Genetics 206, 651–664 (2017).

6. Hozer, C., Pifferi, F., Aujard, F. & Perret, M. The Biological Clock in Gray Mouse Lemur: Adaptive, Evolutionary and Aging Considerations in an Emerging Non-human Primate Model. Front. Physiol. 10, 1033 (2019).

7. Perret, M. & Aujard, F. Daily hypothermia and torpor in a tropical primate: synchronization by 24-h light-dark cycle. Am. J. Physiol. Regul. Integr. Comp. Physiol. 281, R1925–33 (2001).

8. Génin, F., Schilling, A. & Perret, M. Social inhibition of seasonal fattening in wild and captive gray mouse lemurs. Physiol. Behav. 86, 185–194 (2005).

9. Kraska, A. et al. Age-associated cerebral atrophy in mouse lemur primates. Neurobiol. Aging 32, 894–906 (2011).

10. Terrien, J. et al. Metabolic and genomic adaptations to winter fattening in a primate species, the grey mouse lemur (Microcebus murinus). Int. J. Obes. 42, 221–230 (2018).

11. Languille, S. et al. The grey mouse lemur: a non-human primate model for ageing studies. Ageing Res. Rev. 11, 150–162 (2012).

12. Radespiel, U. Sociality in the gray mouse lemur (Microcebus murinus) in northwestern Madagascar. Am. J. Primatol. 51, 21–40 (2000).

13. Yoder, A. D. et al. Remarkable species diversity in Malagasy mouse lemurs (primates, Microcebus). Proc. Natl. Acad. Sci. U. S. A. 97, 11325–11330 (2000).

14. Rina Evasoa, M. et al. Variation in reproduction of the smallest-bodied primate radiation, the mouse lemurs (Microcebus spp.): A synopsis. Am. J. Primatol. 80, e22874 (2018).

15. Atsalis, S. A Natural History of the Brown Mouse Lemur. vol. 6 (Prentice Hall, 2015).

16. Larsen, P. A. et al. Hybrid de novo genome assembly and centromere characterization of the gray mouse lemur (Microcebus murinus). BMC Biol. 15, 110 (2017).

17. Jain, M. et al. Nanopore sequencing and assembly of a human genome with ultra-long reads. Nat. Biotechnol. 36, 338–345 (2018).

18. Wu, A. R. et al. Quantitative assessment of single-cell RNA-sequencing methods. Nat. Methods 11, 41–46 (2013).

19. The Tabula Muris Consortium. Single-cell transcriptomics of 20 mouse organs creates a Tabula Muris. Nature 562, 367–372 (2018).

20. The Tabula Muris Consortium. A single-cell transcriptomic atlas characterizes ageing tissues in the mouse. Nature 583, 590–595 (2020).

21. Travaglini, K. J. et al. A molecular cell atlas of the human lung from single-cell RNA sequencing. Nature 587, 619–625 (2020).

22. Casey, K. M., Karanewsky, C. J., Pendleton, J. L., Krasnow, M. R. & Albertelli, M. A. Fibrous Osteodystrophy, Chronic Renal Disease, and Uterine Adenocarcinoma in Aged Gray Mouse Lemurs (Microcebus murinus). Comp. Med. 71, 256–266 (2021).

23. The Tabula Microcebus Consortium. Mouse lemur transcriptomic atlas elucidates primate genes, physiology, disease, and evolution. bioRxiv 2022.08.06.503035 (2022) doi:10.1101/2022.08.06.503035.

24. Butler, A., Hoffman, P., Smibert, P., Papalexi, E. & Satija, R. Integrating single-cell transcriptomic data across different conditions, technologies, and species. Nat. Biotechnol. 36, 411–420 (2018).

25. Larsson, A. J. M., Stanley, G., Sinha, R., Weissman, I. L. & Sandberg, R. Computational correction of index switching in multiplexed sequencing libraries. Nat. Methods 15, 305–307 (2018).

26. Ming, J. et al. FIRM: Flexible integration of single-cell RNA-sequencing data for large-scale multitissue cell atlas datasets. Brief. Bioinform. 23, bbac167 (2022).

27. Diehl, A. D. et al. The Cell Ontology 2016: enhanced content, modularization, and ontology interoperability. J. Biomed. Semantics 7, 44 (2016).

28. Pisani, D. F., Bottema, C. D. K., Butori, C., Dani, C. & Dechesne, C. A. Mouse model of skeletal muscle adiposity: a glycerol treatment approach. Biochem. Biophys. Res. Commun. 396, 767–773 (2010).

29. Tuttle, L. J., Sinacore, D. R. & Mueller, M. J. Intermuscular adipose tissue is muscle specific and associated with poor functional performance. J. Aging Res. 2012, 172957 (2012).

30. Alexander, M. S. et al. CD82 Is a Marker for Prospective Isolation of Human Muscle Satellite Cells and Is Linked to Muscular Dystrophies. Cell Stem Cell 19, 800–807 (2016).

31. Xu, X. et al. Human Satellite Cell Transplantation and Regeneration from Diverse Skeletal Muscles. Stem Cell Reports 5, 419–434 (2015).

32. Liu, L., Cheung, T. H., Charville, G. W. & Rando, T. A. Isolation of skeletal muscle stem cells by fluorescence-activated cell sorting. Nat. Protoc. 10, 1612–1624 (2015).

33. Sacco, A., Doyonnas, R., Kraft, P., Vitorovic, S. & Blau, H. M. Self-renewal and expansion of single transplanted muscle stem cells. Nature 456, 502–506 (2008).

34. Joe, A. W. B. et al. Muscle injury activates resident fibro/adipogenic progenitors that facilitate myogenesis. Nat. Cell Biol. 12, 153–163 (2010).

35. Uezumi, A., Fukada, S.-I., Yamamoto, N., Takeda, S. ‘ichi & Tsuchida, K. Mesenchymal progenitors distinct from satellite cells contribute to ectopic fat cell formation in skeletal muscle. Nat. Cell Biol. 12, 143–152 (2010).

36. Wosczyna, M. N. et al. Mesenchymal Stromal Cells Are Required for Regeneration and Homeostatic Maintenance of Skeletal Muscle. Cell Rep. 27, 2029–2035.e5 (2019).

37. Uezumi, A. et al. Cell-Surface Protein Profiling Identifies Distinctive Markers of Progenitor Cells in Human Skeletal Muscle. Stem Cell Reports 7, 263–278 (2016).

38. Italiani, P. & Boraschi, D. From Monocytes to M1/M2 Macrophages: Phenotypical vs. Functional Differentiation. Front. Immunol. 5, 514 (2014).

39. Peti-Peterdi, J. & Harris, R. C. Macula densa sensing and signaling mechanisms of renin release. J. Am. Soc. Nephrol. 21, 1093–1096 (2010).

40. Liu, S. et al. An organism-wide atlas of hormonal signaling based on the mouse lemur single-cell transcriptome. Nat. Commun. 15, 2188 (2024).

41. Andrès, M., Solignac, M. & Perret, M. Mating system in mouse lemurs: theories and facts, using analysis of paternity. Folia Primatol. (Basel) 74, 355–366 (2003).

42. Martin, M. D. & Badovinac, V. P. Defining Memory CD8 T Cell. Front. Immunol. 9, 2692 (2018).

43. Farber, D. L., Yudanin, N. A. & Restifo, N. P. Human memory T cells: generation, compartmentalization and homeostasis. Nat. Rev. Immunol. 14, 24–35 (2014).

44. Ben-Moshe, S. & Itzkovitz, S. Spatial heterogeneity in the mammalian liver. Nat. Rev. Gastroenterol. Hepatol. 16, 395–410 (2019).

45. Celton-Morizur, S. & Desdouets, C. Polyploidization of liver cells. Adv. Exp. Med. Biol. 676, 123– 135 (2010).

46. Tabula Sapiens Consortium. The Tabula Sapiens: A multiple-organ, single-cell transcriptomic atlas of humans. Science 376, eabl4896 (2022).

47. Coppiello, G. et al. Meox2/Tcf15 heterodimers program the heart capillary endothelium for cardiac fatty acid uptake. Circulation 131, 815–826 (2015).

48. Iso, T. et al. Capillary endothelial fatty acid binding proteins 4 and 5 play a critical role in fatty acid uptake in heart and skeletal muscle. Arterioscler. Thromb. Vasc. Biol. 33, 2549–2557 (2013).

49. Caprioli, A., Zhu, H. & Sato, T. N. CRBP-III:lacZ expression pattern reveals a novel heterogeneity of vascular endothelial cells. Genesis 40, 139–145 (2004).

50. Hu, C. et al. Retinol-binding protein 7 is an endothelium-specific PPAR cofactor mediating an antioxidant response through adiponectin. JCI Insight 2, e91738 (2017).

51. Du, Y. et al. Single-Cell Transcriptome Atlas of Murine Endothelial Cells. Cell 180, 764–779.e20 (2020).

52. Takeda, A. et al. Single-Cell Survey of Human Lymphatics Unveils Marked Endothelial Cell Heterogeneity and Mechanisms of Homing for Neutrophils. Immunity 51, 561–572.e5 (2019).

53. Fujimoto, N. et al. Single-cell mapping reveals new markers and functions of lymphatic endothelial cells in lymph nodes. PLoS Biol. 18, e3000704 (2020).

54. Jalkanen, S. & Salmi, M. Lymphatic endothelial cells of the lymph node. Nat. Rev. Immunol. 20, 566–578 (2020).

55. Neumann, J. T. et al. Application of High-Sensitivity Troponin in Suspected Myocardial Infarction. N. Engl. J. Med. 380, 2529–2540 (2019).

56. Jarolim, P. High sensitivity cardiac troponin assays in the clinical laboratories. Clin. Chem. Lab. Med. 53, 635–652 (2015).

57. Zipes, D. P., Libby, P., Bonow, R. O., Mann, D. L. & Tomaselli, G. F. Braunwald’s Heart Disease E-Book: A Textbook of Cardiovascular Medicine. (Elsevier Health Sciences, 2018).

58. He, B. et al. Single-cell RNA sequencing reveals the mesangial identity and species diversity of glomerular cell transcriptomes. Nat. Commun. 12, 2141 (2021).

59. Gillich, A. et al. Capillary cell-type specialization in the alveolus. Nature 586, (2020).

60. Plasschaert, L. W. et al. A single-cell atlas of the airway epithelium reveals the CFTR-rich pulmonary ionocyte. Nature 560, 377–381 (2018).

61. Montoro, D. T. et al. A revised airway epithelial hierarchy includes CFTR-expressing ionocytes. Nature 560, 319–324 (2018).

62. Shami, A. N. et al. Single-Cell RNA Sequencing of Human, Macaque, and Mouse Testes Uncovers Conserved and Divergent Features of Mammalian Spermatogenesis. Dev. Cell 54, 529–547.e12 (2020).

63. Ernst, C., Eling, N., Martinez-Jimenez, C. P., Marioni, J. C. & Odom, D. T. Staged developmental mapping and X chromosome transcriptional dynamics during mouse spermatogenesis. Nat. Commun. 10, 1–20 (2019).

64. Qu, J. et al. A reference single-cell regulomic and transcriptomic map of cynomolgus monkeys. Nat. Commun. 13, 4069 (2022).

65. Zhao, J. et al. Adversarial domain translation networks for integrating large-scale atlas-level single-cell datasets. Nature Computational Science 2, 317–330 (2022).

66. Tarashansky, A. J. et al. Mapping single-cell atlases throughout Metazoa unravels cell type evolution. Elife 10, e66747 (2021).

67. Antigny, F. et al. Potassium Channel Subfamily K Member 3 (KCNK3) Contributes to the Development of Pulmonary Arterial Hypertension. Circulation 133, 1371–1385 (2016).

68. von Kopylow, K. & Spiess, A. N. Human spermatogonial markers. Stem Cell Res. 25, 300–309 (2017).

69. Kobayashi, K. et al. Prss55 but not Prss51 is required for male fertility in mice†. Biol. Reprod. 103, 223–234 (2020).

70. Lin, M. et al. Matrix Metalloproteinase-8 Facilitates Neutrophil Migration through the Corneal Stromal Matrix by Collagen Degradation and Production of the Chemotactic Peptide Pro-Gly-Pro. Am. J. Pathol. 173, 144 (2008).

71. Han, L. et al. Cell transcriptomic atlas of the non-human primate Macaca fascicularis. Nature 604, 723–731 (2022).

72. Bakken, T. E. et al. Comparative cellular analysis of motor cortex in human, marmoset and mouse. Nature 598, 111–119 (2021).

73. Hilton, H. G. et al. Single-cell transcriptomics of the naked mole-rat reveals unexpected features of mammalian immunity. PLoS Biol. 17, e3000528 (2019).

74. Cao, C. et al. Comprehensive single-cell transcriptome lineages of a proto-vertebrate. Nature 571, 349–354 (2019).

75. Sofroniew, N. et al. Napari: A Multi-Dimensional Image Viewer for Python. (Zenodo, 2022). doi:10.5281/ZENODO.3555620.

76. Farouni, R., Djambazian, H., Ferri, L. E., Ragoussis, J. & Najafabadi, H. S. Model-based analysis of sample index hopping reveals its widespread artifacts in multiplexed single-cell RNA-sequencing. Nat. Commun. 11, 2704 (2020).

77. Griffiths, J. A., Richard, A. C., Bach, K., Lun, A. T. L. & Marioni, J. C. Detection and removal of barcode swapping in single-cell RNA-seq data. Nat. Commun. 9, 2667 (2018).

78. Megill, C. et al. cellxgene: a performant, scalable exploration platform for high dimensional sparse matrices. bioRxiv (2021) doi:10.1101/2021.04.05.438318.

79. Street, K. et al. Slingshot: cell lineage and pseudotime inference for single-cell transcriptomics. BMC Genomics 19, 477 (2018).

80. Tarashansky, A. J., Xue, Y., Li, P., Quake, S. R. & Wang, B. Self-assembling manifolds in single-cell RNA sequencing data. Elife 8, e48994 (2019).

81. Kolberg, L. et al. g:Profiler—interoperable web service for functional enrichment analysis and gene identifier mapping (2023 update). Nucleic Acids Res. 51, W207–W212 (2023).

82. Fisher, R. A. Frequency Distribution of the Values of the Correlation Coefficient in Samples from an Indefinitely Large Population. Biometrika 10, 507–521 (1915).

83. Shekhar, K. et al. Comprehensive Classification of Retinal Bipolar Neurons by Single-Cell Transcriptomics. Cell 166, 1308–1323 (2016).

84. Peichl, L. et al. Diversity of photoreceptor arrangements in nocturnal, cathemeral and diurnal Malagasy lemurs. J. Comp. Neurol. 527, 13–37 (2019).

85. Boehm, I. et al. Comparative anatomy of the mammalian neuromuscular junction. J. Anat. 237, 827– 836 (2020).

86. Halpern, K. B. et al. Erratum: Single-cell spatial reconstruction reveals global division of labour in the mammalian liver. Nature 543, 742 (2017).

87. Aizarani, N. et al. A human liver cell atlas reveals heterogeneity and epithelial progenitors. Nature 572, 199–204 (2019).

