## Supplementary figures for "Tabula Microcebus: A transcriptomic cell atlas of mouse lemur, an emerging primate model organism"

Fig. S1

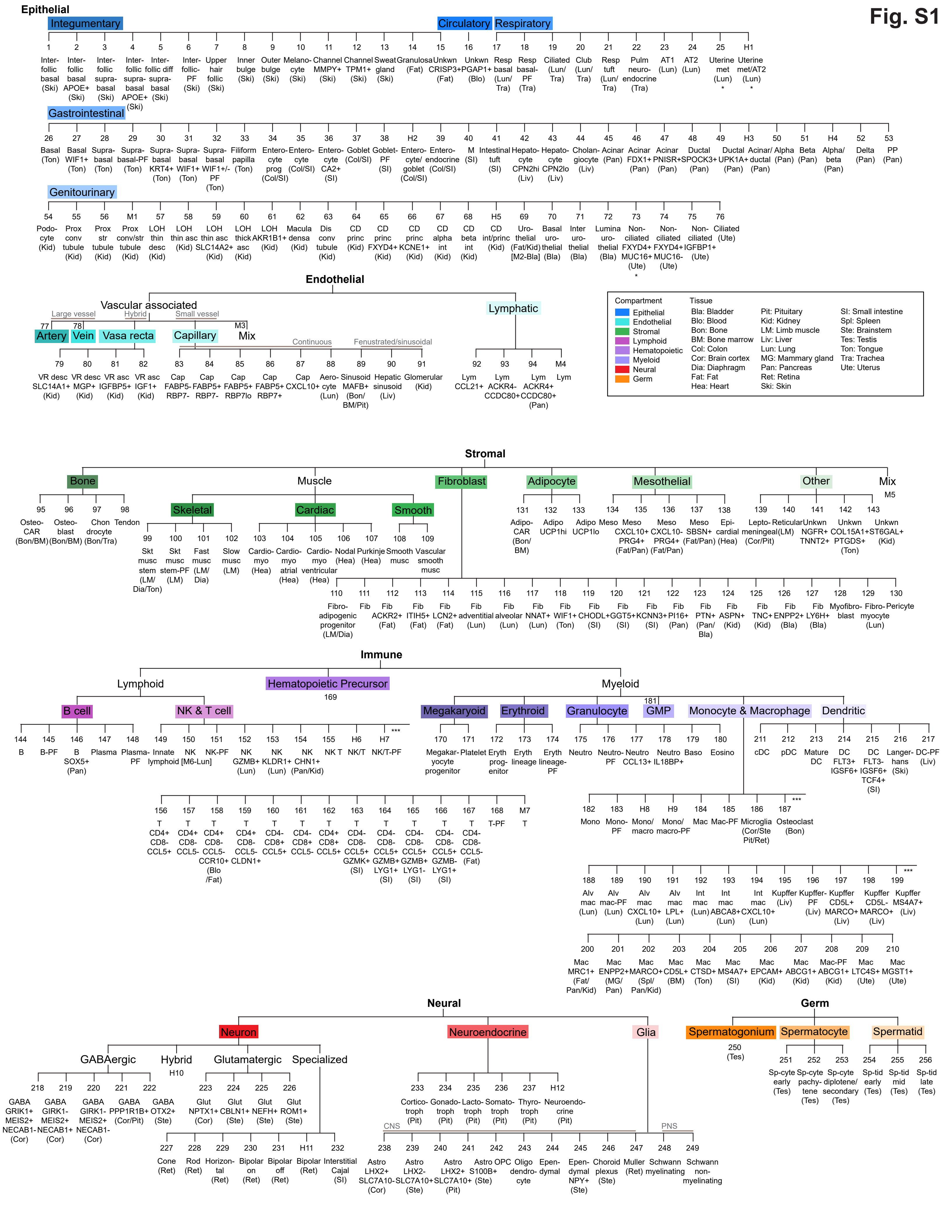

Epithelial
Integumentary
Circulatory
Respiratory
1 Inter-follic basal (Ski)
2 Inter-follic basal APOE+ (Ski)
3 Inter-follic supra-basal (Ski)
4 Inter-follic supra-basal APOE+ (Ski)
5 Inter-follic diff supra-basal (Ski)
6 Inter-follic-PF (Ski)
7 Upper hair follic (Ski)
8 Inner bulge (Ski)
9 Outer bulge (Ski)
10 Melano-cyte (Ski)
11 Channel MMPY+ (Ski)
12 Channel TPM1+ (Ski)
13 Sweat gland (Ski)
14 Granulosa (Fat)
15 Unkwn CRISP3+ (Fat)
16 Unkwn PGAP1+ (Blo)
17 Resp basal (Lun/Tra)
18 Resp basal-PF (Tra)
19 Ciliated (Lun/Tra)
20 Club (Lun/Tra)
21 Resp tuft (Lun/Tra)
22 Pulm neuro-endocrine (Tra)
23 AT1 (Lun)
24 AT2 (Lun)
25 Uterine met (Lun) *
H1 Uterine met/AT2 (Lun) *
Gastrointestinal
26 Basal (Ton)
27 Basal WIF1+ (Ton)
28 Supra-basal (Ton)
29 Supra-basal-PF (Ton)
30 Supra-basal KRT4+ (Ton)
31 Supra-basal WIF1+ (Ton)
32 Supra-basal WIF1+/- PF (Ton)
33 Filiform papilla (Ton)
34 Entero-cyte prog (Col/SI)
35 Entero-cyte (Col/SI)
36 Entero-cyte CA2+ (SI)
37 Goblet (Col/SI)
38 Goblet-PF (SI)
H2 Entero-cyte/ goblet (Col/SI)
39 Entero-endocrine (Col/SI)
40 M (SI)
41 Intestinal tuft (SI)
42 Hepato-cyte CPN2hi (Liv)
43 Hepato-cyte CPN2lo (Liv)
44 Cholan-giocyte (Liv)
45 Acinar (Pan)
46 Acinar FDX1+ (Pan)
47 Acinar PNISR+ (Pan)
48 Ductal SPOCK3+ (Pan)
49 Ductal UPK1A+ (Pan)
H3 Acinar/ ductal (Pan)
50 Alpha (Pan)
51 Beta (Pan)
H4 Alpha/ beta (Pan)
52 Delta (Pan)
53 PP (Pan)
Genitourinary
54 Podo-cyte (Kid)
55 Prox conv tubule (Kid)
56 Prox str tubule (Kid)
M1 Prox conv/str tubule (Kid)
57 LOH thin desc (Kid)
58 LOH thin asc (Kid)
59 LOH thin asc SLC14A2+ (Kid)
60 LOH thick asc (Kid)
61 LOH AKR1B1+ (Kid)
62 Macula densa (Kid)
63 Dis conv tubule (Kid)
64 CD princ (Kid)
65 CD princ FXYD4+ (Kid)
66 CD princ KCNE1+ (Kid)
67 CD alpha int (Kid)
68 CD beta int (Kid)
H5 CD int/princ (Kid)
69 Uro-thelial (Fat/Kid) [M2-Bla]
70 Basal uro-thelial (Bla)
71 Inter uro-thelial (Bla)
72 Lumina uro-thelial (Bla)
73 Non-ciliated FXYD4+ MUC16+ (Ute) *
74 Non-ciliated FXYD4+ MUC16- (Ute)
75 Non-ciliated IGFBP1+ (Ute)
76 Ciliated (Ute)
Endothelial
Vascular associated
Large vessel
Hybrid
Small vessel
77 Artery
78 Vein
Vasa recta
Capillary
M3 Mix
Continuous
Fenustrated/sinusoidal
Lymphatic
79 VR desc SLC14A1+ (Kid)
80 VR desc MGP+ (Kid)
81 VR asc IGFBP5+ (Kid)
82 VR asc IGF1+ (Kid)
83 Cap FABP5- RBP7-
84 Cap FABP5+ RBP7-
85 Cap FABP5+ RBP7lo
86 Cap FABP5+ RBP7+
87 Cap CXCL10+
88 Aero-cyte (Lun)
89 Sinusoid MAFB+ (Bon/BM/Pit)
90 Hepatic sinusoid (Liv)
91 Glomerular (Kid)
92 Lym CCL21+
93 Lym ACKR4- CCDC80+
94 Lym ACKR4+ CCDC80+ (Pan)
M4 Lym
Compartment
Epithelial
Endothelial
Stromal
Lymphoid
Hematopoietic
Myeloid
Neural
Germ
Tissue
Bla: Bladder
Blo: Blood
Bon: Bone
BM: Bone marrow
Col: Colon
Cor: Brain cortex
Dia: Diaphragm
Fat: Fat
Hea: Heart
Pit: Pituitary
Kid: Kidney
LM: Limb muscle
Liv: Liver
Lun: Lung
MG: Mammary gland
Pan: Pancreas
Ret: Retina
Ski: Skin
SI: Small intestine
Spl: Spleen
Ste: Brainstem
Tes: Testis
Ton: Tongue
Tra: Trachea
Ute: Uterus
Stromal
Bone
Muscle
Fibroblast
Adipocyte
Mesothelial
Other
Mix
M5
95 Osteo-CAR (Bon/BM)
96 Osteo-blast (Bon/BM)
97 Chon drocyte (Bon/Tra)
98 Tendon
Skeletal
Cardiac
Smooth
99 Skt musc stem (LM/Dia/Ton)
100 Skt musc stem-PF (LM)
101 Fast musc (LM/Dia)
102 Slow musc (LM)
103 Cardio-myo (Hea)
104 Cardio-myo atrial (Hea)
105 Cardio-myo ventricular (Hea)
106 Nodal (Hea)
107 Purkinje (Hea)
108 Smooth musc
109 Vascular smooth musc
131 Adipo-CAR (Bon/BM)
132 Adipo UCP1hi
133 Adipo UCP1lo
134 Meso
135 Meso CXCL10+ PRG4+ (Fat/Pan)
136 Meso CXCL10- PRG4+ (Fat/Pan)
137 Meso SBSN+ (Fat/Pan)
138 Epi-cardial (Hea)
139 Lepto-meningeal (Cor/Pit)
140 Reticular (LM)
141 Unkwn NGFR+ TNNT2+
142 Unkwn COL15A1+ PTGDS+ (Ton)
143 Unkwn ST6GAL+ (Kid)
110 Fibro-adipogenic progenitor (LM/Dia)
111 Fib
112 Fib ACKR2+ (Fat)
113 Fib ITIH5+ (Fat)
114 Fib LCN2+ (Fat)
115 Fib adventitial (Lun)
116 Fib alveolar (Lun)
117 Fib NNAT+ (Lun)
118 Fib WIF1+ (Ton)
119 Fib CHODL+ (SI)
120 Fib GGT5+ (SI)
121 Fib KCNN3+ (SI)
122 Fib PI16+ (Pan)
123 Fib PTN+ (Pan/Bla)
124 Fib ASPN+ (Kid)
125 Fib TNC+ (Kid)
126 Fib ENPP2+ (Bla)
127 Fib LY6H+ (Bla)
128 Myofibro-blast
129 Fibro-myocyte
130 Pericyte (Lun)
Immune
Lymphoid
Hematopoietic Precursor
169
Myeloid
B cell
NK & T cell
Megakaryoid
Erythroid
Granulocyte
181 GMP
Monocyte & Macrophage
Dendritic
144 B
145 B-PF
146 B SOX5+ (Pan)
147 Plasma
148 Plasma-PF
149 Innate lymphoid [M6-Lun]
150 NK
151 NK-PF
152 NK GZMB+ (Lun)
153 NK KLDR1+ (Lun)
154 NK CHN1+ (Pan/Kid)
155 NK T
H6 NK/T
H7 NK/T-PF
***
170 Megakar-yocyte progenitor
171 Platelet
172 Eryth prog-enitor
173 Eryth lineage
174 Eryth lineage-PF
175 Neutro
176 Neutro-PF
177 Neutro CCL13+
178 Neutro IL18BP+
179 Baso
180 Eosino
211 cDC
212 pDC
213 Mature DC
214 DC FLT3+ IGSF6+
215 DC FLT3- IGSF6+ TCF4+ (SI)
216 Langer-hans (Ski)
217 DC-PF (Liv)
156 T CD4+ CD8- CCL5+
157 T CD4+ CD8- CCL5-
158 T CD4+ CD8- CCL5- CCR10+ (Blo/Fat)
159 T CD4+ CD8- CCL5- CLDN1+
160 T CD4- CD8+ CCL5+
161 T CD4- CD8+ CCL5-
162 T CD4+ CD8+ CCL5+
163 T CD4- CD8- CCL5+ GZMK+ (SI)
164 T CD4- CD8- CCL5+ GZMB+ LYG1+ (SI)
165 T CD4- CD8- CCL5+ GZMB+ LYG1- (SI)
166 T CD4- CD8- CCL5+ GZMB- LYG1+ (SI)
167 T CD4- CD8- CCL5- (Fat)
168 T-PF
M7 T
182 Mono
183 Mono-PF
H8 Mono/ macro
H9 Mono/ macro-PF
184 Mac
185 Mac-PF
186 Microglia (Cor/Ste Pit/Ret)
187 Osteoclast (Bon)
***
188 Alv mac (Lun)
189 Alv mac-PF (Lun)
190 Alv mac CXCL10+ (Lun)
191 Alv mac LPL+ (Lun)
192 Int mac (Lun)
193 Int mac ABCA8+ (Lun)
194 Int mac CXCL10+ (Lun)
195 Kupffer (Liv)
196 Kupffer-PF (Liv)
197 Kupffer CD5L+ MARCO+ (Liv)
198 Kupffer CD5L- MARCO+ (Liv)
199 Kupffer MS4A7+ (Liv)
***
200 Mac MRC1+ (Fat/Pan/Kid)
201 Mac ENPP2+ (MG/Pan)
202 Mac MARCO+ (Spl/Pan/Kid)
203 Mac CD5L+ (BM)
204 Mac CTSD+ (Ton)
205 Mac MS4A7+ (SI)
206 Mac EPCAM+ (Kid)
207 Mac ABCG1+ (Kid)
208 Mac-PF ABCG1+ (Kid)
209 Mac LTC4S+ (Ute)
210 Mac MGST1+ (Ute)
Neural
Neuron
Neuroendocrine
Glia
GABAergic
Hybrid
H10
Glutamatergic
Specialized
218 GABA GRIK1+ MEIS2+ NECAB1- (Cor)
219 GABA GIRK1- MEIS2+ NECAB1+ (Cor)
220 GABA GIRK1- MEIS2+ NECAB1- (Cor)
221 GABA PPP1R1B+ (Cor/Pit)
222 GABA OTX2+ (Ste)
223 Glut NPTX1+ (Cor)
224 Glut CBLN1+ (Ste)
225 Glut NEFH+ (Ste)
226 Glut ROM1+ (Ste)
227 Cone (Ret)
228 Rod (Ret)
229 Horizon-tal (Ret)
230 Bipolar on (Ret)
231 Bipolar off (Ret)
H11 Bipolar (Ret)
232 Interstitial Cajal (SI)
233 Cortico-troph (Pit)
234 Gonado-troph (Pit)
235 Lacto-troph (Pit)
236 Somato-troph (Pit)
237 Thyro-troph (Pit)
H12 Neuroendo-crine (Pit)
CNS
PNS
238 Astro LHX2+ SLC7A10- (Cor)
239 Astro LHX2- SLC7A10+ (Ste)
240 Astro LHX2+ SLC7A10+ (Pit)
241 Astro S100B+ (Ste)
242 OPC
243 Oligo dendro-cyte
244 Epen-dymal
245 Epen-dymal NPY+ (Ste)
246 Choroid plexus (Ste)
247 Muller (Ret)
248 Schwann myelinating
249 Schwann non-myelinating
Germ
Spermatogonium
Spermatocyte
Spermatid
250 (Tes)
251 Sp-cyte early (Tes)
252 Sp-cyte pachy-tene (Tes)
253 Sp-cyte diplotene/ secondary (Tes)
254 Sp-tid early (Tes)
255 Sp-tid mid (Tes)
256 Sp-tid late (Tes)

### Fig. S2

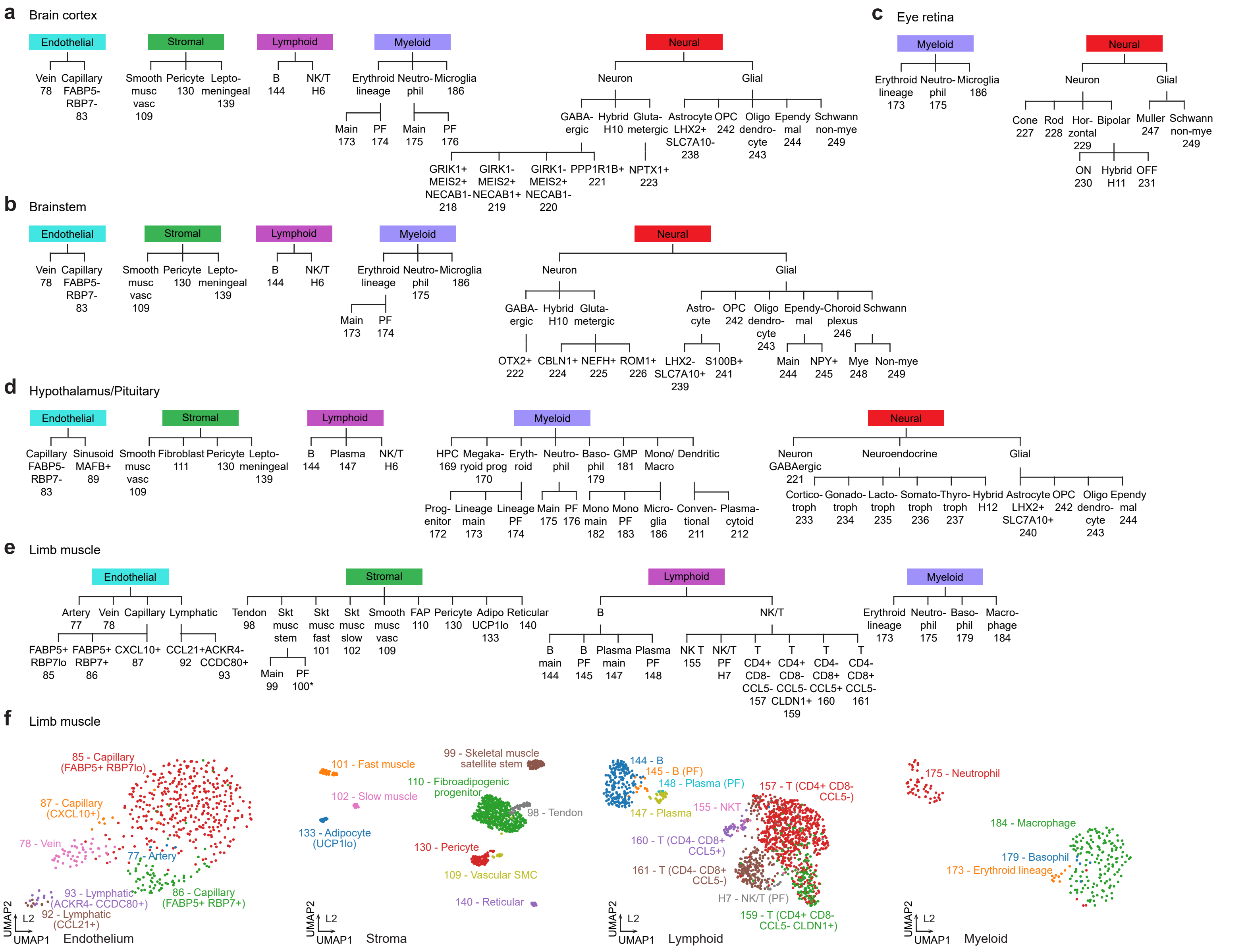

a Brain cortex
Endothelial
Stromal
Lymphoid
Myeloid
Neural
Vein 78
Capillary FABP5- RBP7- 83
Smooth musc vasc 109
Pericyte 130
Lepto-meningeal 139
B 144
NK/T H6
Erythroid lineage
Neutro-phil
Microglia 186
Main 173
PF 174
Main 175
PF 176
Neuron
Glial
GABA-ergic
Hybrid H10
Gluta-metergic
Astrocyte LHX2+ SLC7A10- 238
OPC 242
Oligo dendro-cyte 243
Ependymal 244
Schwann non-mye 249
GRIK1+ MEIS2+ NECAB1- 218
GIRK1- MEIS2+ NECAB1+ 219
GIRK1- MEIS2+ NECAB1- 220
PPP1R1B+ 221
NPTX1+ 223
b Brainstem
Endothelial
Stromal
Lymphoid
Myeloid
Neural
Vein 78
Capillary FABP5- RBP7- 83
Smooth musc vasc 109
Pericyte 130
Lepto-meningeal 139
B 144
NK/T H6
Erythroid lineage
Neutro-phil 175
Microglia 186
Main 173
PF 174
Neuron
Glial
GABA-ergic
Hybrid H10
Gluta-metergic
Astro-cyte
OPC 242
Oligo dendro-cyte 243
Ependymal
Choroid plexus 246
Schwann
OTX2+ 222
CBLN1+ 224
NEFH+ 225
ROM1+ 226
LHX2- SLC7A10+ 239
S100B+ 241
Main 244
NPY+ 245
Mye 248
Non-mye 249
c Eye retina
Myeloid
Neural
Erythroid lineage 173
Neutro-phil 175
Microglia 186
Neuron
Glial
Cone 227
Rod 228
Hor-zontal 229
Bipolar
Muller 247
Schwann non-mye 249
ON 230
Hybrid H11
OFF 231
d Hypothalamus/Pituitary
Endothelial
Stromal
Lymphoid
Myeloid
Neural
Capillary FABP5- RBP7- 83
Sinusoid MAFB+ 89
Smooth musc vasc 109
Fibroblast 111
Pericyte 130
Lepto-meningeal 139
B 144
Plasma 147
NK/T H6
HPC 169
Megaka-ryoid prog 170
Eryth-roid
Neutro-phil
Baso-phil 179
GMP 181
Mono/ Macro
Dendritic
Prog-enitor 172
Lineage main 173
Lineage PF 174
Main 175
PF 176
Mono main 182
Mono PF 183
Micro-glia 186
Conven-tional 211
Plasma-cytoid 212
Neuron GABAergic 221
Neuroendocrine
Glial
Cortico-troph 233
Gonado-troph 234
Lacto-troph 235
Somato-troph 236
Thyro-troph 237
Hybrid H12
Astrocyte LHX2+ SLC7A10+ 240
OPC 242
Oligo dendro-cyte 243
Ependymal 244
e Limb muscle
Endothelial
Stromal
Lymphoid
Myeloid
Artery 77
Vein 78
Capillary
Lymphatic
FABP5+ RBP7lo 85
FABP5+ RBP7+ 86
CXCL10+ 87
CCL21+ 92
ACKR4- CCDC80+ 93
Tendon 98
Skt musc stem
Skt musc fast 101
Skt musc slow 102
Smooth musc vasc 109
FAP 110
Pericyte 130
Adipo UCP1lo 133
Reticular 140
Main 99
PF 100*
B
NK/T
B main 144
B PF 145
Plasma main 147
Plasma PF 148
NK T 155
NK/T PF H7
T CD4+ CD8- CCL5- 157
T CD4+ CD8- CCL5- CLDN1+ 159
T CD4- CD8+ CCL5+ 160
T CD4- CD8+ CCL5- 161
Erythroid lineage 173
Neutro-phil 175
Baso-phil 179
Macro-phage 184
f Limb muscle
85 - Capillary (FABP5+ RBP7lo)
87 - Capillary (CXCL10+)
78 - Vein
77 - Artery
93 - Lymphatic (ACKR4- CCDC80+)
86 - Capillary (FABP5+ RBP7+)
92 - Lymphatic (CCL21+)
UMAP2
UMAP1
L2
Endothelium
99 - Skeletal muscle satellite stem
101 - Fast muscle
110 - Fibroadipogenic progenitor
102 - Slow muscle
98 - Tendon
133 - Adipocyte (UCP1lo)
130 - Pericyte
109 - Vascular SMC
140 - Reticular
Stroma
144 - B
145 - B (PF)
148 - Plasma (PF)
157 - T (CD4+ CD8- CCL5-)
147 - Plasma
155 - NKT
160 - T (CD4- CD8+ CCL5+)
161 - T (CD4- CD8+ CCL5-)
H7 - NK/T (PF)
159 - T (CD4+ CD8- CCL5- CLDN1+)
Lymphoid
175 - Neutrophil
184 - Macrophage
179 - Basophil
173 - Erythroid lineage
Myeloid

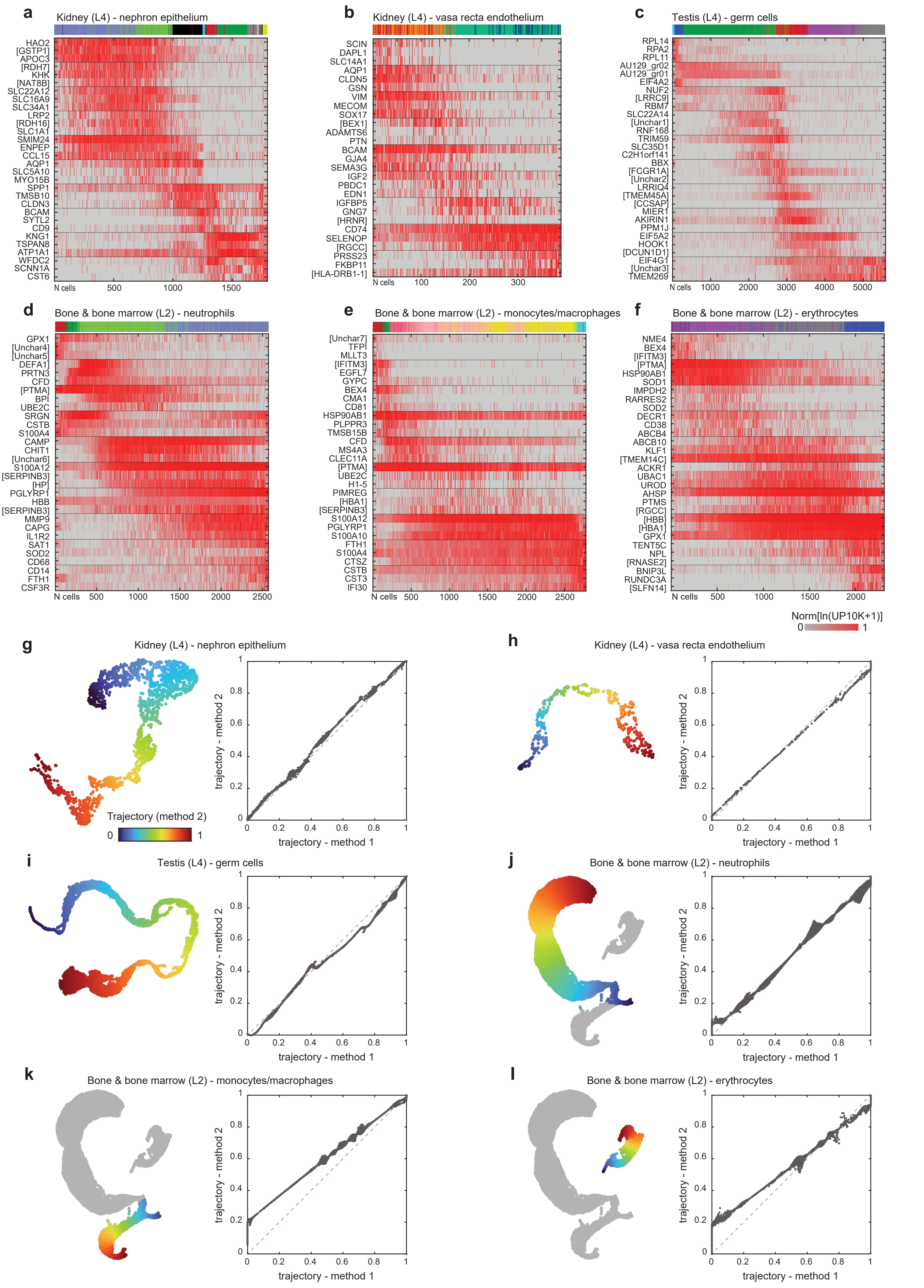
a
Kidney (L4) - nephron epithelium
b
Kidney (L4) - vasa recta endothelium
c
Testis (L4) - germ cells
d
Bone & bone marrow (L2) - neutrophils
e
Bone & bone marrow (L2) - monocytes/macrophages
f
Bone & bone marrow (L2) - erythrocytes
N cells
Norm[ln(UP10K+1)]
g
Kidney (L4) - nephron epithelium
Trajectory (method 2)
h
Kidney (L4) - vasa recta endothelium
i
Testis (L4) - germ cells
j
Bone & bone marrow (L2) - neutrophils
k
Bone & bone marrow (L2) - monocytes/macrophages
l
Bone & bone marrow (L2) - erythrocytes
trajectory - method 1
trajectory - method 2

**Fig. S3 - continued**

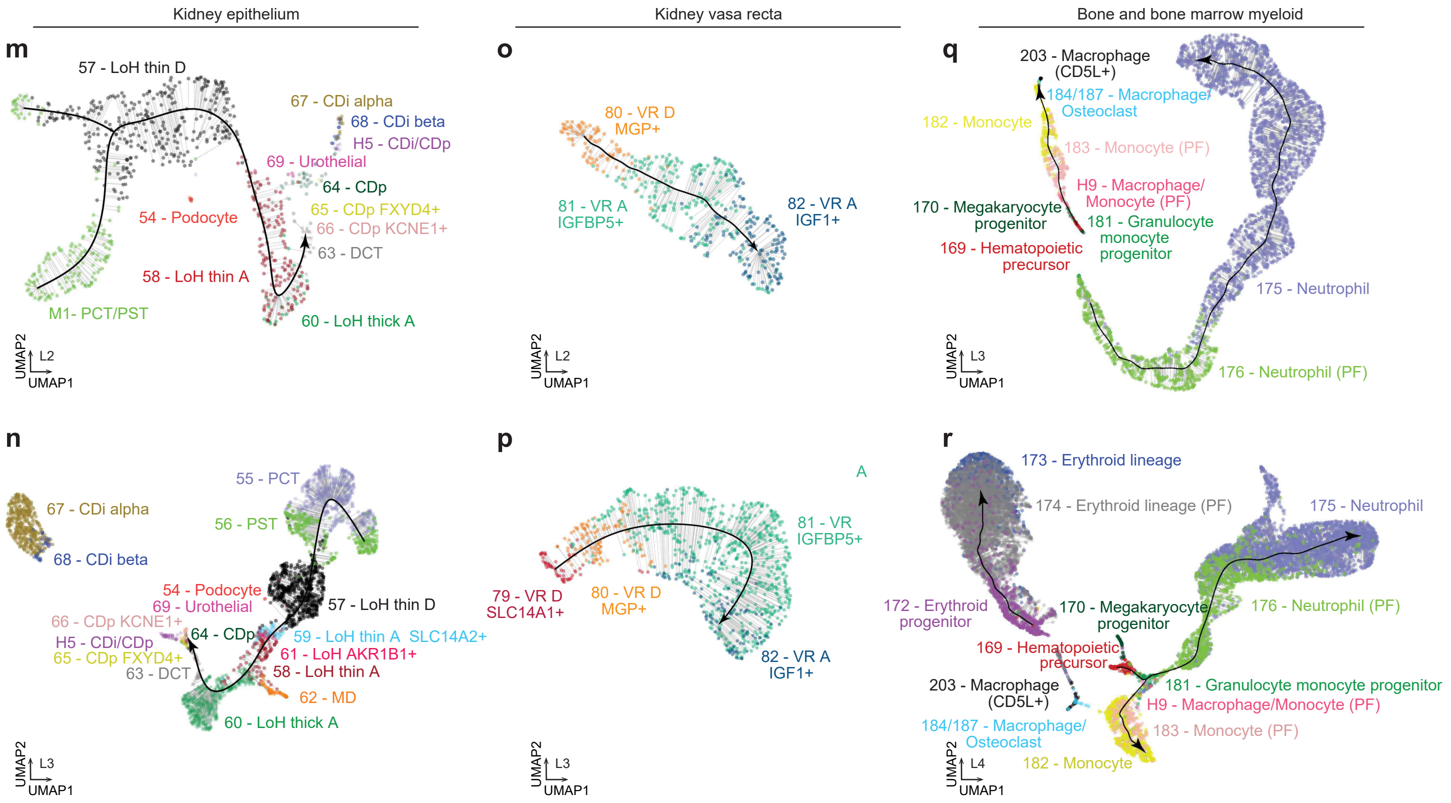

### Fig. S4

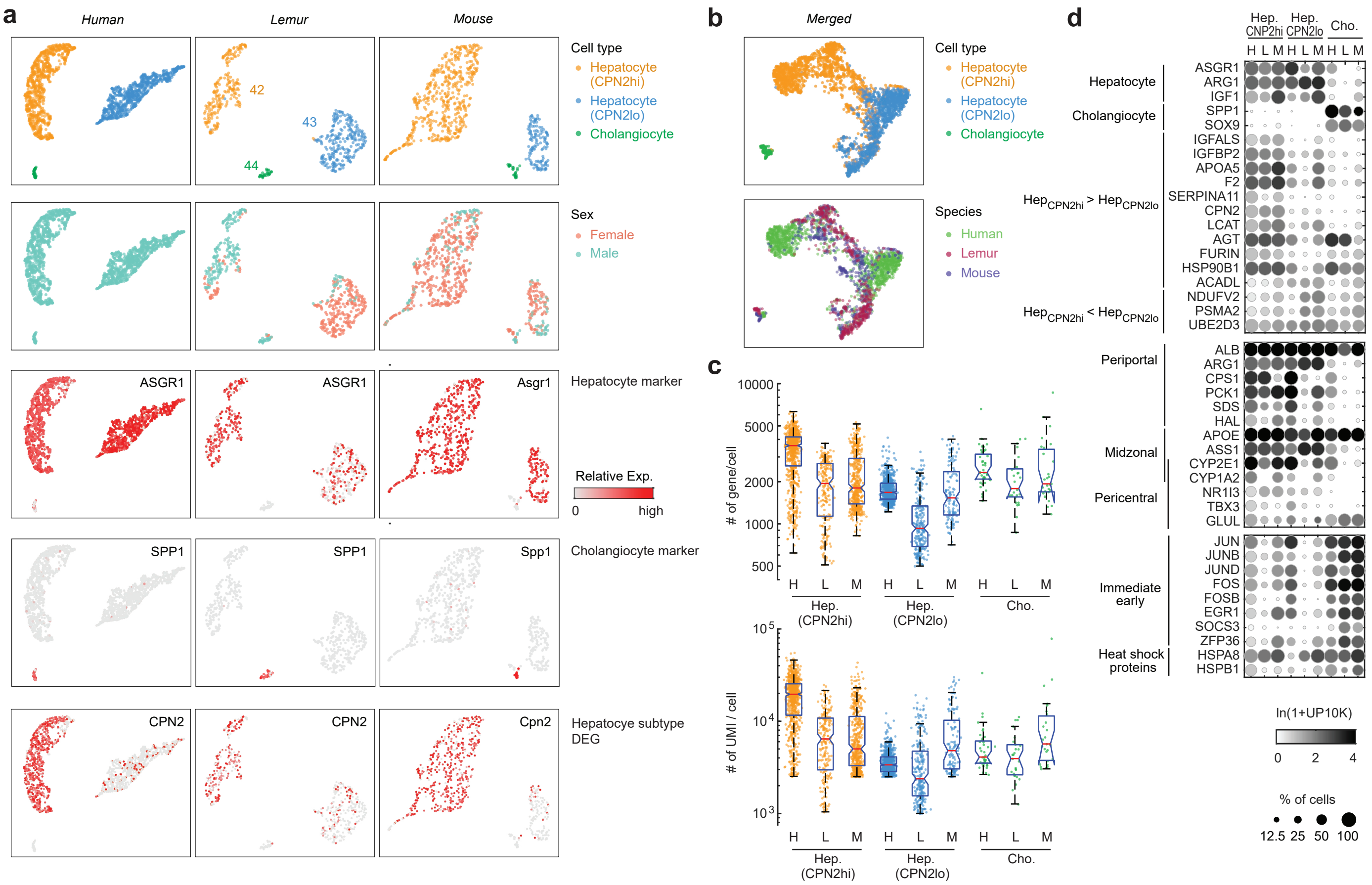

a
Human
Lemur
Mouse
42
43
44
Cell type
Hepatocyte (CPN2hi)
Hepatocyte (CPN2lo)
Cholangiocyte
Sex
Female
Male
ASGR1
Asgr1
Hepatocyte marker
Relative Exp.
0
high
SPP1
Spp1
Cholangiocyte marker
CPN2
Cpn2
Hepatocyte subtype DEG
b
Merged
Species
Human
Lemur
Mouse
c
### of gene/cell
10000
5000
2000
1000
500
### of UMI / cell
$10^5$
$10^4$
$10^3$
H L M H L M H L M
Hep. (CPN2hi)
Hep. (CPN2lo)
Cho.
d
Hep. CNP2hi
Hep. CPN2lo
Cho.
Hepatocyte
Cholangiocyte
$Hep_{CPN2hi}$ > $Hep_{CPN2lo}$
$Hep_{CPN2hi}$ < $Hep_{CPN2lo}$
ASGR1
ARG1
IGF1
SPP1
SOX9
IGFALS
IGFBP2
APOA5
F2
SERPINA11
CPN2
LCAT
AGT
FURIN
HSP90B1
ACADL
NDUFV2
PSMA2
UBE2D3
Periportal
Midzonal
Pericentral
ALB
ARG1
CPS1
PCK1
SDS
HAL
APOE
ASS1
CYP2E1
CYP1A2
NR1I3
TBX3
GLUL
Immediate early
Heat shock proteins
JUN
JUNB
JUND
FOS
FOSB
EGR1
SOCS3
ZFP36
HSPA8
HSPB1
ln(1+UP10K)
0
2
4
% of cells
12.5
25
50
100

Fig. S5

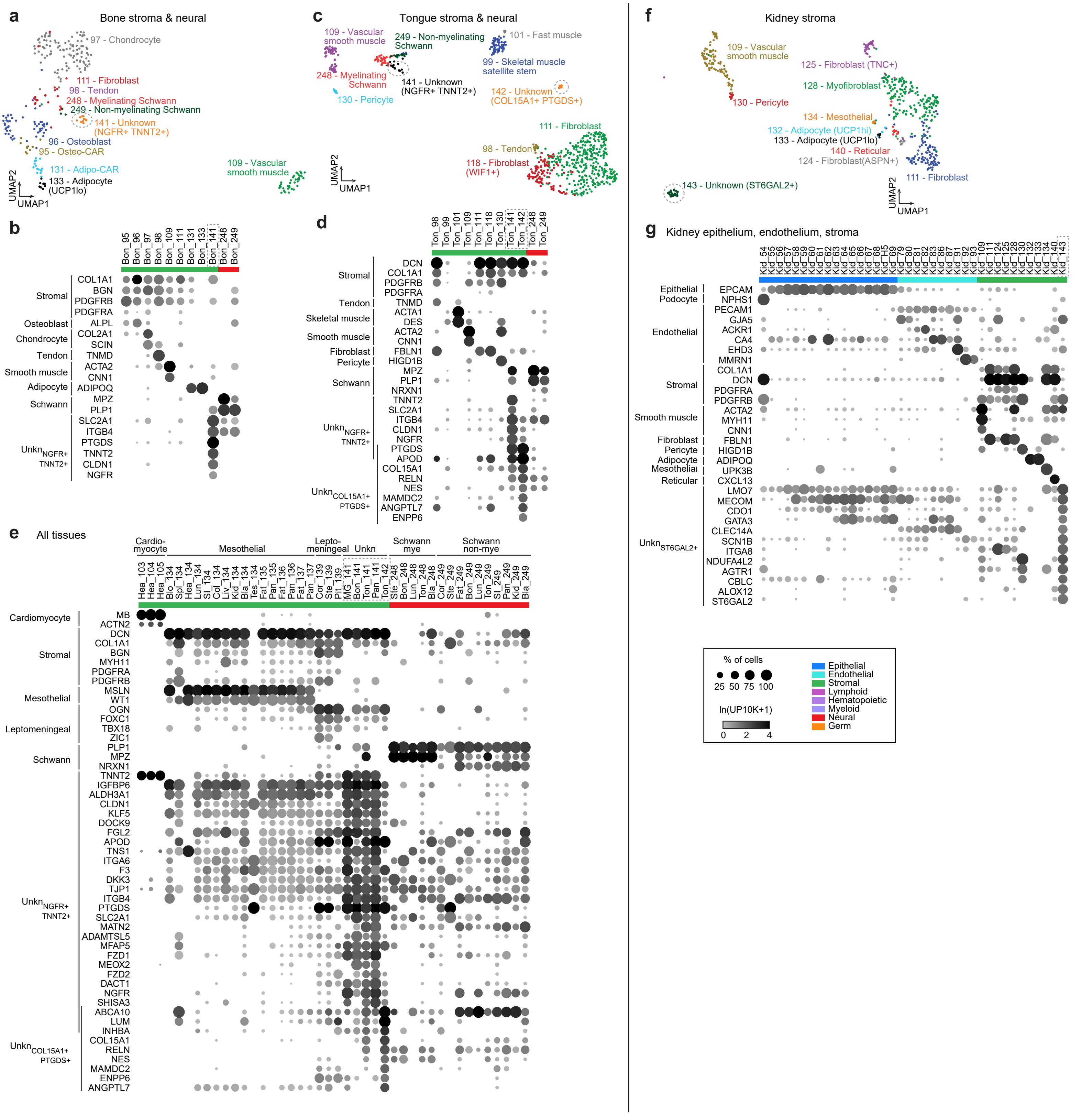

a
Bone stroma & neural
97 - Chondrocyte
111 - Fibroblast
98 - Tendon
248 - Myelinating Schwann
249 - Non-myelinating Schwann
141 - Unknown (NGFR+ TNNT2+)
96 - Osteoblast
95 - Osteo-CAR
109 - Vascular smooth muscle
131 - Adipo-CAR
133 - Adipocyte (UCP1lo)
UMAP2
UMAP1
b
Bon_95 Bon_96 Bon_97 Bon_98 Bon_109 Bon_111 Bon_131 Bon_133 Bon_141 Bon_248 Bon_249
Stromal COL1A1 BGN PDGFRB PDGFRA
Osteoblast ALPL
Chondrocyte COL2A1 SCIN
Tendon TNMD
Smooth muscle ACTA2 CNN1
Adipocyte ADIPOQ
Schwann MPZ PLP1
Unkn NGFR+ TNNT2+ SLC2A1 ITGB4 PTGDS TNNT2 CLDN1 NGFR
c
Tongue stroma & neural
109 - Vascular smooth muscle
249 - Non-myelinating Schwann
101 - Fast muscle
99 - Skeletal muscle satellite stem
248 - Myelinating Schwann
141 - Unknown (NGFR+ TNNT2+)
130 - Pericyte
142 - Unknown (COL15A1+ PTGDS+)
111 - Fibroblast
98 - Tendon
118 - Fibroblast (WIF1+)
UMAP2
UMAP1
d
Ton_98 Ton_99 Ton_101 Ton_109 Ton_111 Ton_118 Ton_130 Ton_141 Ton_142 Ton_248 Ton_249
Stromal DCN COL1A1 PDGFRB PDGFRA
Tendon TNMD
Skeletal muscle ACTA1 DES
Smooth muscle ACTA2 CNN1
Fibroblast FBLN1
Pericyte HIGD1B
Schwann MPZ PLP1 NRXN1
Unkn NGFR+ TNNT2+ TNNT2 SLC2A1 ITGB4 CLDN1 NGFR
Unkn COL15A1+ PTGDS+ PTGDS APOD COL15A1 RELN NES MAMDC2 ANGPTL7 ENPP6
e
All tissues
Cardio-myocyte
Mesothelial
Lepto-meningeal
Unkn
Schwann mye
Schwann non-mye
Hea_103 Hea_104 Hea_105 Blo_134 Spl_134 Hea_134 Lun_134 SI_134 Col_134 Liv_134 Kid_134 Bla_134 Tes_134 Fat_135 Pan_135 Fat_136 Pan_136 Fat_137 Pan_137 Cor_139 Ste_139 Pit_139 MG_141 Bon_141 Ton_141 Pan_141 Ton_142 Ste_248 Bon_248 Lun_248 Ton_248 Bla_248 Cor_249 Ste_249 Fat_249 Bon_249 Lun_249 Ton_249 SI_249 Pan_249 Kid_249 Bla_249
Cardiomyocyte MB ACTN2
Stromal DCN COL1A1 BGN MYH11 PDGFRA PDGFRB
Mesothelial MSLN WT1
Leptomeningeal OGN FOXC1 TBX18 ZIC1
Schwann PLP1 MPZ NRXN1
Unkn NGFR+ TNNT2+ TNNT2 IGFBP6 ALDH3A1 CLDN1 KLF5 DOCK9 FGL2 APOD TNS1 ITGA6 F3 DKK3 TJP1 ITGB4 PTGDS SLC2A1 MATN2 ADAMTSL5 MFAP5 FZD1 MEOX2 FZD2 DACT1 NGFR SHISA3
Unkn COL15A1+ PTGDS+ ABCA10 LUM INHBA COL15A1 RELN NES MAMDC2 ENPP6 ANGPTL7
f
Kidney stroma
109 - Vascular smooth muscle
125 - Fibroblast (TNC+)
128 - Myofibroblast
130 - Pericyte
134 - Mesothelial
132 - Adipocyte (UCP1hi)
133 - Adipocyte (UCP1lo)
140 - Reticular
124 - Fibroblast(ASPN+)
111 - Fibroblast
143 - Unknown (ST6GAL2+)
UMAP2
UMAP1
g
Kidney epithelium, endothelium, stroma
Kid_54 Kid_55 Kid_56 Kid_57 Kid_58 Kid_59 Kid_60 Kid_61 Kid_62 Kid_63 Kid_64 Kid_65 Kid_66 Kid_67 Kid_68 Kid_H5 Kid_69 Kid_79 Kid_80 Kid_81 Kid_82 Kid_83 Kid_86 Kid_87 Kid_91 Kid_92 Kid_93 Kid_109 Kid_111 Kid_124 Kid_125 Kid_128 Kid_130 Kid_132 Kid_133 Kid_134 Kid_140 Kid_143
Epithelial EPCAM
Podocyte NPHS1
Endothelial PECAM1 GJA5 ACKR1 CA4 EHD3 MMRN1
Stromal COL1A1 DCN PDGFRA PDGFRB
Smooth muscle ACTA2 MYH11 CNN1
Fibroblast FBLN1
Pericyte HIGD1B
Adipocyte ADIPOQ
Mesothelial UPK3B
Reticular CXCL13
Unkn ST6GAL2+ LMO7 MECOM CDO1 GATA3 CLEC14A SCN1B ITGA8 NDUFA4L2 AGTR1 CBLC ALOX12 ST6GAL2
% of cells
25 50 75 100
ln(UP10K+1)
0 2 4
Epithelial
Endothelial
Stromal
Lymphoid
Hematopoietic
Myeloid
Neural
Germ

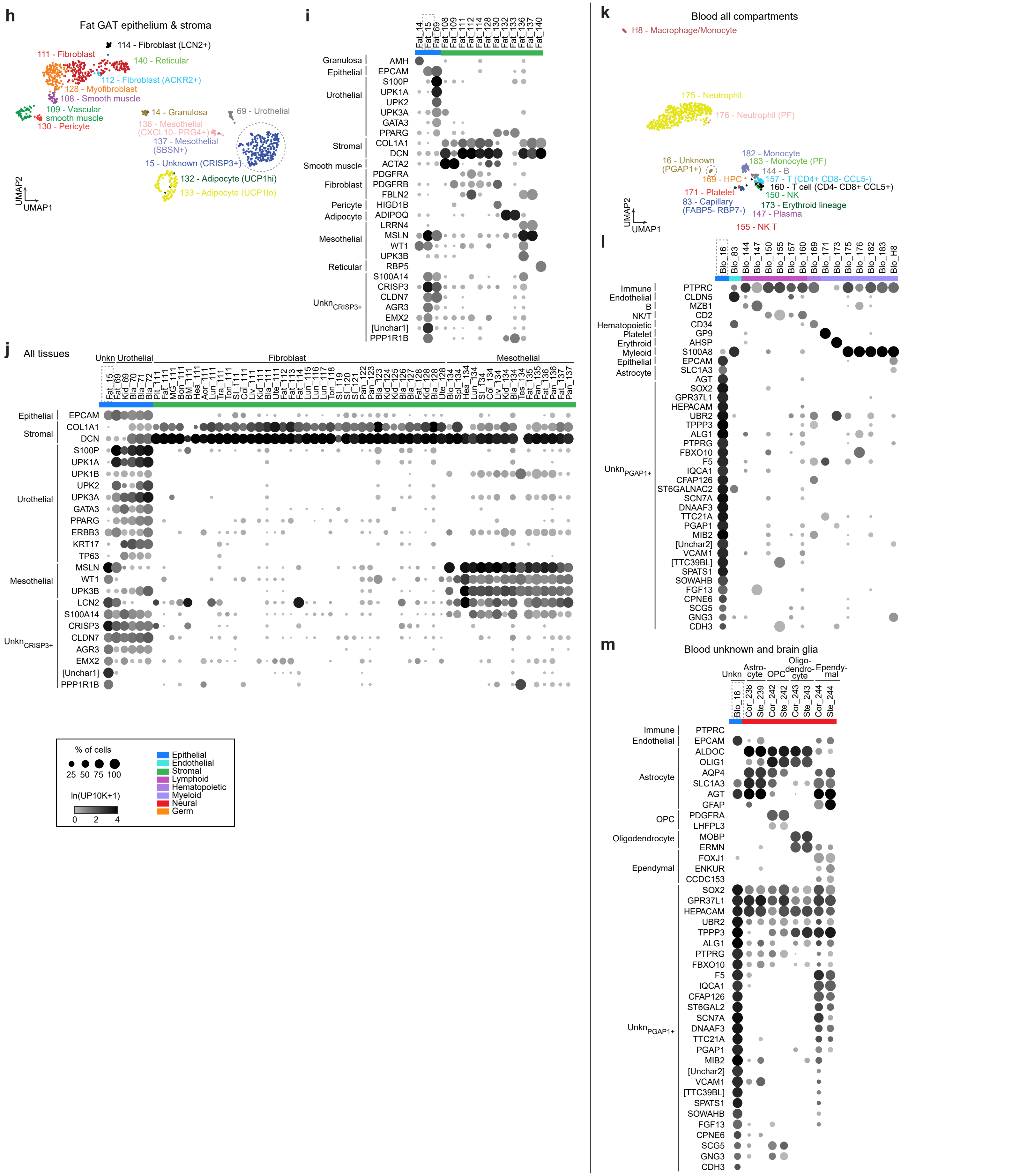
h
Fat GAT epithelium & stroma
114 - Fibroblast (LCN2+)
111 - Fibroblast
140 - Reticular
112 - Fibroblast (ACKR2+)
128 - Myofibroblast
108 - Smooth muscle
109 - Vascular smooth muscle
130 - Pericyte
14 - Granulosa
69 - Urothelial
136 - Mesothelial (CXCL10- PRG4+)
137 - Mesothelial (SBSN+)
15 - Unknown (CRISP3+)
132 - Adipocyte (UCP1hi)
133 - Adipocyte (UCP1lo)
UMAP2
UMAP1
i
j
All tissues
Unkn Urothelial
Fibroblast
Mesothelial
k
Blood all compartments
H8 - Macrophage/Monocyte
175 - Neutrophil
176 - Neutrophil (PF)
16 - Unknown (PGAP1+)
182 - Monocyte
183 - Monocyte (PF)
144 - B
169 - HPC
157 - T (CD4+ CD8- CCL5-)
160 - T cell (CD4- CD8+ CCL5+)
150 - NK
171 - Platelet
83 - Capillary (FABP5- RBP7-)
173 - Erythroid lineage
147 - Plasma
155 - NK T
l
m
Blood unknown and brain glia
Unkn
Astro-cyte
OPC
Oligo-dendro-cyte
Ependy-mal
% of cells
25 50 75 100
ln(UP10K+1)
0 2 4
Epithelial
Endothelial
Stromal
Lymphoid
Hematopoietic
Myeloid
Neural
Germ

**Fig. S6**

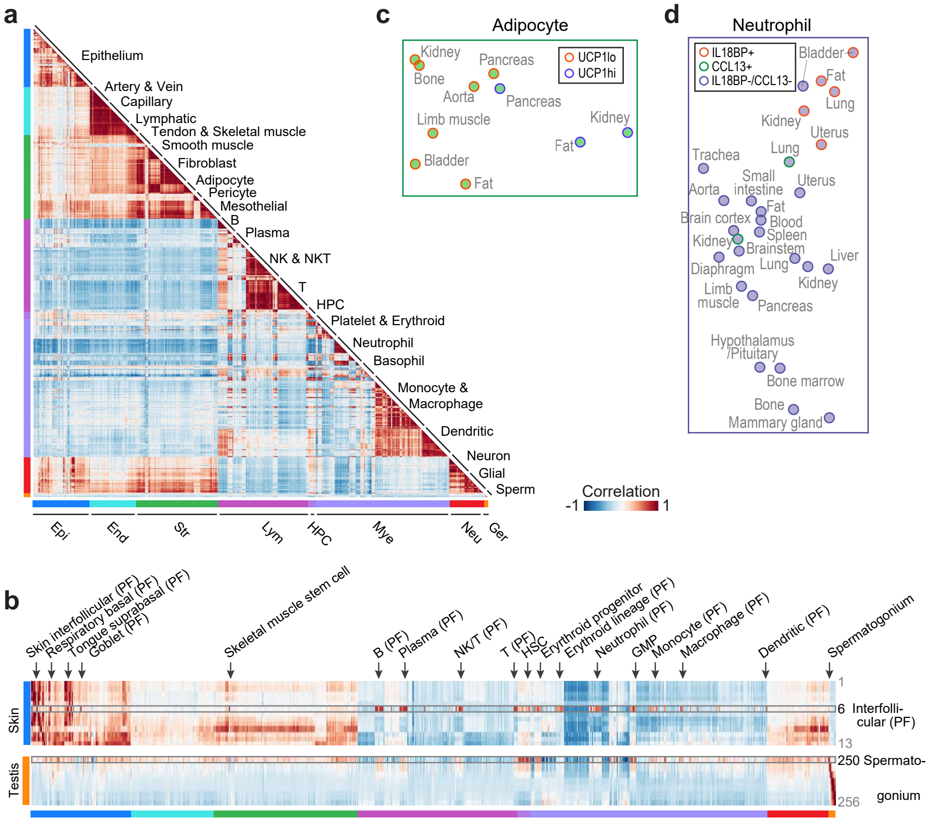
a
Epithelium
Artery & Vein
Capillary
Lymphatic
Tendon & Skeletal muscle
Smooth muscle
Fibroblast
Adipocyte
Pericyte
Mesothelial
B
Plasma
NK & NKT
T
HPC
Platelet & Erythroid
Neutrophil
Basophil
Monocyte &
Macrophage
Dendritic
Neuron
Glial
Sperm
Correlation
-1
1
Epi
End
Str
Lym
HPC
Mye
Neu
Ger
c
Adipocyte
UCP1lo
UCP1hi
Kidney
Bone
Pancreas
Aorta
Pancreas
Limb muscle
Kidney
Fat
Bladder
Fat
d
Neutrophil
IL18BP+
CCL13+
IL18BP-/CCL13-
Bladder
Fat
Lung
Kidney
Uterus
Trachea
Lung
Aorta
Small
intestine
Uterus
Fat
Brain cortex
Blood
Spleen
Kidney
Brainstem
Diaphragm
Lung
Liver
Kidney
Limb
muscle
Pancreas
Hypothalamus
/Pituitary
Bone marrow
Bone
Mammary gland
b
Skin interfollicular (PF)
Respiratory basal (PF)
Tongue suprabasal (PF)
Goblet (PF)
Skeletal muscle stem cell
B (PF)
Plasma (PF)
NK/T (PF)
T (PF)
HSC
Erythroid progenitor
Erythroid lineage (PF)
Neutrophil (PF)
GMP
Monocyte (PF)
Macrophage (PF)
Dendritic (PF)
Spermatogonium
Skin
1
6 Interfolli-
cular (PF)
13
Testis
250 Spermato-
gonium
256

a. Correlation heatmap A - ordered by compartment, cell type, tissue

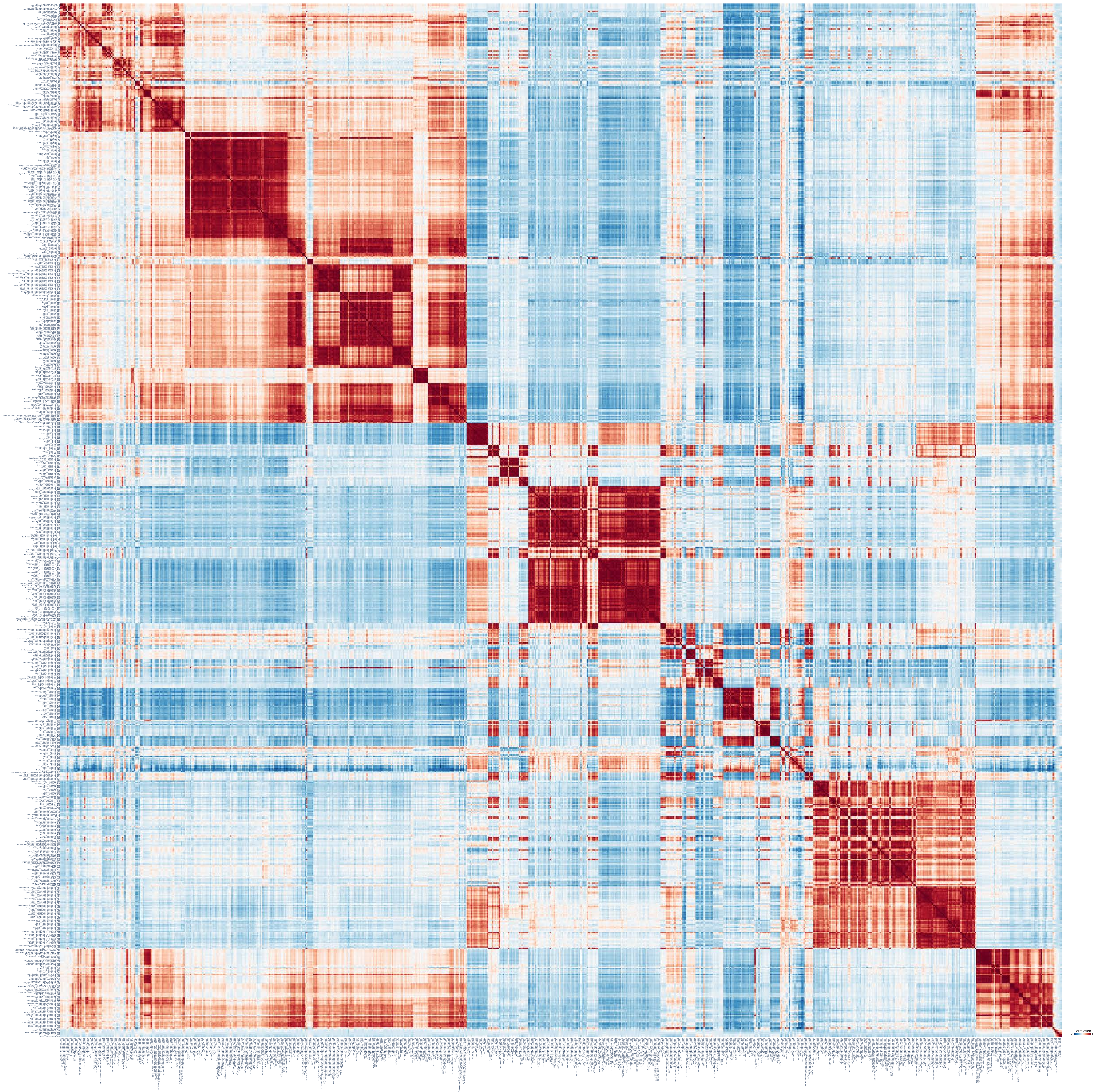

b. Correlation heatmap B - ordered by tissue, compartment, cell type

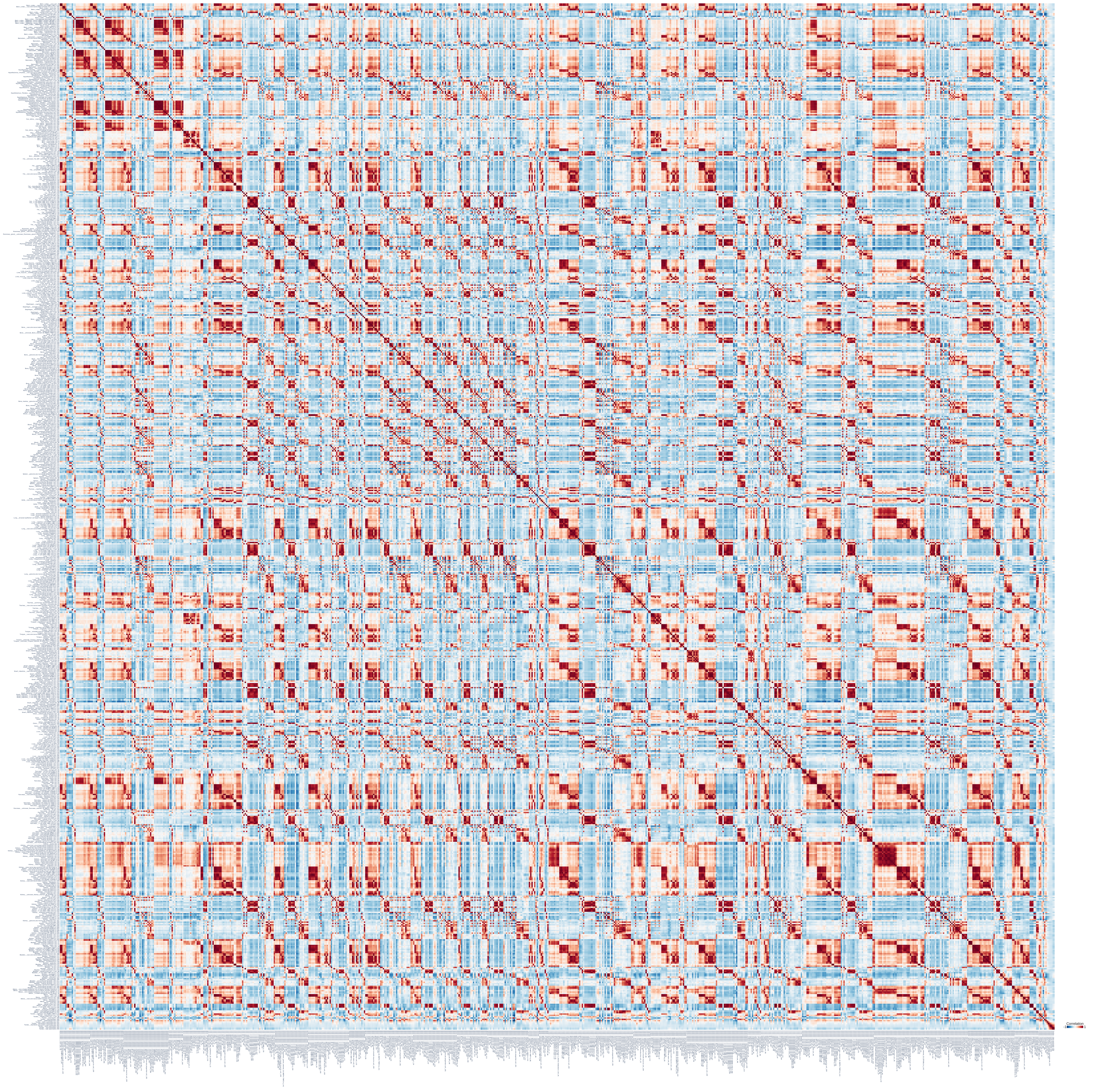

c. Correlation heatmap C - ordered by hierarchical clustering

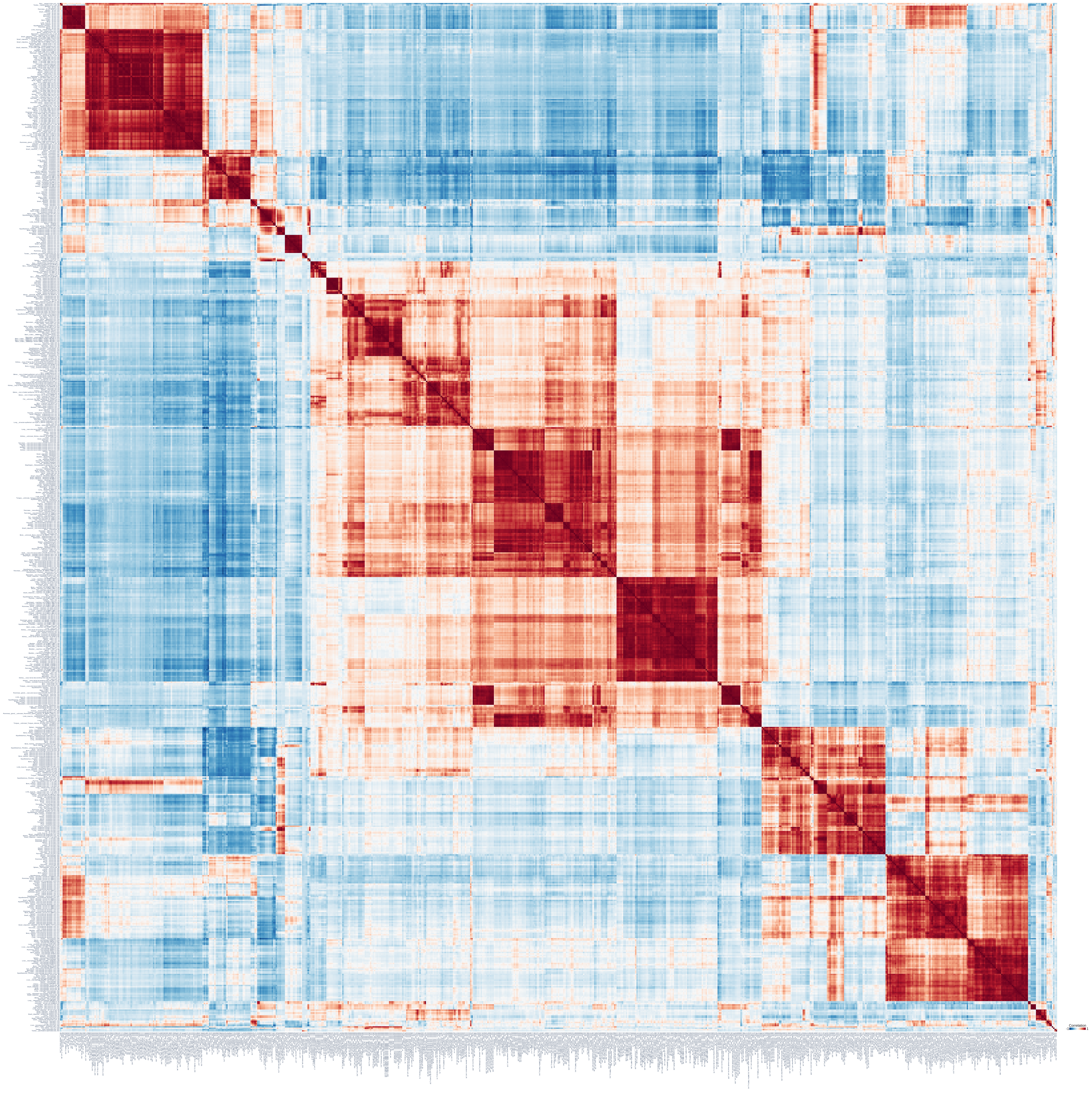

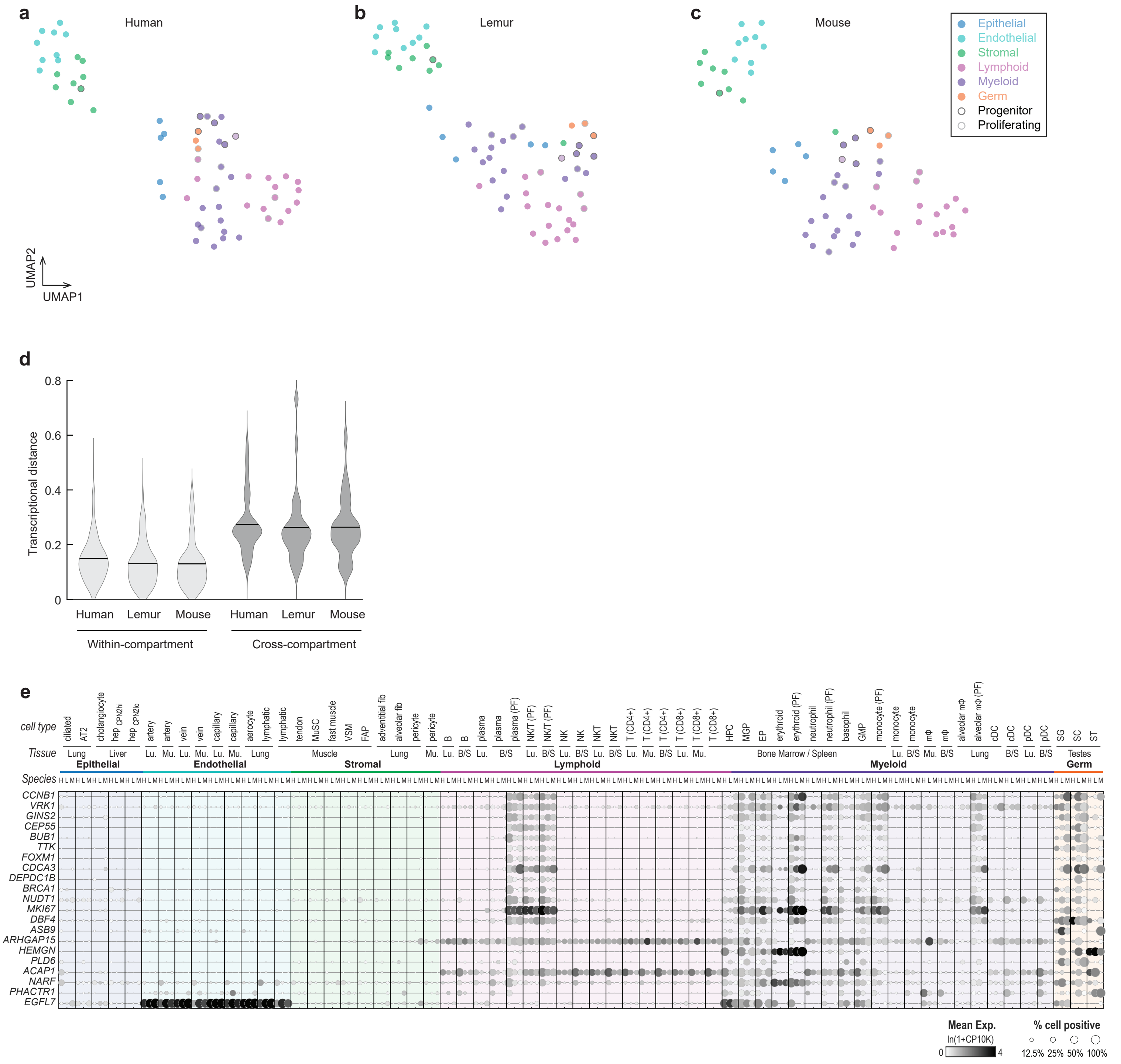
a
Human
b
Lemur
c
Mouse
Epithelial
Endothelial
Stromal
Lymphoid
Myeloid
Germ
Progenitor
Proliferating
UMAP2
UMAP1
d
Transcriptional distance
0.8
0.6
0.4
0.2
0
Human
Lemur
Mouse
Human
Lemur
Mouse
Within-compartment
Cross-compartment
e
cell type
Tissue
Species
Epithelial
Endothelial
Stromal
Lymphoid
Myeloid
Germ
CCNB1
VRK1
GINS2
CEP55
BUB1
TTK
FOXM1
CDCA3
DEPDC1B
BRCA1
NUDT1
MKI67
DBF4
ASB9
ARHGAP15
HEMGN
PLD6
ACAP1
NARF
PHACTR1
EGFL7
Mean Exp.
ln(1+CP10K)
0
4
% cell positive
12.5%
25%
50%
100%

**Fig. S9**

**a** Lung across species (integarated by Portal)

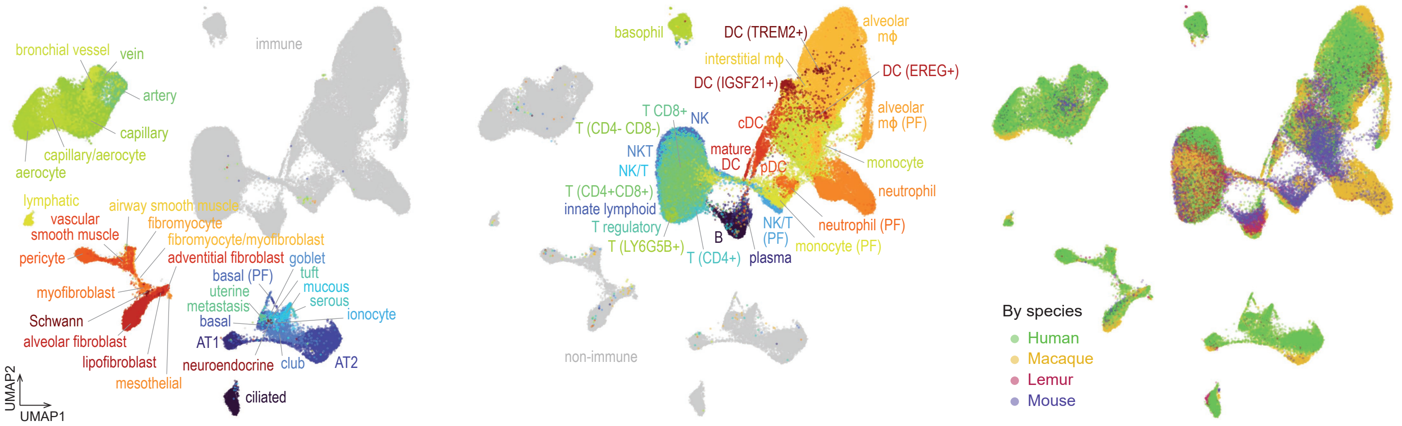


**b** Lung across species (aligned by SAMap)

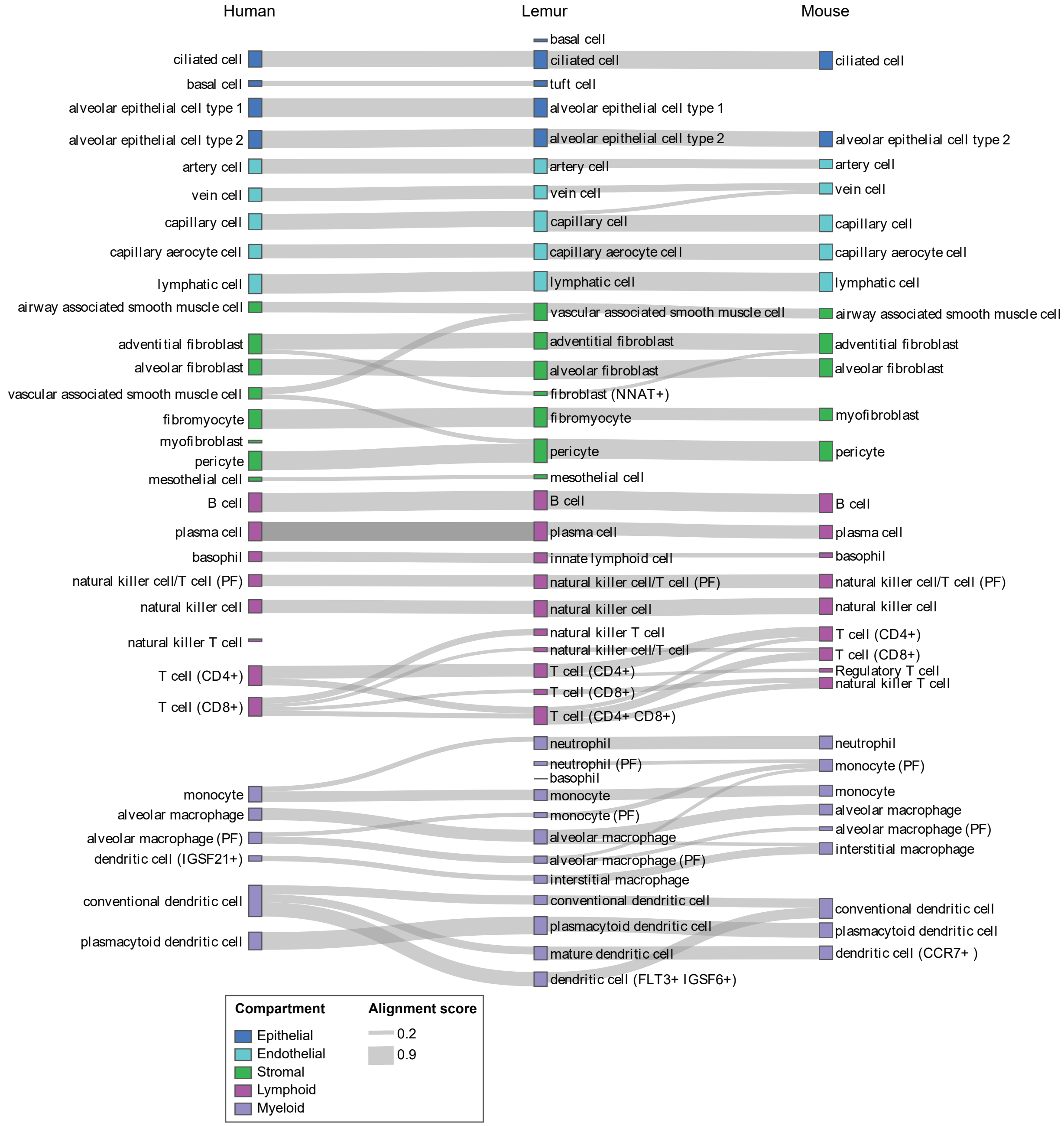

**c** Skeletal muscle across species (integarated by Portal)

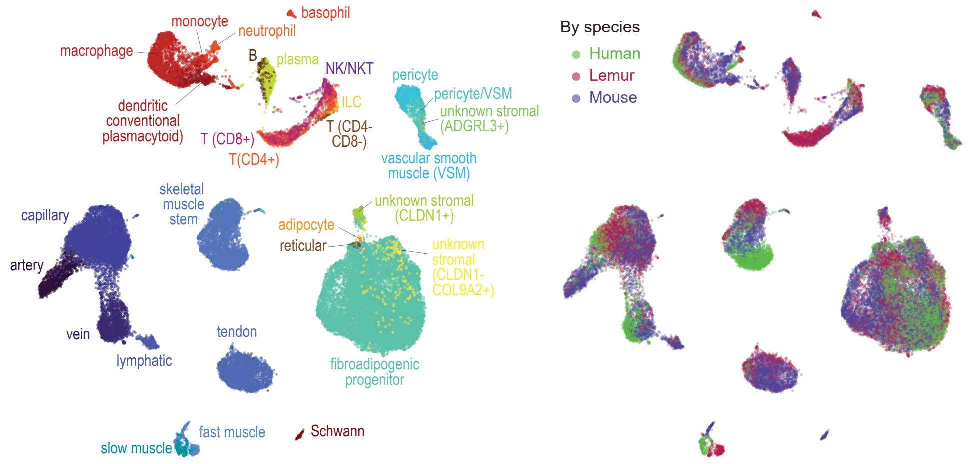


**d** Skeletal muscle across species (aligned by SAMap)

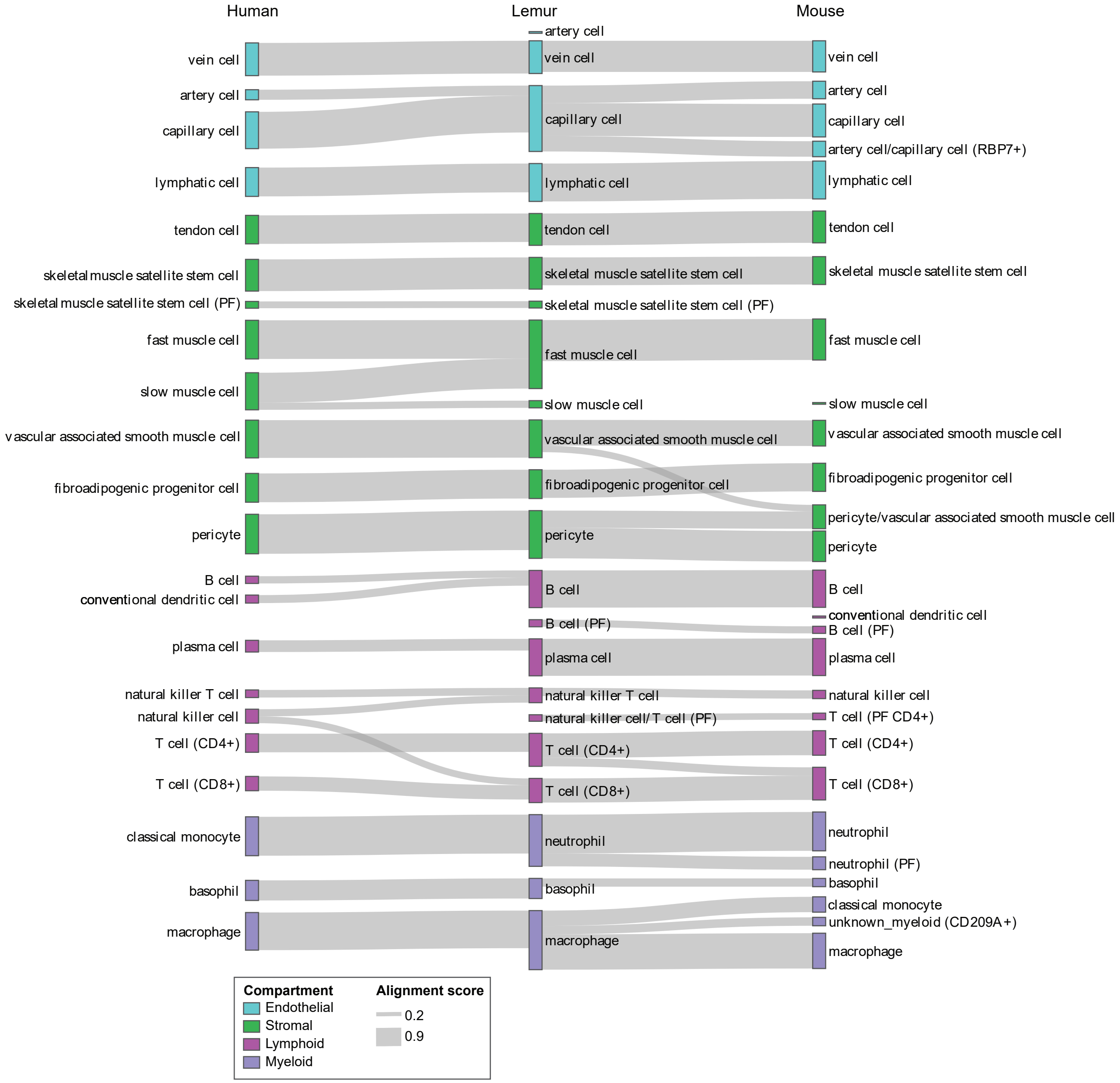

### Fig. S10

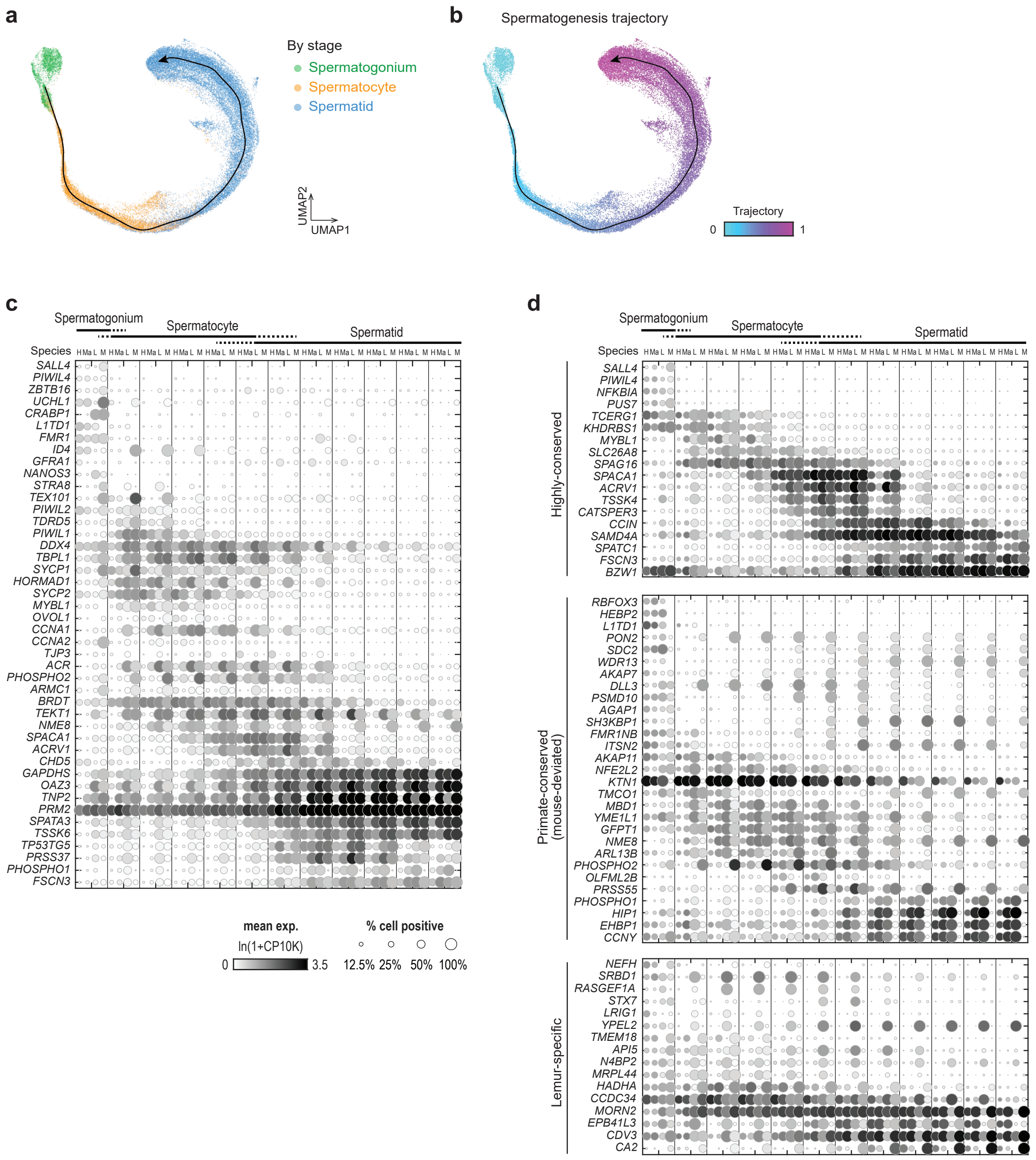

a
By stage
Spermatogonium
Spermatocyte
Spermatid
UMAP2
UMAP1
b
Spermatogenesis trajectory
Trajectory
0
1
c
Spermatogonium
Spermatocyte
Spermatid
Species
SALL4
PIWIL4
ZBTB16
UCHL1
CRABP1
L1TD1
FMR1
ID4
GFRA1
NANOS3
STRA8
TEX101
PIWIL2
TDRD5
PIWIL1
DDX4
TBPL1
SYCP1
HORMAD1
SYCP2
MYBL1
OVOL1
CCNA1
CCNA2
TJP3
ACR
PHOSPHO2
ARMC1
BRDT
TEKT1
NME8
SPACA1
ACRV1
CHD5
GAPDHS
OAZ3
TNP2
PRM2
SPATA3
TSSK6
TP53TG5
PRSS37
PHOSPHO1
FSCN3
mean exp.
ln(1+CP10K)
0
3.5
% cell positive
12.5% 25% 50% 100%
d
Spermatogonium
Spermatocyte
Spermatid
Species
Highly-conserved
SALL4
PIWIL4
NFKBIA
PUS7
TCERG1
KHDRBS1
MYBL1
SLC26A8
SPAG16
SPACA1
ACRV1
TSSK4
CATSPER3
CCIN
SAMD4A
SPATC1
FSCN3
BZW1
Primate-conserved (mouse-deviated)
RBFOX3
HEBP2
L1TD1
PON2
SDC2
WDR13
AKAP7
DLL3
PSMD10
AGAP1
SH3KBP1
FMR1NB
ITSN2
AKAP11
NFE2L2
KTN1
TMCO1
MBD1
YME1L1
GFPT1
NME8
ARL13B
PHOSPHO2
OLFML2B
PRSS55
PHOSPHO1
HIP1
EHBP1
CCNY
Lemur-specific
NEFH
SRBD1
RASGEF1A
STX7
LRIG1
YPEL2
TMEM18
API5
N4BP2
MRPL44
HADHA
CCDC34
MORN2
EPB41L3
CDV3
CA2

**Fig. S10 - continued**

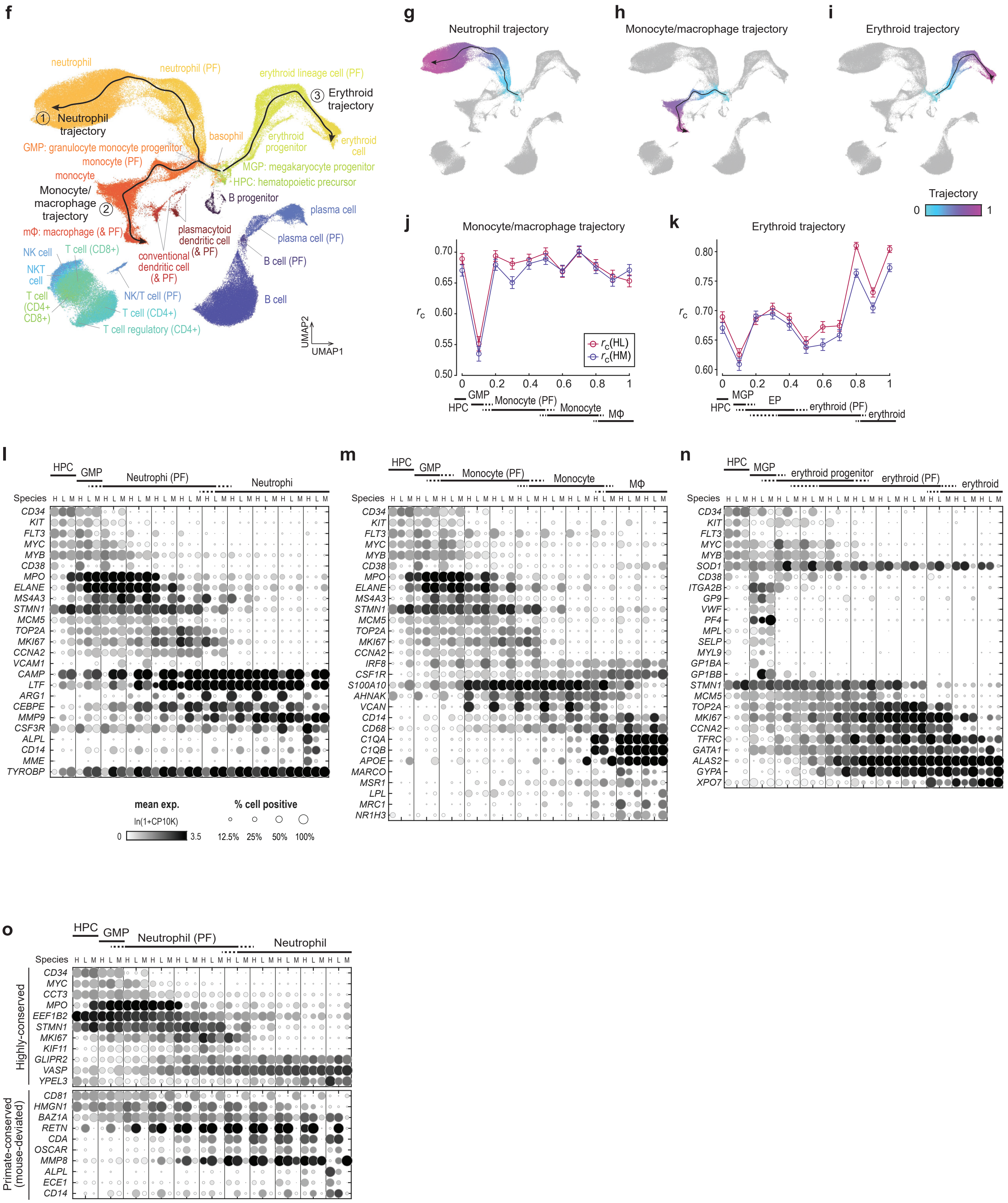

### Fig. S11

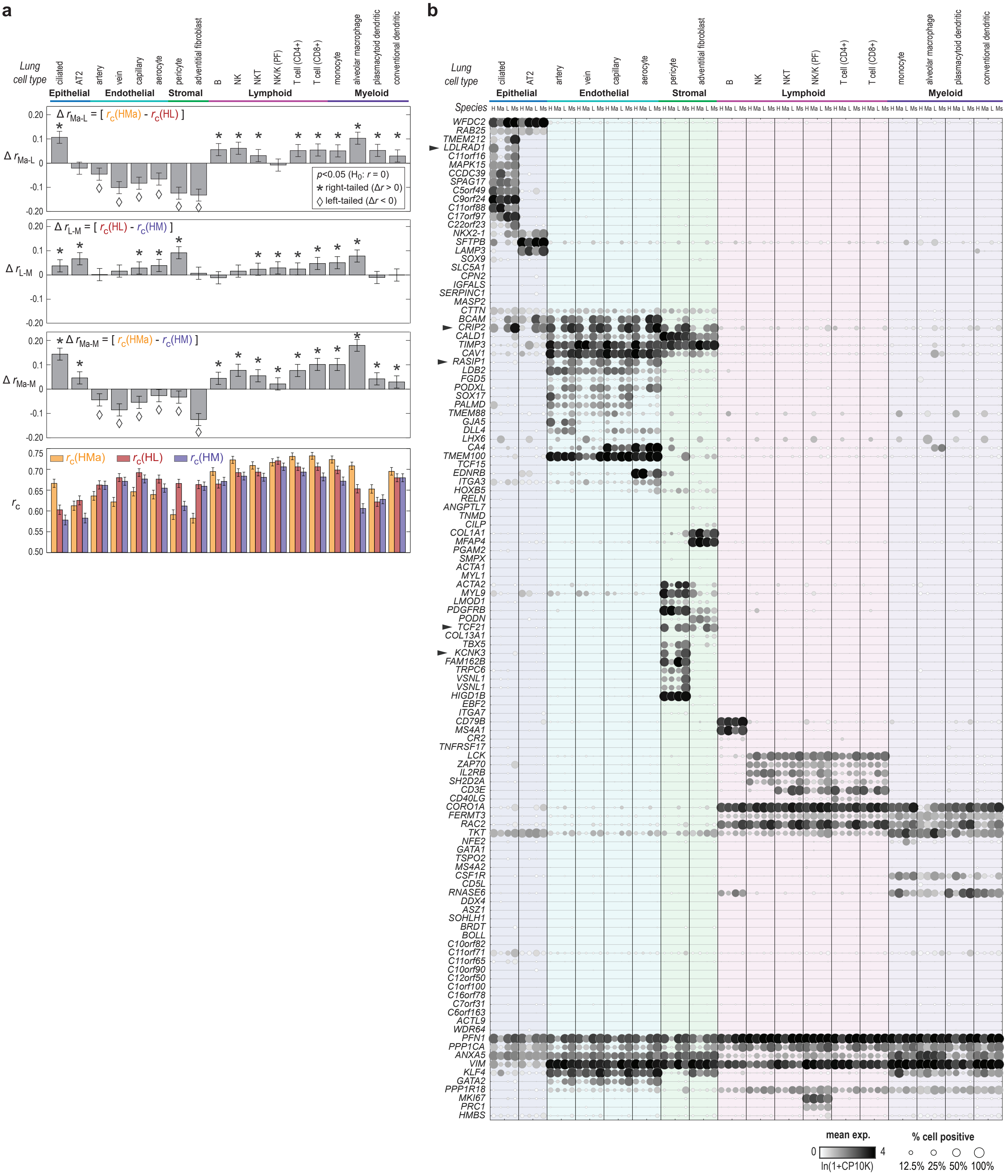

a
b
Lung cell type
ciliated
AT2
artery
vein
capillary
aerocyte
pericyte
adventitial fibroblast
B
NK
NKT
NK/K (PF)
T cell (CD4+)
T cell (CD8+)
monocyte
alveolar macrophage
plasmacytoid dendritic
conventional dendritic
Epithelial
Endothelial
Stromal
Lymphoid
Myeloid
Δ r_Ma-L = [ r_c(HMa) - r_c(HL) ]
Δ r_L-M = [ r_c(HL) - r_c(HM) ]
Δ r_Ma-M = [ r_c(HMa) - r_c(HM) ]
p<0.05 (H0: r = 0)
right-tailed (Δr > 0)
left-tailed (Δr < 0)
r_c(HMa)
r_c(HL)
r_c(HM)
r_c
Species
H Ma L Ms
WFDC2
RAB25
TMEM212
LDLRAD1
C11orf16
MAPK15
CCDC39
SPAG17
C5orf49
C9orf24
C11orf88
C17orf97
C22orf23
NKX2-1
SFTPB
LAMP3
SOX9
SLC5A1
CPN2
IGFALS
SERPINC1
MASP2
CTTN
BCAM
CRIP2
CALD1
TIMP3
CAV1
RASIP1
LDB2
FGD5
PODXL
SOX17
PALMD
TMEM88
GJA5
DLL4
LHX6
CA4
TMEM100
TCF15
EDNRB
ITGA3
HOXB5
RELN
ANGPTL7
TNMD
CILP
COL1A1
MFAP4
PGAM2
SMPX
ACTA1
MYL1
ACTA2
MYL9
LMOD1
PDGFRB
PODN
TCF21
COL13A1
TBX5
KCNK3
FAM162B
TRPC6
VSNL1
VSNL1
HIGD1B
EBF2
ITGA7
CD79B
MS4A1
CR2
TNFRSF17
LCK
ZAP70
IL2RB
SH2D2A
CD3E
CD40LG
CORO1A
FERMT3
RAC2
TKT
NFE2
GATA1
TSPO2
MS4A2
CSF1R
CD5L
RNASE6
DDX4
ASZ1
SOHLH1
BRDT
BOLL
C10orf82
C11orf71
C11orf65
C10orf90
C12orf50
C1orf100
C16orf78
C7orf31
C6orf163
ACTL9
WDR64
PFN1
PPP1CA
ANXA5
VIM
KLF4
GATA2
PPP1R18
MKI67
PRC1
HMBS
mean exp.
0
4
ln(1+CP10K)
% cell positive
12.5% 25% 50% 100%

### Fig. S12

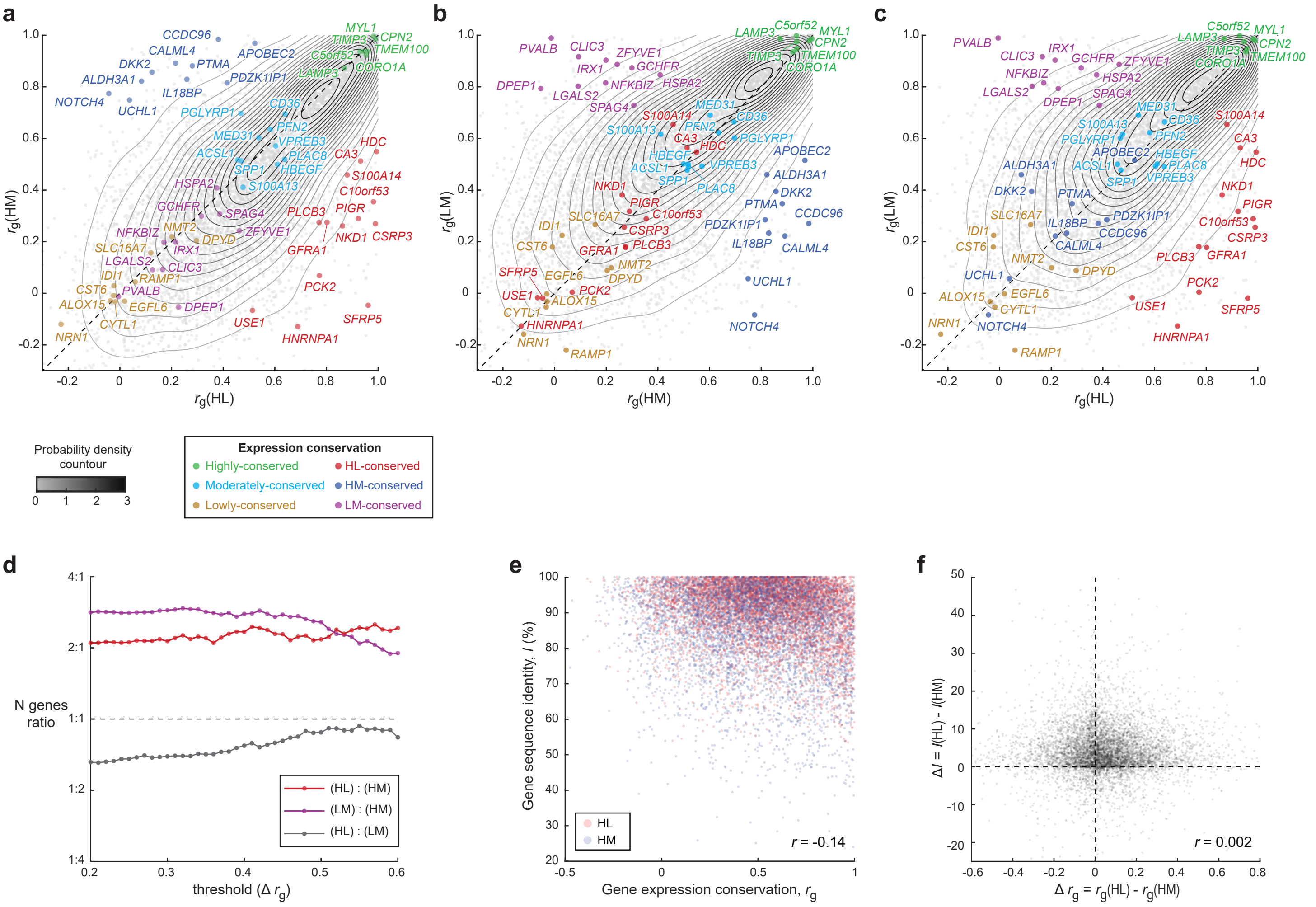

a
b
c
rg(HM)
rg(HL)
rg(LM)
rg(HM)
rg(LM)
rg(HL)
MYL1
CPN2
TIMP3
TMEM100
C5orf52
CORO1A
LAMP3
CCDC96
CALML4
APOBEC2
DKK2
PTMA
ALDH3A1
PDZK1IP1
IL18BP
NOTCH4
UCHL1
PGLYRP1
CD36
PFN2
MED31
VPREB3
ACSL1
PLAC8
SPP1
HBEGF
S100A13
HDC
CA3
S100A14
C10orf53
PIGR
PLCB3
NKD1
CSRP3
GFRA1
PCK2
SFRP5
USE1
HNRNPA1
HSPA2
GCHFR
SPAG4
NMT2
NFKBIZ
DPYD
ZFYVE1
SLC16A7
IRX1
LGALS2
CLIC3
IDI1
RAMP1
CST6
PVALB
ALOX15
EGFL6
DPEP1
CYTL1
NRN1
Probability density countour
0
1
2
3
Expression conservation
Highly-conserved
Moderately-conserved
Lowly-conserved
HL-conserved
HM-conserved
LM-conserved
d
e
f
N genes ratio
4:1
2:1
1:1
1:2
1:4
threshold (Δ rg)
(HL) : (HM)
(LM) : (HM)
(HL) : (LM)
Gene sequence identity, I (%)
Gene expression conservation, rg
HL
HM
r = -0.14
ΔI = I(HL) - I(HM)
Δ rg = rg(HL) - rg(HM)
r = 0.002

Fig. S13

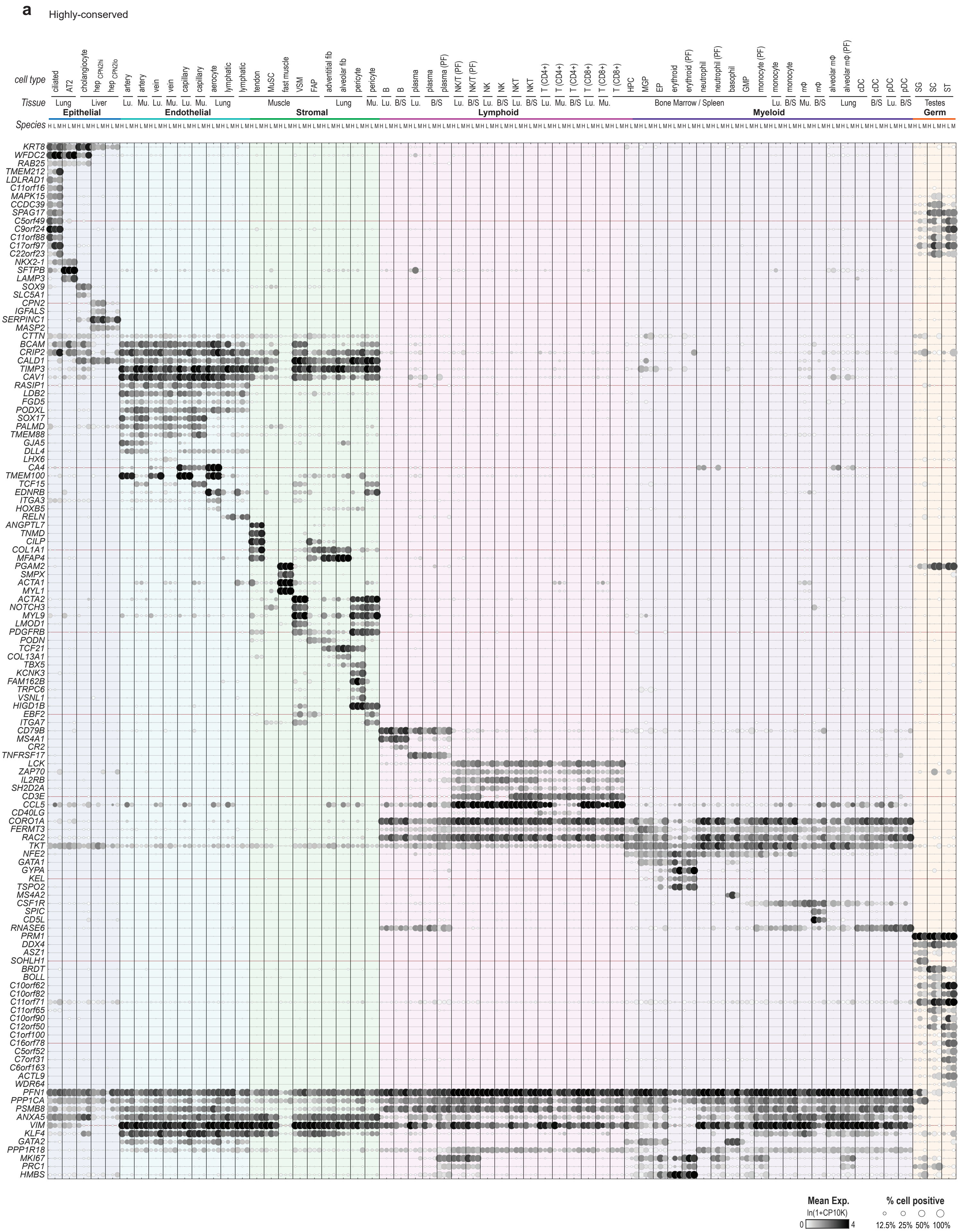
a
Highly-conserved
cell type
Tissue
Species
ciliated
AT2
cholangiocyte
hep CPN2hi
hep CPN2lo
artery
vein
capillary
aerocyte
lymphatic
tendon
MuSC
fast muscle
VSM
FAP
adventitial fib
alveolar fib
pericyte
B
plasma
plasma (PF)
NK/T (PF)
NK
NKT
T (CD4+)
T (CD8+)
HPC
MGP
EP
erythroid
erythroid (PF)
neutrophil
neutrophil (PF)
basophil
GMP
monocyte (PF)
monocyte
mΦ
alveolar mΦ
alveolar mΦ (PF)
cDC
pDC
SG
SC
ST
Lung
Liver
Lu.
Mu.
Muscle
B/S
Bone Marrow / Spleen
Testes
Epithelial
Endothelial
Stromal
Lymphoid
Myeloid
Germ
KRT8
WFDC2
RAB25
TMEM212
LDLRAD1
C11orf16
MAPK15
CCDC39
SPAG17
C5orf49
C9orf24
C11orf88
C17orf97
C22orf23
NKX2-1
SFTPB
LAMP3
SOX9
SLC5A1
CPN2
IGFALS
SERPINC1
MASP2
CTTN
BCAM
CRIP2
CALD1
TIMP3
CAV1
RASIP1
LDB2
FGD5
PODXL
SOX17
PALMD
TMEM88
GJA5
DLL4
LHX6
CA4
TMEM100
TCF15
EDNRB
ITGA3
HOXB5
RELN
ANGPTL7
TNMD
CILP
COL1A1
MFAP4
PGAM2
SMPX
ACTA1
MYL1
ACTA2
NOTCH3
MYL9
LMOD1
PDGFRB
PODN
TCF21
COL13A1
TBX5
KCNK3
FAM162B
TRPC6
VSNL1
HIGD1B
EBF2
ITGA7
CD79B
MS4A1
CR2
TNFRSF17
LCK
ZAP70
IL2RB
SH2D2A
CD3E
CCL5
CD40LG
CORO1A
FERMT3
RAC2
TKT
NFE2
GATA1
GYPA
KEL
TSPO2
MS4A2
CSF1R
SPIC
CD5L
RNASE6
PRM1
DDX4
ASZ1
SOHLH1
BRDT
BOLL
C10orf62
C10orf82
C11orf71
C11orf65
C10orf90
C12orf50
C1orf100
C16orf78
C5orf52
C7orf31
C6orf163
ACTL9
WDR64
PFN1
PPP1CA
PSMB8
ANXA5
VIM
KLF4
GATA2
PPP1R18
MKI67
PRC1
HMBS
Mean Exp.
ln(1+CP10K)
0
4
% cell positive
12.5%
25%
50%
100%

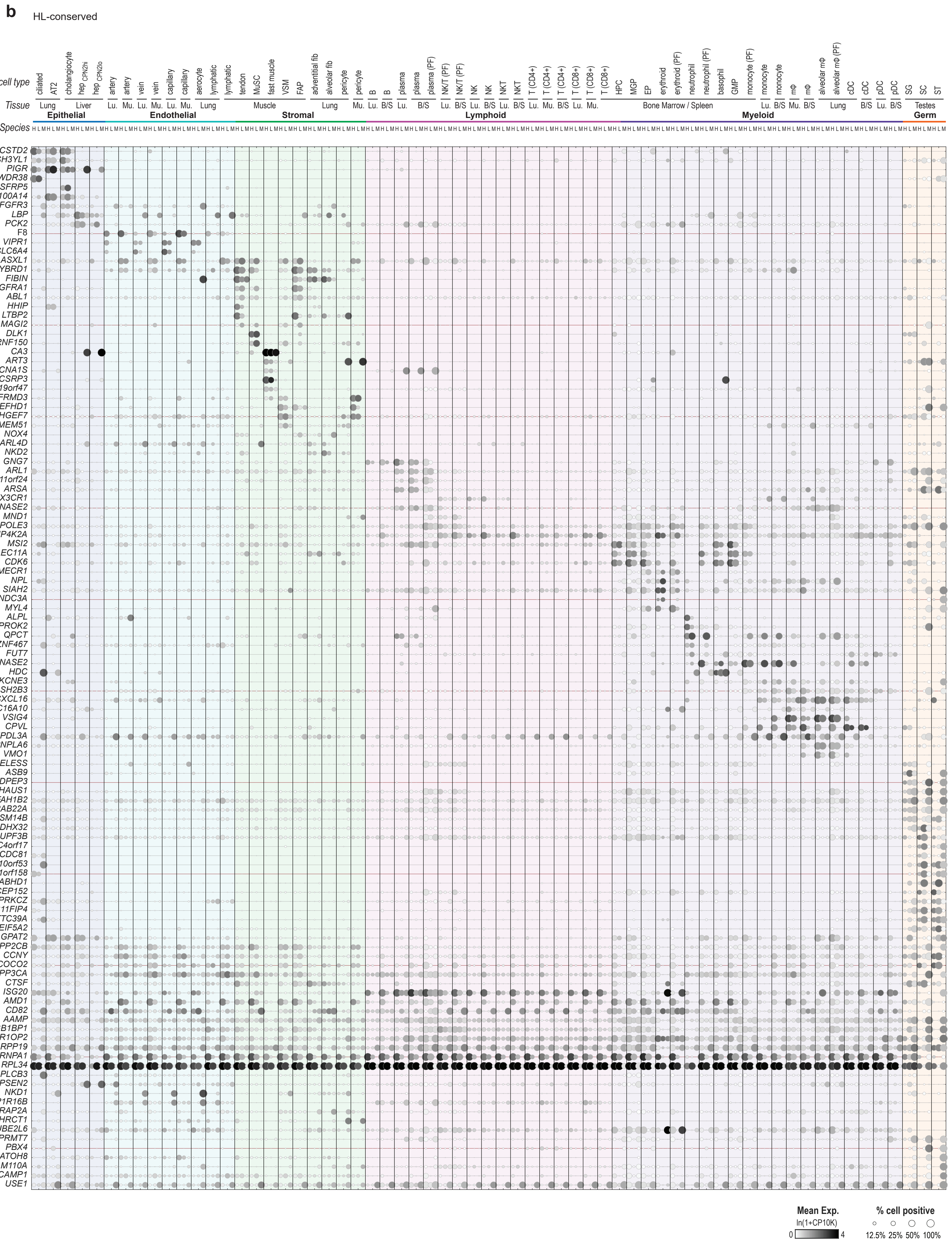
b
HL-conserved
cell type
Tissue
Species
Epithelial
Endothelial
Stromal
Lymphoid
Myeloid
Germ
Mean Exp.
ln(1+CP10K)
% cell positive
12.5% 25% 50% 100%

### Fig. S13 - continued

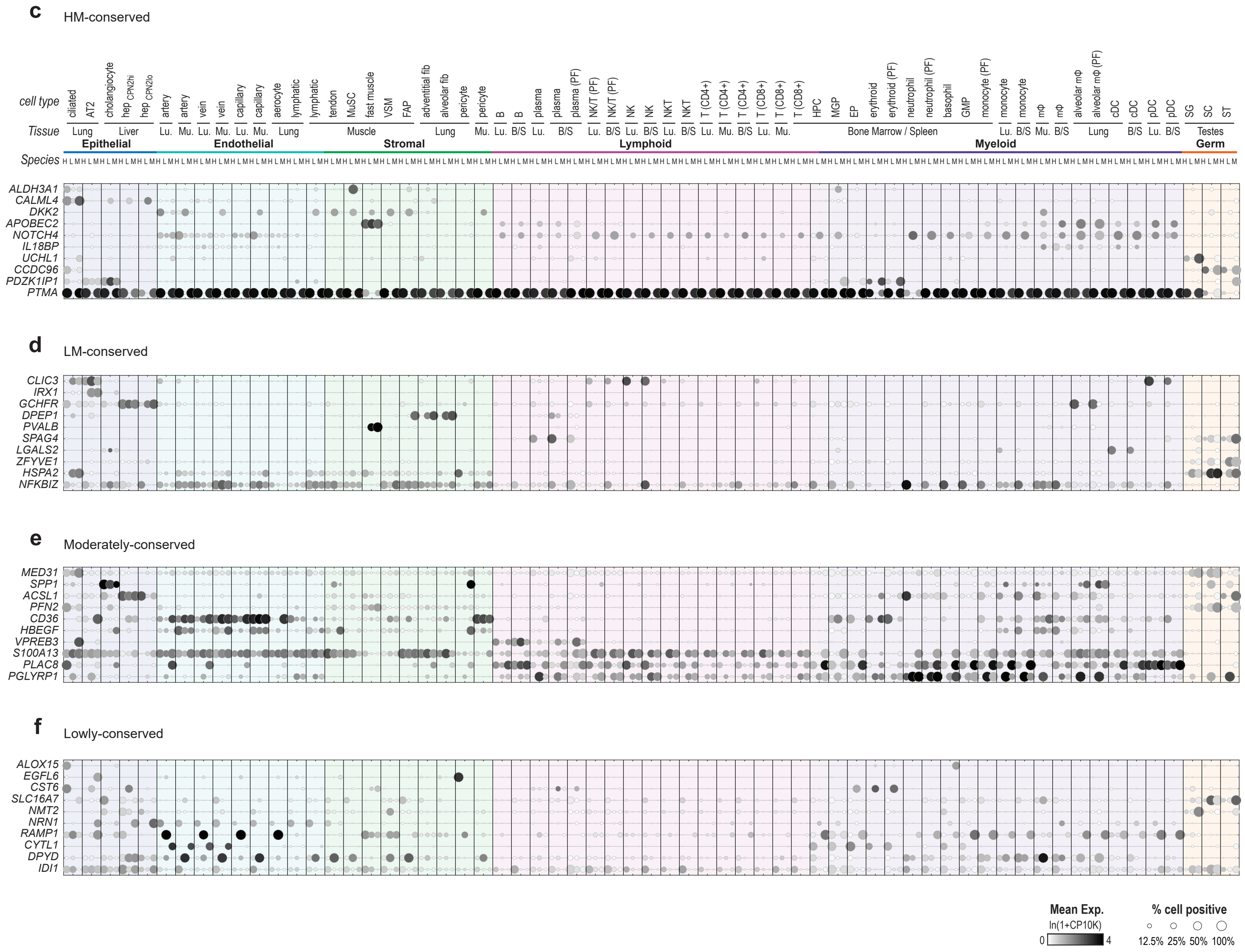
