## Supplementary methods for "Tabula Microcebus: A transcriptomic cell atlas of mouse lemur, an emerging primate model organism"

### Tissue procurement

At the time of euthanasia, blood was immediately collected from each individual via cardiocentesis for sequencing of leukocytes, biobanking, and plasma blood tests, including a complete blood count with cell differential (CBC) (Table S1) and a complete metabolic panel (CMP). See (Casey et al., 2021) for CMP values and the detailed histopathology report of the four lemurs.

**Table S1** - Complete blood count (CBC) and other parameters of lemurs at time of euthanasia.

| | Parameter | Unit | Lemur ID: L1 | L2 | L3 | L4 |
|---|---|---|---|---|---|---|
| At euthanasia | Age | Years | 9.8 | 10.1 | 11.8 | 11.8 |
| | Season (Date of euthanasia) | | LD (06/01) | LD (09/06) | LD (07/24) | LD (08/29) |
| | Number of months in LD prior to euthanasia | | 3 | 6 | 5 | 6 |
| CBC | White blood cells | K/ul | 11.09 | 32.07 ↑ | - | 1.37 ↓ |
| | Red blood cells | M/ul | 9.14 | 4.42 ↓ | - | 7.24 ↓ |
| | Hemoglobin | gm/dL | 14.0 | 7.4 | - | 11.3 |
| | Hematocrit | % | 38.1 | 23.2 | - | 37.6 |
| | Platelet | K/ul | 735 | 237 | - | 39 |
| | Neutrophils | % | 56 ↑ | 91 ↑ | - | 43 ↑ |
| | Lymphocytes | % | 34 ↓ | 4 ↓ | - | 48 ↓ |
| | Monocytes | % | 7 | 5 | - | 6 |
| | Eosinophils | % | 3 | 0 | - | 3 |
| | Basophils | % | 0 | 0 | - | 0 |

↑/ ↓ = higher/lower than reference value according to (Der Goukassian, 1983); SD = short-day season (10:14-h light:dark); LD = long-day season (14:10-h light:dark).
Note the significant leukocytosis and neutrophil left-shift observed in L2 which likely reflects the high inflammatory state of the animal due to uterine cancer metastatic to the lung, Klebsiella pneumonia and metritis with resultant septicemia (more detail in The Tabula Microcebus Consortium, manuscript 2 in prep). The relative thrombocytopenia seen in L4 is the likely cause for the spontaneous pulmonary and renal hemorrhages found on necropsy. CBC values were not obtained for L3 given limited sample that was prioritized for chemistry panel and sequencing.

Organs and tissues were then sequentially collected by a veterinary pathologist. The following delineates the sequential order of organ and tissue harvest for scRNA-seq for each individual:

Lemur #1 (male): blood, lung, frontal cortex of brain (left hemisphere)

Lemur #2 (female): blood, lung, heart and aorta, tongue, trachea, spleen, liver, diaphragm, pancreas, small intestine, colon, kidney, uterus, bladder, brain cortex (right hemisphere) and brainstem, bone and bone marrow, adipose tissue, skin, limb muscle

Lemur #3 (female): blood, heart, spleen, trachea, lung, liver, kidney, uterus, mammary gland, bone marrow. Ovaries and an abdominal mass (presumed mesenteric lymph node tumor) were also harvested for scRNA-seq, however the dissociation protocols were unsuccessful with too few viable cells to proceed with sequencing.

Lemur #4 (male): blood, thymus, heart, pancreas, liver, brain cortex (right hemisphere) and brainstem, hypothalamus and pituitary gland, lung, aorta, tongue, trachea, spleen, diaphragm, adipose tissue, small intestine, colon, kidney, testis, bladder, eye, skin, limb muscle, bone and bone marrow

Organs were processed immediately after collection by tissue experts, except L3's which were processed one to two days post-necropsy due to an unforeseen sudden decline in the animal's physical condition. L3 was found unconscious and unresponsive in her cage during routine daily monitoring, and therefore was urgently euthanized. Necropsied tissues were preserved immediately in HypoThermosol FRS solution (Biolife Solutions #101102) at 4°C overnight. However, because of an unexpected refrigerator malfunction, samples were exposed to 4-16°C for up to 10 hours before downstream processing (i.e., dissociation into single cell suspensions, followed by cDNA and library preparation using the 10x protocol). Despite this prolonged ischemic exposure, expression of immediate early genes (e.g., FOS, FOSB, JUNB, JUBD, EGR1, ZFP36) and those encoding heat shock proteins (e.g., HSPA8, HSPB1) were not significantly higher in L3. In fact, the overall transcriptomic profiles of L3 were similar to that of L1, L2, and L4, with the same cell types clustering together across all animals regardless of individual.

**Histopathological image repository**

Slides from all animals in this study were scanned in collaboration with the Human Pathology and Histology Service Center of the Stanford University School of Medicine Department of Pathology. A plugin (https://github.com/czbiohub/svs-polygon-cropping) for Napari, a multi-dimensional image viewer was used to crop out candidate single images of organs and tissues from each histology slide. The slides can be viewed online at https://tabula-microcebus.ds.czbiohub.org/. See (Casey et al., 2021) for the detailed histopathology report of the four lemurs. The following is a breakdown (in alphabetical order) of all organs and tissues harvested and imaged for each individual:

Lemur #1 (male): adrenal glands, cecum, esophagus, eye, gallbladder, kidney, large intestine, larynx, liver (left stifle joint), lungs, radius/ulna (right-mid) fracture, reproductive tract, salivary gland, scapulohumeral joint (right), skull/nasal cavity, small intestine, spleen, stifle joint (right), stomach, thyroid, tibia/fibula, tongue, trachea, vertebral column

Lemur #2 (female): adrenal glands, aorta, carpus, cecum, eye, heart, kidney, large intestine, larynx, liver, lung, lymph node (tracheobronchial), pancreas, scapula/elbow (sagittal), skeletal muscle, skin, skull/nasal, small intestine, spine, spleen, sternum, stifle joint (sagittal), stomach, tarsus, thymus, thyroid, tongue, trachea, urinary bladder, uterus

Lemur #3 (female): abdominal mass, adrenal glands, aorta, cecum, cervix, colon (proximal), ears, esophagus, eyes, heart, kidney, liver, lungs, nasal cavity, nasal/mandible/maxilla, ovary, rectum, ribs, salivary glands, small intestine, spine, spleen, stifle joint (right and left), stomach, thyroid, tibia (right and left), tongue, trachea, uterus, vagina

Lemur #4 (male): aorta, brain (left hemisphere), cecum, colon (proximal and distal), duodenum, esophagus, eye, heart, ileum, jejunum, kidney, larynx, liver, lumbar, lung, male reproductive tract, nose, pancreas, pericardial tissue, sacral vertebrae), skull, spine (cervical, spleen, stifle joint, stomach, thoracic, thymus, thyroid, tongue

**Preparation of single cell suspensions and FACS-sorting for scRNA-seq**

Lysis plates for flow sorted cells were prepared as previously described (Tabula Muris Consortium, 2020; Tabula Muris Consortium et al., 2018) by dispensing lysis buffer into each well (0.4 ul and 4 ul for 384-well and 96-well plates, respectively) using a Mantis liquid handler (Formulatrix). Plates were then sealed, centrifuged, flash frozen on dry ice, and stored at -80°C until used for FACS.

FACS sorting with Sony (SH800s) or Becton Dickinson (Aria III) sorters was performed on cell suspensions to isolate single cells into 384- or 96-well plates. Tissue dissociation and single cell suspension protocols are detailed below for each tissue separately. The majority of tissues were sorted into 384-well plates, except the following were sorted into 96-well plates: blood, brain, and lung for L1; hand-picked cardiomyocytes for L2; blood for L4 (the latter of which was also sorted into 384-well plates). No sorting was done for L3.

Sorters were calibrated for dispensing accuracy before cell loading and repeated after every 8-10 sorted plates. Up to 5 mL of single cell suspension was filtered, gently vortexed and then loaded onto the FACS machine. Pressure adjustments were employed to check cell concentration and suspensions diluted with FACS buffer (2% FBS in PBS) or plain PBS as needed. Single cells were selected by forward scatter, with initial gating set to exclude cell debris and platelet aggregates. Cells were sorted using the highest purity setting to enrich for single cells. Sytox blue nucleic acid stain (ThermoFisher #S34857) was used for all samples to identify and exclude dead cells.

After FACS, plates were immediately sealed and labeled, spun down, and snap frozen on dry ice. Sorting was typically completed for each 384-well plate within 6-10 minutes, and for each 96-well plate in <5 minutes. FACS-sorting was performed only for tissues in the SS2 pipeline, and not for any samples in the 10x pipeline (see directly below).

**Library preparation, quality control, and sequencing**

**10x Genomics**

Single cells were isolated and analyzed with the 10x Genomics single cell RNA-sequencing pipeline (Chromium Single Cell 3’ Library and Gel Bead v2 Chemistry kit) as previously described (Tabula Muris Consortium, 2020; Tabula Muris Consortium et al., 2018;

Travaglini et al., 2020). Based on the concentration of the cell suspension (determined after tissue dissociation by manual counting using a hemocytometer or by automated cell counter), samples were concentrated or diluted with 2% FBS in PBS to a target concentration of about 10^6 cells/mL. Then, cell suspensions were loaded directly into microfluidic chips (~5000 cells per sample) on a 10x Chromium Controller, which facilitated single cell capture, followed by reverse transcription to cDNA and barcoding within droplet emulsions. For cDNA amplification, 12 PCR cycles were used (Biorad C1000 Touch Thermal-cycler). Quality of the amplified cDNA was determined on a 12 or 96 capillary Fragment Analyzer using the High Sensitivity NGS Analysis Kit (Advanced Analytical). Similarly, the quality and average fragment length of cDNA libraries were quantified by capillary-based fragment analysis. After this, library concentration was determined by qPCR using the KAPA Library Quantification Kit (Roche) optimized for Illumina sequencing. Each library was normalized to 2 nM and then equal volumes of up to 16 libraries were combined together to constitute the final sequencing sample pool. A PhiX control library was spiked in at 0.2% to 1%. Library pools were sequenced on S2 flow cells of the NovaSeq 6000 System (Illumina) using 100 cycle reagent kits (Illumina, 20012862) with the following Read and Index barcode lengths: Read 1 (26 bp), Read 2 (90 bp), Index (8 bp).

**Smart-seq2 (SS2)**

As previously performed in the Tabula Muris (Tabula Muris Consortium, 2020; Tabula Muris Consortium et al., 2018), the Smart-seq2 protocol (Ramsköld et al., 2012; Wu et al., 2014) was employed for cDNA synthesis and library construction. FACS plates were thawed on ice, spun down, and then used for first strand synthesis. Master mixes were prepared with identical reagents at the same concentrations and dispensed into individual wells (0.6 ul for 384-well, 6 ul for 96-well plates) using a Mantis liquid handler (Formulatrix). The reverse transcription step was carried out at 42°C for 90 min and terminated by heating at 72°C for 5 min on a Biorad C1000 Touch Thermal-cycler. For second strand synthesis, PCR master mixes were prepared in the same manner as our prior Tabula projects. 1.5 ul and 15 ul of master mix were robotically added into each well (384-well and 96-well plate, respectively). DNA preamplification was performed on a Biorad thermal-cycler utilizing the following program: Step 1 (37°C for 30 min), Step 2 (95°C for 3 min), Step 3 (93°C for 20 sec, 67°C for 15 sec, 72°C for 4 min), Step 4 (72°C for 5 min). Step 3 was repeated for 21 cycles (96-well plates) or 23 cycles (384-well plates).

cDNA purification for 384-well plates was identical to our prior work (Tabula Muris Consortium, 2020; Tabula Muris Consortium et al., 2018). For 96-well plates, cDNA purification was performed with AMPure XP beads (Beckman Coulter #A63881) in the following manner. 18 ul of beads (0.7x) were added to each well by multichannel pipetting. Plates were sealed, gently vortexed, spun down, and then incubated at room temperature for 10 min. Beads were sequestered using a magnet until the supernatant was clear followed by decantation. Two rounds of washing were then performed (170 ul of 80% ethanol for each well). After removal of ethanol, plates were allowed to dry at room temperature for 5 to 15 min. Subsequently, beads were resuspended in 16 ul of EB elution buffer (Qiagen #19086), incubated for 5 min, and finally eluted.

cDNA quality was assessed by capillary electrophoresis on a Fragment Analyzer (AATI) for all 384- and 96-well plates. Candidate wells were selected for downstream processing if their cDNA concentrations were at least one standard deviation beyond the mean concentration of the blanks and the size range of their cDNA was consistent with the size spectrum of RNAs expected for a primate species (500 – 5000 bp). Samples from 96-well plates were reformatted into 384-

well plates at a target concentration between 0.05 ng – 0.16 ng/ul using a Mosquito X1 single channel pipetting robot (TTP Labtech), with dilutions in EB buffer performed using a Mantis liquid handler.

SS2 sequencing libraries were prepared for the Illumina platform as described previously (Darmanis et al., 2015; Tabula Muris Consortium, 2020; Tabula Muris Consortium et al., 2018) using a starting cDNA volume of 0.4 ul. For L1 and L2, tagmentation, neutralization, and indexed PCR reactions (with dual unique i5 and i7 indexing primers) were performed using the Nextera XT Library Sample Preparation Kit (Illumina #FC-131-1096). Tagmentation was carried out at 55°C for 10 min followed by neutralization at room temperature for 5 min. Thermo-cycler settings for PCR amplification entailed: Step 1 (72°C for 3 min), Step 2 (95°C for 30 sec), Step 3 (95°C for 10 sec, 55°C for 30 sec, 72°C for 1 min; repeated 12x), Step 4 (72°C for 5 min). For L4, an in-house library preparation protocol was employed and optimized, using a different source of Tn5 enzyme (Quantitative Biosciences QB3 MacroLab core facility, UC Berkeley). No significant differences in tagmentation size, library quality, and sequencing read depth were seen between libraries made using our in-house protocol versus the Nextera kit.

Library pooling and quality control were performed as previously detailed (Tabula Muris Consortium, 2020; Tabula Muris Consortium et al., 2018). After library construction, wells of each library plate were pooled initially using a Mosquito liquid handler (TTP Labtech), followed by hand pipetting into a single microcentrifuge tube (one tube per library plate). Two rounds of purification using AMPure XP beads (0.8x followed by 0.7x) were then performed involving two washes with 80% ethanol for each purification, followed by elution (100 ul elution buffer for the first round and 40 ul for the final elution). Library quality and the average fragment size were then measured by capillary electrophoresis on a Fragment Analyzer (AATI) and library concentrations subsequently quantified by qPCR (Kapa Biosystems #KK4923) on a CFX06 Touch Real-Time PCR Detection System (Biorad). Each library plate pool was normalized to 2 nM (L1 and L2) or 4 nM (L4) and equal volumes for up to 20 plates were mixed together to make the final sequencing sample pool. A PhiX control library was spiked in at 0.2% prior to sequencing.

Libraries were sequenced on a NovaSeq 6000 System (Illumina) using 2 x 100 bp paired-end reads and 2 x 12 bp dual unique index reads with 200 or 300 cycle kits designed for either S2 or S4 flow cell configurations (Illumina, 20012861, 20012860, 20027466, 20012866).

**Genome alignment**

Individual chromosome scaffolds of the *Microcebus murinus* genome assembly (Mmur 3.0, Refseq assembly accession: GCF_000165445.2 available from NCBI https://www.ncbi.nlm.nih.gov/assembly/GCF_000165445.2/) were concatenated together in Linux to create a whole genome FASTA file. The gene annotation file was retrieved from the NCBI Microcebus murinus Annotation Release 101 FTP site. For tRNA genes, we summed all identically named tRNA gene loci (e.g. 'TRNAA-AGC-1', 'TRNAA-AGC-2', etc. into one ‘gene’, 'TRNAA-AGC'). This resulted in a total of 31,509 genes.

10x alignment as described in Main Methods. For SS2 alignment, the following options were used for STAR mapping: --alignIntronMin 20, --alignIntronMax 1000000, --alignMatesGapMax 1000000, --alignSJoverhangMin 8, --alignSJDBoverhangMin 1, --outBAMcompression 10, --outFilterMultimapNmax 20, --outFilterMismatchNmax 999, --outFilterMismatchNoverReadLmax 0.04, --outFilterType BySJout, --outSAMattributes NH HI AS NM MD, --outSAMstrandField intronMotif, --outSAMtype BAM SortedByCoordinate, --

outSAMunmapped Within, --outSJfilterReads Unique, --quantMode TranscriptomeSAM GeneCounts, --quantTranscriptomeBAMcompression 10, --sjdbOverhang 99, --sjdbScore 1

SS2 reads were first pre-processed with skewer (version 0.2.2), an adapter trimmer optimized for Illumina paired-end reads. Then, STAR alignment was performed, with first pass mapping to identify splice junctions. These junctions were subsequently incorporated to generate a new genome, after which second pass mapping was initiated. A remapped BAM file with second pass mapping statistics was created using samtools. FPKM was calculated with cuffquant, and gene expression estimated using RSEM (version 1.3.1). Read quality was then assessed with HTSEQ, and the count tool used to sum up for each gene the total number of aligned reads overlapping its exons, generating a gene counts table.

**Cell filtering**

For all tissues except the heart the final cutoff was 500 genes, 1000 UMI (10X), 5000 reads (SS2). For the heart the cutoff was 50 genes, 100 UMI (10X), and 5000 reads (SS2).

**FIRM integration**

The FIRM algorithm can integrate multiple scRNA-seq datasets by iteratively integrating two datasets together. The tissue-level integrated datasets (https://tabula-microcebus.ds.czbiohub.org/organs) were constructed by first integrating via the FIRM algorithm the tissue datasets of the same individual that were sequenced by different methods (10x or SS2), and then further integrating datasets from different lemur individuals. This created a single object for each tissue that combines scRNA-seq data from each individual and sequencing method.

The atlas-wide integrated dataset was created by first integrating datasets across individuals, then tissues, and lastly sequencing methods. Typically, tissues with more overlapped highly variable genes were integrated first. Because testes cells (predominantly germ cells) showed the most distinctive transcriptional profiles compared with cells of the other tissues, the testes dataset was first separately integrated across different sequencing methods and then integrated with the rest of the tissues.

When applying FIRM, the top 4000 highly variable genes for each of the two dataset were identified using the "FindVariableFeatures" function in Seurat. FIRM then used the intersection, union, or the top ranked genes of the detected highly variable genes of the two datasets for principal component analysis (method chosen according to best performance). For dimensionality reduction, the number of principal components was chosen as the larger number of principal components used for dimensionality reduction of the two pre-integration datasets.

**Comparison of expression profiles among mouse lemur cell types - differentially expressed genes (related to Fig. 5c-e)**

The Wilcoxon rank-sum test was used to identify differentially expressed genes between groups of cell types. To search for genes that were preferentially expressed in the sperm/sperm progenitor cells and immune progenitor/proliferating cells in comparison to cells of other compartments (epithelial, endothelial, stromal), we first applied a right-tailed hypothesis rank-sum test to genes that were expressed at a moderate level (>0.3 in relative expression level, natural log transformed and normalized to the cell type maximal expression level across the atlas) in at least one cell type in the sperm/sperm progenitor cells and in immune progenitors. These genes were then tested to remove the genes that were not differentially expressed between sperm/sperm progenitor cells versus immune progenitor/proliferating cells. The Bonferroni method was used for multiple testing correction of all *p*-values. Genes that passed all criteria in the top 1% of *p*-values were manually examined to confirm consistency of expression. The same workflow was performed to search for genes preferentially expressed in spermatogonia and immune progenitor/precursor cells in comparison to proliferating cells of other compartments (i.e., epithelial, stromal, neural), in Schwann cells and stromal cells in comparison to cells of the neural compartment, as well as in glial cells of the central nervous system in comparison to Schwann cells and stromal cells.

### TISSUE DISSOCIATION PROTOCOLS

### NERVOUS SYSTEM

*Brain cortex, brainstem, hypothalamus, and pituitary gland*

After removal from the skull, cerebral tissue was placed directly in ice cold Hibernate A medium (Gibco #A1247501). The relevant brain regions were microdissected for the following animals: L1, left frontal cortex; L2, entire right cortex and entire right brainstem; and L4, entire right cortex and entire right brainstem with the hypothalamus and pituitary gland dissected apart from the brain. Each region was mechanically minced into ~1 $mm^3$ pieces with a scalpel, and then digested under gentle agitation in Hibernate A minus Calcium (BrainBits #HACA) containing 20 U/mL Papain (Worthington #LS003126) at 34°C for 1 hour (except for the pituitary gland which was dissociated for 10 min). The digested homogenate was subsequently washed with 10% FBS and 10 mM HEPES in cold L-15 medium (Gibco #11415), and triturated 10 times using a P1000 pipette, and then 10 more times with a 500 µm sterile glass pipette. Next, the cell suspension was overlaid on the top surface of a Percoll solution containing 1 mL of Percoll (Sigma #P1644) and 4 mL of L-15 and 10 mM HEPES. The gradient was centrifuged at 400g, 4°C for 9 minutes and the solution decanted except for a 1 mL layer at the bottom. This layer was spun down at 400g, 4°C for 4 min, and the resulting cell pellet resuspended in buffer consisting of phenol-red free L-15 (Gibco #21083027), 10 mM HEPES, 0.5% BSA, and 0.01% DNase (Sigma #DN25). The cell suspension was filtered through a sterile 100 µm strainer (Falcon #352360). Concentration was determined by manual cell counting on a hemocytometer and adjusted to 10^6 cells/mL. A portion of the sample was aliquoted for 10x processing with the remaining cells stained with Sytox blue (ThermoFisher #S34857) prior to FACS-sorting for SS2 processing. Due to a shortage of reagent during tissue processing of L4, Hibernate A minus Calcium was used in place of L-15 for all the protocol steps detailed above.

*Eye Retina*

Extracted eye tissue was placed in cold DMEM (Gibco #11054020). The sclera, choroid, and retinal pigment epithelial (RPE) layers were separated from the retina manually and mechanically using microsurgical instruments. The retinal sample was further dissociated in 0.25% trypsin-EDTA (Gibco #12605010) at 37 °C for 30 minutes, and detached cells collected every 5 mins in cold DMEM containing 10% FBS.  Cell suspensions were resuspended in PBS containing 10% FBS, then filtered through a 100 µm nylon cell strainer (Falcon #352360). Concentration was determined by manual cell counting on a hemocytometer and adjusted to 10^6 cells/mL. A portion of the sample was aliquoted for 10x processing with the remaining cells stained with Sytox blue (ThermoFisher #S34857) prior to FACS-sorting for SS2 processing. Dissection of RPE cells was performed as previously described (Fernandez-Godino et al., 2016), however the sample was not sequenced due to low viable cell count.

### INTEGUMENTARY SYSTEM

#### *Skin*

A similar dissociation protocol was employed in the Tabula Muris (Tabula Muris Consortium, 2020; Tabula Muris Consortium et al., 2018). After extracting the skin, the underlying subcutaneous fat was removed with a scalpel. The remaining skin layer was then incubated with gentle agitation in digestion buffer consisting of 0.25% trypsin (Thermo #15050057) and 15% FBS in DMEM (Gibco #11054020) at 37°C for 30 min. The top layer of epidermal cells was scraped away from the dermis, releasing keratinocytes in a single cell suspension. After an additional digestion in trypsin for 5 minutes, cells were filtered in succession through 70 µm and 40 µm cell strainers (Falcon #352350, #352340, respectively) on ice, then washed with cold 5% FBS in PBS. Cells were spun down at 500g, 4°C for 5 min and the final pellet resuspended in cold 5% FBS in PBS. Concentration was determined by manual cell counting on a hemocytometer and adjusted to 10^6 cells/mL. A portion of the sample was aliquoted for 10x processing with the remaining cells stained with Sytox blue (ThermoFisher #S34857) prior to FACS-sorting for SS2 processing.

#### *Fat*

A similar dissociation protocol was employed in the Tabula Muris (Tabula Muris Consortium, 2020; Tabula Muris Consortium et al., 2018), aimed at extracting the stromal vascular fraction (SVF) from adipose tissue. After extraction, fat tissues including inguinal subcutaneous adipose tissue (SCAT, L2 and L4), perigonadal (GAT, L2 only), mesenteric (MAT, L2 and L4), and interscapular brown (BAT, L2 and L4), were individually dissected out and minced using fine surgical instruments in a petri dish on ice until forming a slurry. Tissue was then digested under continuous gentle shaking at 37°C for 30 minutes in Ham's F-10 medium (Gibco #11550043) containing 760 U/mL collagenase II (Worthington #LS004177) and 1 U/mL Dispase II (Gibco #17105-041). After trituration, cells were filtered sequentially through 100 µm (Falcon #352360) and 40 µm (Falcon #352340) strainers on ice and then washed with cold Ham's F-10 medium containing 10% horse serum (Invitrogen #16050114). Concentration was determined by manual cell counting on a hemocytometer and adjusted to 10^6 cells/mL for 10x processing. Note that the perigonadal fat was collected from L2 (female) and surrounding tissue was likely sampled (including from the ovary), given that granulosa cells and mesothelial cells were found in the scRNA-seq dataset. L4's fat samples were combined for 10x sequencing due to limited space on the microfluidic chips.

#### *Mammary gland*

L3's mammary gland tissue was placed at 4°C in HypoThermosol FRS solution (Biolife solutions #101102) for 1 day. Subsequently, the gland tissue was placed in 10 mL of cold DMEM/12 medium (Gibco #12634010) and directly minced with a sterile razor blade into approximately 1 mm fractions. The samples were then incubated under gentle shaking in digestion buffer (Stem Cell Technology #07912) consisting of 150 U/mL collagenase and 50 U/mL hyaluronidase in DMEM at 37°C for 2 hrs, interspersed by mild pipetting every 30 minutes to promote further tissue dissociation. The digested homogenate was subsequently spun down at 1500 rpm, followed by treatment with 5 mL of ACK lysing buffer (Lonza #10-548E) on ice for 5 minutes to remove red blood cells. Two additional digestion steps were then performed in succession, first with 5 mL of 0.25% trypsin-EDTA (Invitrogen #252001144) for up to 5 min, followed by a shorter digestion with DNase I (Worthington #LS002139). Dissociated cells were then filtered through a sterile 40 μm strainer (Falcon #352340) and washed with buffer containing 1% BSA and 2% FBS in HBSS. Concentration was determined by manual cell counting on a hemocytometer and adjusted to 10^6 cells/mL for 10x processing.

### MUSCULOSKELETAL SYSTEM

*Limb muscle and Diaphragm*

Muscle was extracted from the hind limb below the knee. Diaphragm was isolated in one piece by cutting along the inner side of the ribcage. After extraction, each sample was washed in cold wash media containing Ham's F-10 (HyClone #SH30025.01), 10% horse serum (Invitrogen #16050114), and 1% pen/strep (Omega Scientific #PS-20), and then gently dried with Kimwipes. The samples were further mechanically dissociated with surgical scissors, and subsequently digested under gentle continuous shaking at 37°C for 35 minutes in dissociation buffer (0.2% (w/v) collagenase II (Worthington #LS004177) in wash media). Cells were spun down at 4°C, 1600g for 5 minutes, the supernatant decanted and the pellet further digested in PBS containing 1% Dispase (Gibco #17105-041) and 0.5% collagenase II. After vortexing, the samples were incubated at 37°C for 20 min under gentle continuous shaking. Post-digestion, cell suspensions were pulled five times through a 10 mL syringe (BD #309604) using a 20-gauge needle (Fisher #305175), and then filtered through a 40 μm nylon cell strainer (Falcon #352340). The samples were washed and spun down for 5 min at 1600g, decanted, and resuspended in wash media and filtered in 5 mL filter-cap tubes (Falcon #352235). Concentration was determined by manual cell counting on a hemocytometer and adjusted to 10^6 cells/mL. A portion of the sample was aliquoted for 10x processing with the remaining cells stained with Sytox blue (ThermoFisher #S34857) prior to FACS-sorting for SS2 processing. Some cells were also stained with anti-human antibodies, NCAM1/CD56 (Biolegend #304628), THY1/CD90 (BioLegend #328114), ITGB1/CD29 (Biolegend #303007), and Sytox blue, and immediately FACS-sorted for functional experiments (see de Morree et al, accompanying manuscript).

*Bone*

Bones were first processed to collect bone marrow (see below). The unfiltered residual bone tissue was further minced with a razor blade and resuspended in digestion buffer supplemented with 3000 U/mL collagenase II (Sigma-Aldrich #C6885) and 100 U/mL DNase I (Worthington #NC9199796). The sample was incubated at 37°C for 1 hour under gentle agitation, then filtered through a 70 μm nylon mesh strainer (Falcon #352350) and the digestion reaction quenched with 2% FBS in PBS (Gibco #C14190500BT). The skeletal cells were separated from red blood cells and bone dust by density gradient separation using 1:1 of room temperature Ficoll Histopaque 1.119 g/mL density gradient media (Sigma-Aldrich #11191). The buffy coat was collected, washed with 2% FBS in PBS. Concentration was determined by manual cell counting on a hemocytometer and adjusted to 10^6 cells/mL. A portion of the sample was aliquoted for 10x processing with the remaining cells stained with Sytox blue (ThermoFisher #S34857) prior to FACS-sorting for SS2 processing.

### CIRCULATORY SYSTEM

*Bone marrow*

After bone harvesting (humerus and spine), a paper towel was used to remove any residual fibers and muscle. A mortar and pestle were used to crush the bone and extract the marrow. Several mLs of 2% FBS in PBS containing 1:100 DNase I (Worthington #LS006343) were added to the mortar to keep cells alive and prevent clumping of cells. Crushing of bone continued until no further red fluid leaked from the dissociated tissue. The sample was then filtered through a 40 µm strainer (Falcon #352340) into a 50 mL Falcon conical tube containing 2% FBS in PBS  and DNA I placed on ice. The unfiltered residue was collected and processed for bone cells (see above). The filtrate was spun down at 4°C, 300g for 5 minutes, and resuspended in 2% FBS in PBS and filtered through a 70 µm strainer (Fisherbrand #22363548). Concentration was determined by manual cell counting on a hemocytometer and adjusted to 10^6 cells/mL. A portion of the sample was aliquoted for 10x processing with the remaining cells stained with Sytox blue (ThermoFisher #S34857) prior to FACS-sorting for SS2 processing. Note, L3’s bone marrow was processed on the day of the euthanasia using protocol above, but the single cell solution was suspended in Bambanker solution (Fisher Scientific #NC9582225) and frozen at -80°C using a cell freezing container for adequate cryopreservation. On the day of 10x processing, cells were thawed in 37°C water bath and then resuspended in 2% FBS in PBS solution at the appropriate cell concentration.

*Blood*

After cardiocentesis of the left ventricle, blood was gently aspirated through a fine surgical needle and collected in EDTA-coated tubes. The sample was diluted 1:10 with 2% FBS in PBS, and additional DNase I (Worthington #LS006343) supplemented if blood started to coagulate. Diluted room temperature blood was then gently layered, without mixing, into a 15 mL sterile Falcon conical tube containing half of the diluted sample volume of room temperature Ficoll Histopaque 1.119 g/mL density gradient media (Sigma-Aldrich #11191), forming 1 part Ficoll solution at the bottom and 2 parts sample at the top. Using the higher density Ficoll mix compared to the standard Ficoll density (1.077 g/ml) allows for polymorphonuclear cells to be separated, in addition to mononuclear cells. The sample was spun down at room temperature, 400g for 30 minutes with the brake setting off. The leukocyte-rich buffy coat was then carefully transferred with use of a pipette to a new 15 mL Falcon tube. Cold 2% FBS in PBS was added to fill up the tube and the solution was mixed by inverting. The sample was then spun down at 4°C, 500g for 5 minutes (prior to this, all steps had been carried out at room temperature). After removing the supernatant, cells were resuspended in 1 mL cold 2% FBS in PBS. The single cell solution was gently pipetted up and down to further dissociate any cell clumps. If cells persisted to clump, 0.1 mg/mL of DNase I was added to the sample, incubated on ice for 10 min, and then washed (spun down at 4°C, 500g for 5 minutes). If the sample remained red/pink colored after Ficoll (indicating presence of red blood cells), it was diluted 1:10 with 1X BD Pharm Lyse, a red blood cell lysing buffer (BD Biosciences #555899), incubated on ice for 5-10 min until the cloudy solution turned transparent red, and then washed (spun down at 4°C, 500g for 5 minutes). Cells were filtered through a 70 µm strainer (Fisherbrand #22363548). Concentration was determined by manual cell counting on a hemocytometer and adjusted to 10^6 cells/mL. A portion of the sample was aliquoted for 10x processing with the remaining cells stained with

Sytox blue (ThermoFisher #S34857) prior to FACS-sorting for SS2 processing.  Note, L3's blood was processed on the day of the euthanasia using protocol above, but the single cell solution was suspended in Bambanker solution (Fisher Scientific #NC9582225) and frozen at -80°C using a cell freezing container for adequate cryopreservation. On the day of 10x processing, cells were thawed in 37°C water bath and then resuspended in 2% FBS in PBS solution at the appropriate cell concentration. L4's blood clotted after collection, and so the red blood cell lysis was done prior to the Ficoll separation, in addition to use of DNase I, in order to dissociate the clot as much as possible.

*Spleen*

After extraction, splenic tissue was washed in 1x HBSS (ThermoFisher #14175), minced with razor blades, and further dissociated mechanically using the flat-end of a plunger. The sample was then triturated sequentially through 100 µM, 70 µM, and 40 µM strainers (Falcon #352360, #352350, #352340, respectively). Cells were washed in 2% FBS in PBS followed by ACK treatment (Gibco #A10492-01) at room temperature for 5 minutes. Residual debris was removed using debris removal solution (Miltenyi Biotec #130-109-398) per factory instructions. Cells were resuspended in 2% FBS in PBS. Concentration was determined by manual cell counting on a hemocytometer and adjusted to 10^6 cells/mL. A portion of the sample was aliquoted for 10x processing with the remaining cells stained with Sytox blue (ThermoFisher #S34857) prior to FACS-sorting for SS2 processing. Note, L3's extracted splenic tissue was first preserved at 4°C for 2 days in HypoThermosol FRS solution (Biolife Solutions #101102) before continuing on with the above delineated steps.

*Thymus*

Thymic tissue was crushed with a 70 µm strainer (Falcon #352350) and centrifuged at 5°C, 270g for 5 minutes. The cell pellet was digested with 2.2 mg/mL collagenase II (Sigma #C6885) at 37°C for 10 min, then incubated under gentle agitation at 37°C for 30 more minutes. Digestion was quenched with FACS buffer containing 2% FBS, 1% antibiotics (Gibco #15240-062), and 10% pluronics (ThermoFisher #24040032) in PBS. The cell suspension was pelleted at 5°C, 270g for 5 min, then washed, resuspended in FACS buffer, and stained with Sytox blue (ThermoFisher #S34857) prior to FACS-sorting for SS2 processing.

*Heart*

After extraction, cardiac tissue was placed on ice in a sterile 60-mm polystyrene culture dish (Thermofisher #174888) containing perfusion buffer (135 mM NaCl, 4 mM KCl, 1 mM $MgCl_2$, 10 mM HEPES, 0.33 mM $NaH_2PO_4$, 10 mM glucose, 10 mM 2,3-butanedione monoxime, 5 mM taurine; overall pH 7.2) to inhibit myosin activity and induce cardioplegia. The 4 cardiac chambers were further dissected apart from each other (left atrium, LA; left ventricle LV, right atrium, RA; right ventricle, RV) using fine surgical forceps and scissors. Each chamber fraction was then moved to individual 15 mL sterile conical tubes (Falcon) containing digestion buffer (0.3 mg/g body weight collagenase D, Roche #11088858001; 0.4 mg/g body weight collagenase B, Roche #11088807001; 0.05 mg/g body weight Protease type XIV, Sigma #P5147; in sterile water), and incubated with continuous gentle shaking at 37°C for 10-20 minutes. During this period, samples were gently pipetted up and down two times to facilitate dissociation. After trituration of the majority of tissue into a single cell suspension, the sample was filtered through a 70 µm nylon mesh strainer (Falcon #08-771-1). The concentration of the eluent (non-myocyte fraction) was determined by manual cell counting on a hemocytometer and adjusted to 10^6,  and subsequently divided into 2 aliquots: one for the 10x processing and the second stained with Sytox blue (ThermoFisher #S34857) prior to FACS-sorting for SS2 processing. The fraction caught by the mesh strainer (cardiomyocyte fraction) was handled similarly, but divided into 3 aliquots (10x, SS2, and from the remaining fraction cardiomyocyte cells were hand-picked under light microscopy with a 20 µl pipette and deposited cell by cell into a hard-shell 96-well plate (BioRad #HSP9601) for further SS2 processing (without FACS-sorting). Note, L3's extracted cardiac tissue was first preserved at 4°C for 1 day in HypoThermosol FRS solution (Biolife Solutions #101102) before continuing on with the above delineated steps.

*Aorta*

After extraction, aortic tissue was minced using sterile microsurgery tools, then digested with 2.2 mg/mL collagenase II (Sigma #C6885) at 37°C for 10 min, and incubated under gentle agitation at 37°C for an additional 30 minutes. The digestion reaction was quenched with FACS buffer containing 2% FBS, 1% antibiotics (Gibco #15240-062), and 10% pluronics (ThermoFisher #24040032) in PBS. Cells were pelleted at 5°C, 270g for 5 min, and resuspended and washed in FACS buffer. Concentration was determined by manual cell counting on a hemocytometer and adjusted to 10^6 cells/mL for 10x processing.

### RESPIRATORY SYSTEM

*Lung*

Tissue samples were processed into single cell suspensions in a manner similar to the protocol used in Tabula Muris (Tabula Muris Consortium, 2020; Tabula Muris Consortium et al., 2018). After extraction of both right and left lungs from the thoracic cavity, the tissue was triturated using sterile microsurgery tools and placed in gentleMACS c-tubes (Miltenyi #130-096-334) with digestion buffer consisting of 400 µg/mL Liberase DL (Sigma #5466202001) in RPMI (Gibco #72400120). A partial dissociation was performed on a gentleMACS Dissociator (Miltenyi #130-093-235) with the 'm_lung_01' program. The sample was then incubated at 37°C on a nutator set to gentle agitation for 30 minutes, followed by a final complete dissociation step on the gentleMACS using the 'm_lung_02' program. The resulting homogenate was washed with cold 5% FBS in PBS, spun down at 300g, 4°C for 5 minutes, resuspended in cold 5% FBS in PBS, and subsequently filtered through a 70 µm strainer (Fisherbrand #22363548). The suspension was centrifuged and resuspended in FACS buffer (2% FBS in PBS). Concentration was determined by manual cell counting on a hemocytometer and adjusted to 10^6 cells/mL. A portion of the sample was aliquoted for 10x processing with the remaining cells stained with Sytox blue (ThermoFisher #S34857) prior to FACS-sorting for SS2 processing. Note, L3's extracted lung tissue was first preserved at 4°C for 1 day in HypoThermosol FRS solution (Biolife Solutions #101102) before continuing on with the above delineated steps.

*Trachea*

The dissected trachea was rinsed with cold PBS. The tissue was then cut longitudinally with a sterile scalpel to expose the epithelium and placed in 2 mL of digestion buffer containing 5 U/mL dispase (Gibco #17105041), 40 U/mL collagenase I (Gibco #17018029), 2% FBS, and 1x penicillin-streptomycin (Gibco #15140-122) in HBSS (Gibco #14175095). Tissue was minced with sterile dissecting tools in the digestion buffer and incubated at 37°C for 60 minutes, with gentle pipetting performed at regular intervals to assist dissociation. The resulting cell suspension was filtered through a 40 µm strainer (BD #08-771-1), pelleted at 21°C, 2000g for 5 mins and then washed twice with cold PBS. Subsequently, the pellet was resuspended in ACK lysis buffer at 21°C for 1 min and spun down at 21°C, 2000g for 5 minutes. Following two more wash steps in cold PBS with pelleting at 4°C, 2000g for 5 mins, the final pellet was resuspended in FACS buffer containing 2% FBS and 1x penicillin-streptomyocin in cold PBS. Cells were filtered again through a 40 µm strainer. Concentration was determined by manual cell counting on a hemocytometer and adjusted to 10^6 cells/mL for 10x processing. Note, L3's extracted tracheal tissue was first preserved at 4°C for 1 day in HypoThermosol FRS solution (Biolife Solutions #101102) before continuing on with the above delineated steps.

### GASTROINTESTINAL SYSTEM

*Tongue*

The dissociation protocol used for L2 was the same as that in Tabula Muris (Tabula Muris Consortium, 2020; Tabula Muris Consortium et al., 2018). The protocol was then improved for L4 to also capture mesenchymal tissues underneath the epithelium (stromal, muscle tissue and vessels were omitted in the Tabula Muris). After extraction, tongue tissue from L4 was immediately placed on ice. The tongue was further dissected to separate muscles and arteries. The remaining epithelium and underlying mesenchyme were minced with a razor blade, then digested at 37°C on an orbital shaker for 60 minutes in base media, M199-HEPES (ThermoFisher #12340030) supplemented with 2500 U/mL collagenase type IV (Worthington #LS004188) and 100 U/mL DNase I (Worthington #LS006343). A total of two sequential digestion steps were performed (using fresh digestion media each time). Cells were then filtered through a 70 µm strainer (Falcon #352340), pelleted (500g, 5°C, 5 min), and resuspended in PBS containing 1x penicillin-streptomycin (ThermoFisher #15140122), 1x pluronic F-68 (ThermoFisher #24040032), and 2% FBS (Atlanta Biologicals #S11550H). Concentration was determined by manual cell counting on a hemocytometer and adjusted to 10^6 cells/mL. A portion of the sample was aliquoted for 10x processing with the remaining cells stained with Sytox blue (ThermoFisher #S34857) prior to FACS-sorting for SS2 processing.

*Small intestine*

After extraction, small bowel tissue was washed with cold PBS in a 50 mL sterile conical tube (Falcon), then placed on a petri dish and minced with a razor blade. The sample was subsequently incubated at 37°C for 120 minutes in digestion buffer containing the following: 200 U/mL collagenase III (Worthington #LS004183), 100 U/mL DNase I (Worthington #LS002139), 1x antibiotic-antimycotic (ThermoFisher #15240062) dissolved in advanced DMEM/F12 (ThermoFisher #12634028) with 10 mM HEPES (ThermoFisher #15630080) and 1 mM sodium pyruvate (Lonza Switzerland #BW13-115E). Pipetting was performed every 15 min to facilitate complete dissociation. Cells were spun down at 1500 rpm, 4°C, for 5 min, decanted, and resuspended in FACS buffer (2% FBS in PBS) containing 100 U/mL DNase I. The sample was filtered through a 70 µm strainer (Falcon #352350). Concentration was determined by manual cell counting on a hemocytometer and adjusted to 10^6 cells/mL. A portion of the sample was aliquoted for 10x processing with the remaining cells stained with Sytox blue (ThermoFisher #S34857) prior to FACS-sorting for SS2 processing.

*Colon*

After extraction, colonic tissue was washed sequentially in PBS, then in PBS containing 5 mM EDTA. The tissue was then placed in fresh 5 mM EDTA/PBS and incubated at 37°C for 5-30 minutes with intermittent shaking (until complete dissociation of the crypt regions). The sample was spun down at 4°C, 1000 rpm for 2 min, decanted, and resuspended in serum-free media consisting of: advanced DMEM/F12 (ThermoFisher #12634028) with 1x antibiotic-antimycotic (ThermoFisher #15240062), 10 mM HEPES (ThermoFisher #15630080), and 1 mM sodium pyruvate (Lonza Switzerland #BW13-115E). This was followed by incubation at 37°C for 60 min, with pipetting performed every 15 minutes to ensure complete dissociation. Subsequently, the sample was spun down at 4°C, 1500 rpm for 5 min, decanted, and resuspended in digestion solution: 2% FBS in PBS with 100 U/mL DNase I (Worthington #LS002139). After filtering the cells through a 70 µm strainer (Falcon #352350). Concentration was determined by manual cell counting on a hemocytometer and adjusted to 10^6 cells/mL. A portion of the sample was aliquoted for 10x processing with the remaining cells stained with Sytox blue (ThermoFisher #S34857) prior to FACS-sorting for SS2 processing.

*Liver*

The dissected liver (~3 $cm^3$) was immediately placed on ice in a sterile 50 mL conical tube (Falcon) containing cold PBS. Tissue was transferred to a sterile 10 cm polystyrene culture dish (ThermoFisher #150350) containing perfusion buffer (Life Technology #17701-038) prewarmed to 37°C. Two 27G olive-tip regular bevel needles (BD #305109) were inserted into the tissue and a peristaltic pump (Watson-Marlow 120U) used to perfuse the liver for 15 minutes at 7 ml/min. The liver was then moved to a new 10 cm dish containing liver digestion media (Life Technology #17701-034) prewarmed to 37°C, and further perfused with this prewarmed media for 20 minutes at 10 ml/min. The digested media was collected in conical tubes and the softened liver tissue subsequently placed within the media, followed by incubation at 37°C for 20 min with gentle pipetting every 5 min to enhance dissociation. After additional trituration, the single cell suspension was filtered through a 100 µm nylon mesh strainer (BD #352360). The eluent was centrifuged for 5 min, 500g, and resuspended in 2% FBS in PBS. Concentration was determined by manual cell counting on a hemocytometer and adjusted to 10^6 cells/mL. A portion of the sample was aliquoted for 10x processing with the remaining cells stained with Sytox blue (ThermoFisher #S34857) prior to FACS-sorting for SS2 processing. Note, L3's extracted liver tissue was first preserved at 4°C for 1 day in HypoThermosol FRS solution (Biolife Solutions #101102) before continuing on with the above delineated steps.

*Pancreas*

Following necropsy, pancreatic tissue was placed in digestion buffer consisting of 2 mg/mL collagenase VIII (Sigma #C2139) and 0.2 mg/mL trypsin inhibitor (Sigma #T6522) in PBS. Tissue was inflated within the digestion solution using a 27g needle and then incubated in a shaker at 37°C for 8-15 minutes. Following collapse of the tissue, 5 mL of stock FACS buffer (2% FBS in PBS) was added to inactivate collagenase and the cells spun down (all centrifugation steps performed for 5 min at 300 g, 4°C). The cell pellet was resuspended in FACS buffer and filtered through a 100 µm cell strainer (BD #352360). Filtered cells were washed 1x with FACS buffer and subsequently resuspended in 3-5 mL 1x red blood cell lysis buffer (Ebioscience #00-4300) to remove erythrocytes and incubated for 8-10 min at RT. After spinning down, cells were resuspended and washed twice using FACS buffer. After the final wash, cells were resuspended in FACS buffer containing 10 µg/mL DNase I (Roche #4716728001). Concentration was determined by manual cell counting on a hemocytometer and adjusted to 10^6 cells/mL. A portion of the sample was aliquoted for 10x processing with the remaining cells stained with Sytox blue (ThermoFisher #S34857) prior to FACS-sorting for SS2 processing. Note that the pancreas is surrounded by lymph nodes that are challenging to discriminate during dissection. Given that L4's 10x and SS2 pancreas datasets contain a relatively large proportion of immune cells, it is possible that a neighboring lymph node was sampled and processed with the pancreatic tissue.

**URINARY SYSTEM**

*Kidney*

After extraction, renal tissue was first mechanically homogenized using a gentleMACS dissociator (Miltenyi #130-093-235) and then digested under gentle agitation at 37°C for 30 minutes in RPMI buffer containing 10 U liberase TM enzyme (Roche #5401119001), 2% FBS, and 1x antibiotic-antimycotic (Gibco #15240-062). Following trituration with a 5 mL serological pipette, cells were filtered sequentially through 100 µm, 70 µm, and 40 µm strainers (Falcon #352340 #352350 #352360, respectively) with a syringe plunger. Cells were spun down at 4°C, 400g for 10 minutes, then treated with ACK (Gibco #A10492-01) for 5 min at 21°C, and washed with 2% FBS and 1x antibiotic-antimycotic in RPMI. The final pellet was resuspended in 2% FBS in PBS and filtered into 35 µm polystyrene tubes (Falcon #352235). Concentration was determined by manual cell counting on a hemocytometer and adjusted to 10^6 cells/mL. A portion of the sample was aliquoted for 10x processing with the remaining cells stained with Sytox blue (ThermoFisher #S34857) prior to FACS-sorting for SS2 processing. Note, the protocol above was used for L2 and L4. L3's extracted kidney tissue was first preserved at 4°C for 1 day in HypoThermosol FRS solution (Biolife Solutions #101102). The dissociation protocol used for L3 was the same as that used in Tabula Muris (Tabula Muris Consortium, 2020; Tabula Muris Consortium et al., 2018).

*Bladder*

Bladder tissue was placed directly on ice after extraction, then minced. The sample was digested at 37°C on an orbital shaker for 1 hour in Medium 199 (Gibco #11150059) containing 1875 U/mL collagenase type IV (Worthington #LS004188) and 25 U/mL DNase I (Worthington #LS006343), followed by another digestion for 30 minutes in 1x trypLE (ThermoFisher #A1217701) with 25 U/mL DNase I. Cells were subsequently filtered through a 40 µm strainer (Falcon #352340), pelleted (500g, 4°C, 5 min), and resuspended in FACS buffer in PBS pH 7.4 (ThermoFisher #100100-23) containing 1x penicillin-streptomycin (ThermoFisher #15140122), 1x pluronic F-68 (ThermoFisher #24040032), and 2% FBS (Atlanta Biologicals #S11550H). Concentration was determined by manual cell counting on a hemocytometer and adjusted to 10^6 cells/mL. A portion of the sample was aliquoted for 10x processing with the remaining cells stained with Sytox blue (ThermoFisher #S34857) prior to FACS-sorting for SS2 processing.

**REPRODUCTIVE SYSTEM**

*Uterus*

L3's uterine tissue was placed at 4°C in HypoThermosol FRS solution (Biolife solutions #101102) for 2 days. Subsequently, tissue was washed in 1x HBSS (ThermoFisher #14175) and then minced with razor blades. The sample was digested at 37°C for 20 min in 10 U/mL liberase TM (Sigma #5401119001) in RPMI media (Thermofisher #11875119). The sample was then triturated sequentially through 100 µM, 70 µM, and 40 µM strainers (Falcon #352360, #352350, #352340, respectively). Cells were washed, spun down, and resuspended in 2% FBS in PBS. Concentration was determined by manual cell counting on a hemocytometer and adjusted to 10^6 cells/mL for 10x processing.

*Testes*

Immediately after extraction, the testes were detunicated, gently dissociated in PBS, and then incubated at 32°C for 10 min in PBS containing 1 mg/mL type I collagenase (Worthington, #LS004196). Cells were then pelleted via centrifugation for 5 minutes at 250g and the supernatant removed. Another round of digestion with collagenase I was employed, followed by further digestion in trypLE express enzyme (ThermoFisher #12605010) at 32°C for 15 min. During these digestion steps, the seminiferous tubules were mechanically fragmented by vigorous pipetting every 5 minutes. The sample was filtered sequentially through 70 µm and 40 µm strainers (Falcon #352350, #352340, respectively), and then resuspended in cold FACS buffer (2% FBS, 1mM EDTA in PBS). Concentration was determined by manual cell counting on a hemocytometer and adjusted to 10^6 cells/mL. A portion of the sample was aliquoted for 10x processing with the remaining cells stained with Sytox blue (ThermoFisher #S34857) prior to FACS-sorting for SS2 processing.

### References

Casey, K. M., Karanewsky, C. J., Pendleton, J. L., Krasnow, M. R., & Albertelli, M. A. (2021). Fibrous Osteodystrophy, Chronic Renal Disease, and Uterine Adenocarcinoma in Aged Gray Mouse Lemurs (Microcebus murinus). *Comparative Medicine*, *71*(3), 256–266.

Darmanis, S., Sloan, S. A., Zhang, Y., Enge, M., Caneda, C., Shuer, L. M., Hayden Gephart, M. G., Barres, B. A., & Quake, S. R. (2015). A survey of human brain transcriptome diversity at the single cell level. *Proceedings of the National Academy of Sciences of the United States of America*. https://doi.org/10.1073/pnas.1507125112

Der Goukassian, P. (1983). Hematology of the lesser mouse lemur (Microcebus murinus). A preliminary study. *Folia Primatologica; International Journal of Primatology*, *41*(1-2), 129–136.

Fernandez-Godino, R., Garland, D. L., & Pierce, E. A. (2016). Isolation, culture and characterization of primary mouse RPE cells. *Nature Protocols*, *11*(7), 1206–1218.

Ramsköld, D., Luo, S., Wang, Y.-C., Li, R., Deng, Q., Faridani, O. R., Daniels, G. A., Khrebtukova, I., Loring, J. F., Laurent, L. C., Schroth, G. P., & Sandberg, R. (2012). Full-length mRNA-Seq from single-cell levels of RNA and individual circulating tumor cells. *Nature Biotechnology*, *30*(8), 777–782.

Tabula Muris Consortium. (2020). A single-cell transcriptomic atlas characterizes ageing tissues in the mouse. *Nature*, *583*(7817), 590–595.

Tabula Muris Consortium, Overall coordination, Logistical coordination, Organ collection and processing, Library preparation and sequencing, Computational data analysis, Cell type annotation, Writing group, Supplemental text writing group, & Principal investigators.

(2018). Single-cell transcriptomics of 20 mouse organs creates a Tabula Muris. *Nature*, *562*(7727), 367–372.

Travaglini, K. J., Nabhan, A. N., Penland, L., Sinha, R., Gillich, A., Sit, R. V., Chang, S., Conley, S. D., Mori, Y., Seita, J., Berry, G. J., Shrager, J. B., Metzger, R. J., Kuo, C. S., Neff, N., Weissman, I. L., Quake, S. R., & Krasnow, M. A. (2020). A molecular cell atlas of the human lung from single-cell RNA sequencing. *Nature*, *587*(7835), 619–625.

Wu, A. R., Neff, N. F., Kalisky, T., Dalerba, P., Treutlein, B., Rothenberg, M. E., Mburu, F. M., Mantalas, G. L., Sim, S., Clarke, M. F., & Quake, S. R. (2014). Quantitative assessment of single-cell RNA-sequencing methods. *Nature Methods*, *11*(1), 41–46.
